# How a disordered linker in the Polycomb protein Polyhomeotic tunes phase separation and oligomerization

**DOI:** 10.1101/2023.10.26.564264

**Authors:** Tim M. Gemeinhardt, Roshan M. Regy, Tien M. Phan, Nanu Pal, Jyoti Sharma, Olga Senkovich, Andrea J. Mendiola, Heather J. Ledterman, Amy Henrickson, Daniel Lopes, Utkarsh Kapoor, Ashish Bihani, Djamouna Sihou, Young C. Kim, David Jeruzalmi, Borries Demeler, Chongwoo A. Kim, Jeetain Mittal, Nicole J. Francis

**Author notes:** Co-first authors (Tim M. Gemeinhardt, Roshan M. Regy, Tien M. Phan). Correspondence: Jeetain Mittal –, Nicole J. Francis –.

## Abstract

Biomolecular condensates are increasingly appreciated for their function in organizing and regulating biochemical processes in cells, including chromatin function. Condensate formation and properties are encoded in protein sequence but the mechanisms linking sequence to macroscale properties are incompletely understood. Cross species comparisons can reveal mechanisms either because they identify conserved functions or because they point to important differences. Here we use *in vitro* reconstitution and molecular dynamics simulations to compare *Drosophila* and human sequences that regulate condensate formation driven by the sterile alpha motif (SAM) oligomerization domain in the Polyhomeotic (Ph) subunit of the chromatin regulatory complex PRC1. We discover evolutionarily diverged contacts between the conserved SAM and the disordered linker that connects it to the rest of Ph. Linker-SAM interactions increase oligomerization and regulate formation and properties of reconstituted condensates. Oligomerization affects condensate dynamics but, in most cases, has little effect on their formation. Linker-SAM interactions also affect condensate formation in *Drosophila* and human cells, and growth in *Drosophila* imaginal discs. Our data show how evolutionary sequence changes in linkers connecting conserved structured domains can alter condensate properties.

**In brief:** Linking sequence to macroscale properties of biomolecular condensates remains elusive. The authors dissect the function of a disordered linker connecting conserved structured domains in a Polycomb protein with biochemistry and molecular dynamics simulations revealing how sequence changes between *Drosophila* and humans alter condensates and growth in cells and developing flies.

Graphical abstract

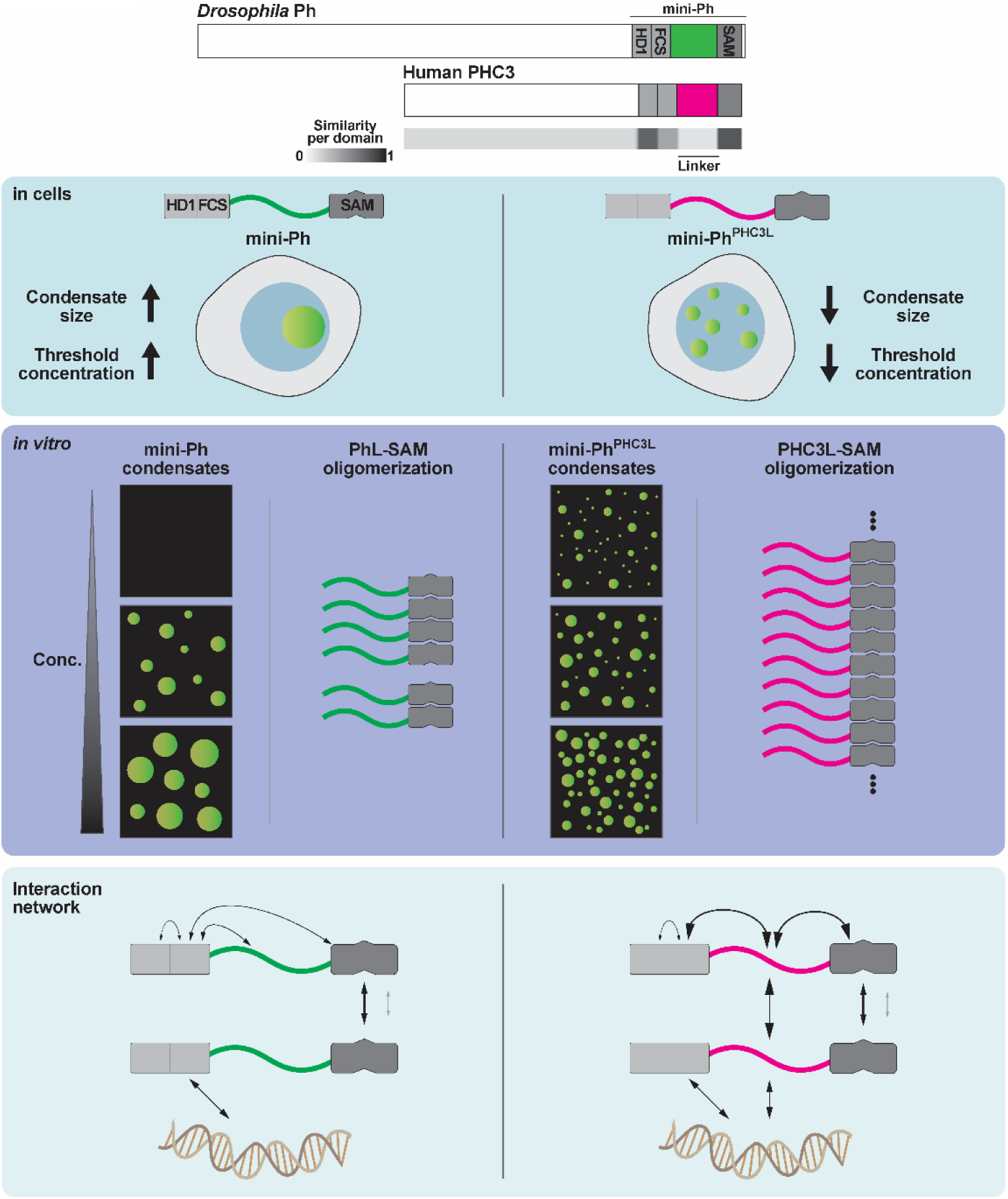

**Highlights:**

- PRC1 condensates form partly through the conserved SAM domain of Polyhomeotic (Ph)
- Linker connecting Ph domains regulates SAM oligomerization and phase separation
- Linker-SAM contacts with human but not fly linker in molecular dynamics simulations
- Linker-SAM contacts tune condensates in *vitro* and in cells, and affect cell growth

## Introduction

The formation of biomolecular condensates and their emergent properties are increasingly recognized as central to the functional organization of molecules in cells^1,2^. This includes chromatin and nuclear proteins that regulate it^3–6^. Condensates form through multivalent interactions that can involve intrinsically disordered regions (IDRs), oligomerization domains, protein interaction domains, nucleic acid binding domains, nucleic acids, or combinations of these^7–10^. Despite intensive investigations, in most cases we still do not know the precise function of condensates in cells^11–13^ in part because the molecular interactions underlying their formation and their biochemical and biophysical properties are not fully known.

The Polycomb system plays a unique and essential role in development by mediating transcription memory—heritable patterns of gene repression triggered by transient developmental cues^14,15^. PcG proteins assemble into complexes that act on chromatin. The two major PcG complexes, PRC1 and PRC2, catalyze the deposition of histone modifications, H2AK119ub and H3K27me3, respectively^16,17^. PcG proteins also regulate chromatin architecture, from local compaction to the formation of large-scale domains^18^. An early observation was that PcG proteins form condensates (originally described as “PcG bodies”) in nuclei^19–22^, and some of the earliest models for PcG function posited that PcG might regulate large chromatin domains by forming a specialized nuclear compartment^23^. More recent studies implicate condensates in epigenetic repression by the Polycomb group (PcG) proteins, including the finding that repressed genes colocalize with condensates^21,22,24–26^. But exactly how PcG condensates form, how they are regulated, how they relate to PcG functions in histone modification and chromatin architecture, and ultimately how they contribute to heritable gene regulation are not understood. Phase separation and related processes^27^ provide a potential mechanistic explanation for PcG condensates and a path to dissecting their function^5,28–34^.

While both PRC1 and PRC2 localize to PcG condensates, PRC1 is more implicated in their formation and in both long– and short-range chromatin organization^19,21,22,25,35–38^. PRC1 (canonical PRC1) consists of 4 subunits in *Drosophila* (human homologs in brackets): Ph (PHC), Pc (CBX), Psc (PCGF), and dRING (RNF2)^17^. Of these four, Pc and Ph are most implicated in regulating chromatin architecture^22,25,26,38,39^.

Within mammalian PRC1, CBX proteins have been shown to drive condensate formation through phase separation with chromatin^28,30,33,40^. CBX2 undergoes phase separation *in vitro*^28,30,41^, and forms condensates *in vivo*^28,30,34^. Phase separation by CBX2 depends on a charged IDR that is also implicated in nucleosome-level chromatin compaction *in vitro*^28,30,34,42^, and chromatin architecture and gene regulation *in vivo*^43^. PRC1 formed with a different CBX protein, CBX8, but without a PHC protein forms condensates with chromatin that also depend on a charged IDR^33^. Ph (PHC) and Pc (CBX) activities must be coordinated in PRC1, and recent work indicates prominent roles for both CBXs and PHCs in PRC1^39,41^. In *Drosophila* embryos, Ph condensates form in embryos lacking Pc, although they are less prominent, while Pc condensates do not form in embryos lacking Ph, implicating both proteins, but implying Ph can form condensates at least partially independent of Pc^18,25^.

Phase separation and regulation of chromatin architecture by Ph/PHCs depends on a conserved oligomerization domain called the Sterile Alpha Motif (SAM) that has head-to-tail polymerization activity^24,44–46^. Disrupting this activity with point mutations disrupts PcG protein clustering, long range chromatin contacts, gene regulation and growth control in developing *Drosophila*^21,22^. However, experiments in *Drosophila* indicate that the SAM has essential functions not fully explained by polymerization: disrupting *ph* in embryos causes patterning defects and lethality; Ph lacking the SAM cannot rescue any of these functions, while Ph with the polymerization interface mutated rescues some but not all functions^24^.

Ph is a large protein (1589 amino acids) (**Figure 1A**). In mammals, Ph has three homologues, PHC1-3. The basic architecture of Ph proteins is conserved and includes three domains: HD1 that is required for assembly into PRC1^47^, an FCS Zn finger that can bind DNA and RNA^48,49^, and the SAM. A long unstructured linker connects the SAM to the FCS, and a large, complex N-terminal IDR comprises more than half of the protein (**Figure 1A, B**). PHC2 also has a prominent short isoform (PHC2_short_) consisting only of the three structured domains and the connecting linker (**Figure 1A**). PHC2_short_ forms condensates in cells and *in vitro* as part of PRC1^21,39^. *Drosophila* Ph lacks a short isoform but we previously showed that the equivalent region (termed “mini-Ph”, **Figure 1A**) can undergo phase separation with DNA or chromatin *in vitro*^31^. Unexpectedly, phase separation by mini-Ph *in vitro* requires the SAM but not its polymerization activity, and Ph with a polymerization mutation can form condensates in cells (unlike Ph lacking the SAM)^31^. Phase separation may thus be the polymerization-independent function of the SAM in development^24^. These data also indicate that the SAM must have interactions outside of its oligomerization interface that are important for phase separation but these interactions have not been identified yet (**Figure 1B**).

**Figure 1:**
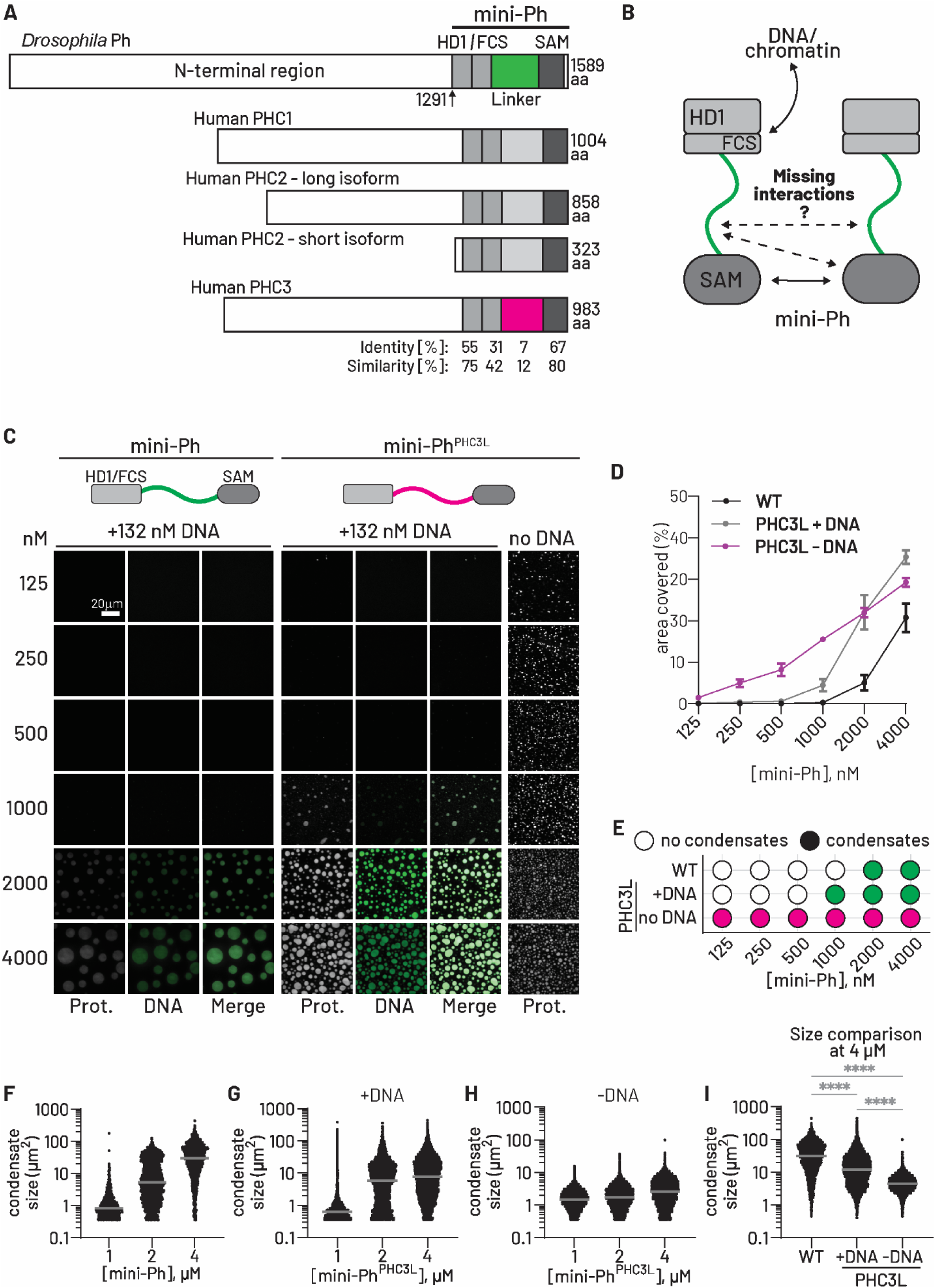
Effect of linker sequence on phase separation of mini-Ph *in vitro*. A. Schematics of *Drosophila melanogaster* Ph and human homologs (PHCs). Identity/Similarity refer to comparison between Ph and PHC3. B. Model of known interactions in mini-Ph underlying phase separation. The HD1 mediates assembly into PRC1. C. Phase separation assays of mini-Ph or mini-Ph^PHC3L^ titrations after overnight incubation. Proteins were labelled with Cy3 and DNA (132 nM, 156 bp dsDNA) was visualized with YOYO1. Reaction conditions: 20 mM HEPES pH 7.9, 60 mM KCl,17 µM ZnCl_2_, 4% glycerol. D. Quantification of area covered by condensate from three phase separation assays as in C. Points show mean and S.D. E. Summary of titrations based on quantification in D. F-H. Quantification of condensate sizes from three experiments. Grey bars indicate median. I. Comparison between condensate sizes of mini-Ph variants at 4 µM. ****: p<0.0001. Data for condensate size comparison were filtered to remove the smallest structures that include artifacts (see Methods for details).

In *Drosophila* cells, mini-Ph forms a single large condensate, and only when expressed at high levels^31^, while PHC2_short_ forms multiple condensates in mammalian cells^21,39^. We wondered what the key differences between mammalian and fly proteins are that direct distinct condensates. In the mini-Ph region, the linker is the least conserved between fly and human proteins (**Figure 1A, B**). The sequence and properties of the three PHC linkers are similar to each other but highly diverged from fly linkers^46^. The human versus fly linker was previously shown to regulate SAM polymerization differently^45^. While Ph SAM alone forms open-ended polymers *in vitro*, attaching the Ph linker (PhL) restricts polymerization^45^. In contrast, the human PHC3 linker (PHC3L) promotes open ended polymerization^45^. The divergent effects of the linkers are independent of which SAM is used (human or fly)^45^. We reasoned that comparing how fly and human linkers function in mini-Ph could reveal conserved and distinct mechanisms underlying phase separation of fly and human Ph proteins, including the role of SAM oligomerization. We chose the linker from PHC3 because it is the best characterized of the three.

Here, we swapped PhL with PHC3L in *Drosophila* (mini-)Ph and found that PHC3L increases phase separation propensity of mini-Ph while reducing the size of the condensates it forms. Using molecular dynamics (MD) simulations, we identified interactions between the PHC3L and SAM and validated their role in phase separation and oligomerization by engineering the SAM domain to form charge-based contacts with PhL. Introducing PhL-SAM contacts promotes SAM oligomerization and mini-Ph phase separation, similar to the effect of the PHC3L. Mutation of the SAM polymerization interface revealed that linker-SAM interactions influence condensate formation largely independent of polymerization, but SAM polymerization impacts condensate dynamics. MD simulations of mini-Ph uncovered unexpected interactions between the HD1/FCS region and both linker and SAM, with distinct patterns depending on whether PhL or PHC3L is present. Together, our data suggest that coupling between the HD1/FCS region and the SAM domain, primarily through linker contacts, plays an important role in condensate formation. In both *Drosophila* and human cells, different linkers and linker-SAM interactions alter condensate formation thresholds and size regulation, corroborating our *in vitro* results. Notably, PHC3L slows cellular growth in *Drosophila* and both PHC3L and engineered linker-SAM interactions suppress imaginal disc development. These findings highlight a critical regulatory role for linker-SAM interactions in phase separation and suggest that condensate dynamics may be directly linked to cellular growth control.

## Results

To understand the role of the linker in phase separation, whether through SAM oligomerization or other mechanisms, we created *Drosophila*-human chimeric proteins by replacing PhL with PHC3L in mini-Ph (mini-Ph^PHC3L^) (**Figure S1**). These chimeric proteins allow us to isolate the effects of the linker from other sequence differences between Ph and PHC3. We tested phase separation of mini-Ph and mini-Ph^PHC3L^ with a 156 bp DNA and found that mini-Ph^PHC3L^ phase separates at approximately half of the concentration required by mini-Ph (**Figures 1C-E, S2A**) and forms smaller condensates (**Figures 1F, G**). Notably, mini-Ph does not phase separate in the absence of DNA^31^. In contrast, mini-Ph^PHC3L^ undergoes phase separation as soon as the salt concentration is lowered, without a requirement for DNA (**Figure 1C**). Tiny condensates are observed at concentrations as low as 125 nM, 4x lower than for mini-Ph^PHC3L^ + DNA (**Figures 1D, E, G, H**). At equivalent protein concentration (4 µM), mini-Ph^PHC3L^ alone forms the smallest condensates, followed by mini-Ph^PHC3L^ + DNA, and finally mini-Ph + DNA (**Figure 1I**). We visualized proteins in condensates by sparse labeling of lysine residues (see Methods), and DNA using the fluorescent dye YOYO-1. To confirm that fluorescent labeling does not affect condensate formation, we formed condensates with no labelled components and visualized them by differential interference contrast (DIC) microscopy (**Figure S2C**). Additionally, we confirmed that condensates are reversible as they are dissolved by increasing [KCl] to 300 mM (**Figure S2D**). These results demonstrate that the PHC3L influences mini-Ph phase separation in three distinct ways: 1) it lowers concentration threshold required for phase separation in the presence of DNA; 2) it enables phase separation without DNA; and 3) it results in the formation of smaller condensates.

### Molecular Dynamics (MD) simulations predict that PHC3L but not PhL contacts the SAM and forms self-interactions

To identify interactions involving linker and SAM that could regulate polymerization, phase separation, or both we carried out MD simulations. We used a recently developed coarse-grained (CG) model, the Hydropathy Scale (HPS) – Urry model^50^ (see Methods), which has been shown to have good agreement with the experimentally observed phase behavior of a diverse set of intrinsically disordered protein (IDP) sequences and multidomain proteins such as HP1α and TDP43^50–53^ CG simulations of a single molecule of PhL or PHC3L attached to the Ph SAM domain were conducted (a snapshot of a linker-SAM construct is shown in **Figure 2A**). To quantify interactions of PhL and PHC3L with SAM, we calculated the average intramolecular contacts formed between the linker and the SAM in these single chain simulations, which are shown as heatmaps (**Figure 2B, C)**.

**Figure 2:**
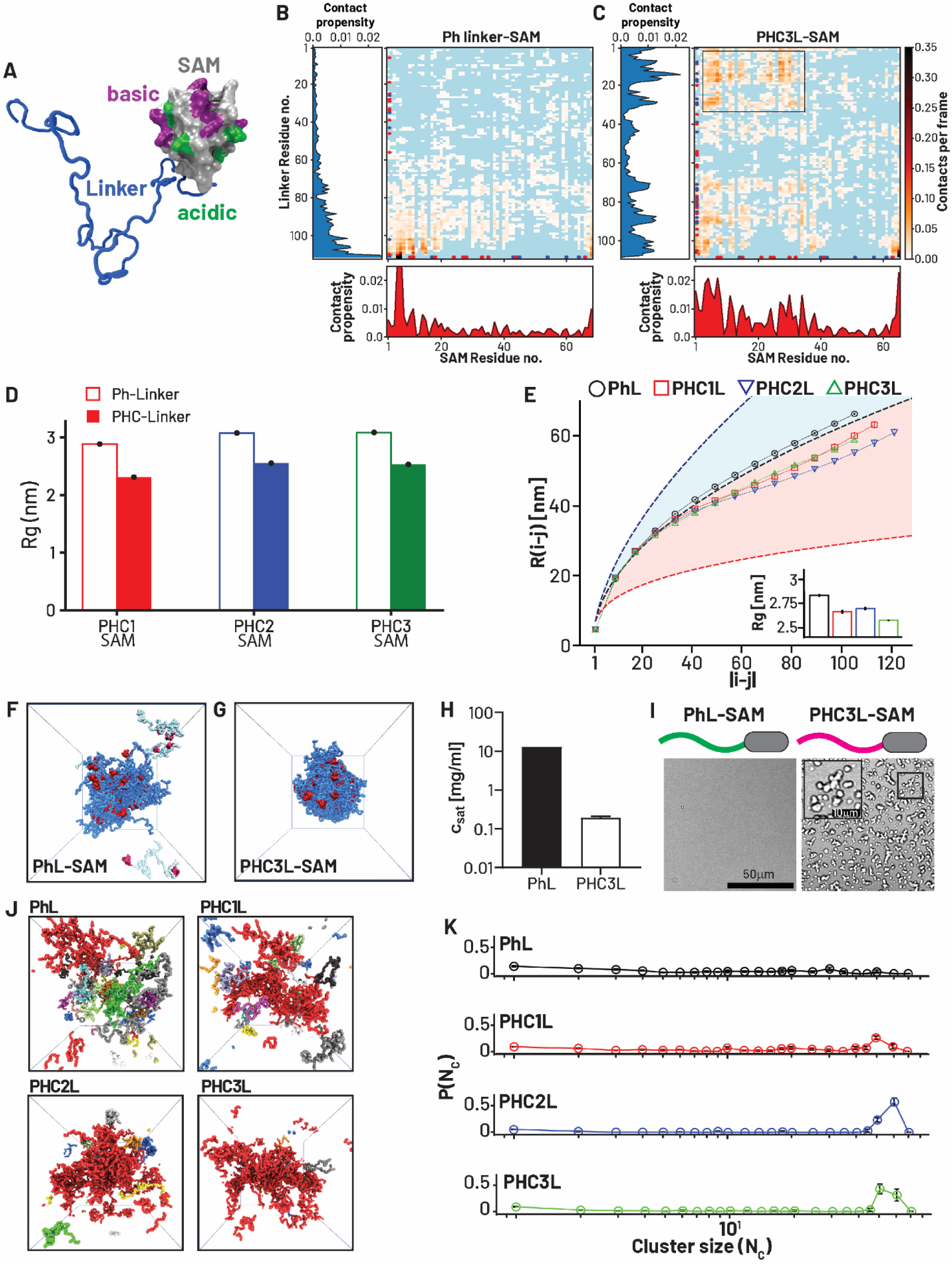
Molecular Dynamics simulations of Ph linker and PHC3L with SAM, and alone. A. Snapshot of linker-SAM model used for simulations. The SAM (PDB 1KW4) is simulated as a rigid body; basic and acidic patches on the SAM are highlighted. B, C. Contact maps from single chain simulations show contact frequencies between linker residues and SAM residues (non-interacting residues colored in light blue) for PhL-SAM (B), and PHC3L-SAM (C). Basic residues (K and R) are highlighted by blue dots and acidic residues (D and E) by red dots. D. Radius of gyration of human PHC SAMs with either Ph linker or corresponding PHC linker from single-chain simulations. Combinations with PHC linker are more compacted reflecting increased intra-chain contacts. E. Intrachain distance distributions (i.e. distribution of distances between ith and jth residue on the linker sequence) and radius of gyration (R_g_) (inset) calculated from single chain simulations show PHC1-3L with higher compaction compared to PhL. Blue dashed line represents the expanded chain limit, black dashed line represents the ideal chain limit and the red dashed line represents the collapsed or globular chain limit. F, G. Snapshots from multichain simulations of PhL-SAM (F) and PHC3L-SAM (G). SAM is red, and linkers are blue. Linkers that are not part of the largest cluster are colored light blue. H. Computed saturation concentration from multi chain simulation is lower for PHC3L-SAM than for PhL-SAM chains I. Phase contrast images of PhL-SAM and PHC3L-SAM upon cleavage of SUMO tag with Ulp (1. 6 µM protein, 50 mM Tris pH 8, 60 mM NaCl, 20 µM ZnCl_2_). J. Snapshots from simulations of Ph and PHC linker chains, where chains belonging to the largest cluster are colored red. K. Quantification of linker clustering from simulations presented in J. Linkers do not phase separate avidly in simulations, but higher clustering is observed for PHCLs compared to PhL.

Comparing both contact maps, we observe that multiple segments of PHC3L interact with the SAM, whereas PhL exhibits only local interactions near its SAM attachment site (**Figure 2B, C**). PHC3L is basic (pI=11) with a high <u>F</u>raction of <u>C</u>harged <u>R</u>esidues (FCR 34%, net charge +5), while PhL is acidic (pI=4) with a lower fraction of charged residues (FCR 15%, net charge –7)^31^.

Simulations reveal that charged residues play a key role in these interactions—specifically, R and K residues in PHC3L frequently interact with D and E residues on the SAM surface (**Figure S2F**). Two prominent acidic patches on the SAM feature conserved acidic residues in tandem, namely Ph residues D1516/D1517 (referred to as “16/17”) and residues D1533/D1534 (referred to as “33/34”) (**Figure S2F**). Strong interactions are observed between these patches and the N-terminal region of the PHC3L. To determine whether linker-SAM interactions are conserved in human proteins, we carried out simulations with each PHCL with its own SAM and found that all native PHC-SAM combinations exhibited strong interactions, while the PHC SAMs showed sparse interactions with PhL (**Figure S2G, H**). We also calculated the radius of gyration (R_g_) of human PHC SAMs paired with either PhL or their corresponding PHCL; smaller R_g_ values with PHC linkers indicate compaction of the chain due to interactions (**Figure 2D**).

In addition to heterotypic interactions between linker and SAM, linkers could form homotypic interactions, which can promote phase separation. We calculated the R_g_ and intrachain distances for each linker sequence (**Figure 2E**). Both metrics provide complementary information on the single chain conformational properties, which have been shown to have a good correlation with phase separation propensity for natural sequences^54,55^. We calculated the average inter-residue distance R_ij_ between the i^th^ and j^th^ residue as a function of residue separation, |i-j|. We then fit the distribution of inter-residue distances to the power law R_ij_ = b|i-j|^ν^ where b is the Kuhn length, which is set to 0.55 for disordered proteins, and ν is the polymer scaling exponent. A ν value of 0.33 or below indicates a collapsed or globular polymer conformation, while a value of 0.5 corresponds to an ideal chain, and a value close to 0.588 reflects an expanded chain. As shown in **Figure 2E**, PhL lies between the expanded and ideal chain regimes, but it is quite close to behaving like an ideal chain. All three PHCLs on the other hand are more collapsed and lie between the ideal chain and the globular chain limit. The R_g_ behaves similarly, in that PhL is the most expanded chain and PHC3L is most collapsed. Contact maps also show increased contact frequencies among PHC3L residues compared to PhL (**Figure S2I**) highlighting the influence of intramolecular interactions on linker conformations.

To determine whether linker-SAM interactions can capture the changes in phase separation observed with mini-Ph with different linkers, we conducted multichain MD simulations of PhL-(d)SAM and PHC3L-(d)SAM. Simulations were conducted at 300 K and 100 mM salt concentration. Exemplary simulation snapshots are shown in **Figure 2F and G**. Densities of the coexisting dense and dilute phases were calculated from these simulations. PHC3L-SAM shows higher phase separation propensity (nearly two orders of magnitude lower c_sat_) than PhL-SAM (**Figure 2H**).

To test the prediction that PHC3L-SAM has a higher propensity for phase separation than PhL-SAM, we produced linker-SAM proteins in *E. coli* using the cleavable SUMO solubility tag. Consistent with previous findings^31^, PhL-SAM does not form condensates when the salt concentration is lowered, either before or after SUMO cleavage by Ulp1. In contrast, PHC3L-SAM forms condensates when the salt concentration is lowered, both before and after Ulp1 cleavage (**Figure 2I, Figure S1B, S2J**). Condensates formed with the intact SUMO tag are large and round, while those formed after cleavage are small and irregularly shaped (compare **Figure S2J and Figure 2I**). Taken together, these results demonstrate that the PHC3L lowers the threshold for phase separation, consistent with predictions from MD simulations.

To identify the direct contribution of linker-linker interactions to phase separation, we also conducted multichain simulations with each of the three PHC linkers and PhL on their own. All linker sequences showed weak tendency to form phase separated droplets, and instead formed few dispersed clusters that varied in size (**Figure 2J)**. Even upon increasing the solution concentration further by reducing the simulation box size, we did not observe the formation of a distinct condensed phase indicating that homotypic linker interactions alone are not sufficient to explain mini-Ph or mini-Ph^PHC3L^ phase separation. Nevertheless, we compared the self-assembly of each linker by calculating the distribution of cluster sizes from these multichain simulations (**Figure 2K**). Single chain collapse and cluster sizes are correlated, with both increasing for human linkers, particularly PHC2 and PHC3 linkers. Collectively, these results suggest that human linkers have higher homotypic interactions than their fly counterpart.

Taken together, MD simulations indicate that linker-SAM and linker-linker contacts differentiate PhL from PHCLs. These contacts, especially linker-SAM contacts, are driven by charged residues, and are predicted to drive the increased phase separation propensity of mini-Ph^PHC3L^ (and PHC3L-SAM) compared to mini-Ph (and PhL-SAM) in agreement with our experimental data (**Figure 1E**). They also likely explain the increase in SAM oligomerization that occurs when the SAM is fused to PHC3L versus PhL^45^.

### Engineered charge complementarity between Ph linker and SAM increases oligomerization and phase separation

The simulations suggest that charge-based interactions between basic residues in PHC3L and acidic residues on the surface of the SAM could explain PHC3L effects on polymerization and phase separation. To test this hypothesis, we designed mutations on the surface of the SAM that could make charge-based interactions with the (acidic) PhL. Specifically, the two pairs of tandem aspartic acid (D) residues that interact with PHC3L were mutated to arginine (R) and lysine (K) (“D16K/D17R” and “D33K/D34R”). This switches small negative patches to positive, to potentially drive interactions with negatively charged residues in PhL. The negative patches are conserved in PHC3 SAM, although the D34 position is an E in PHC3 (**Table S1**). The net charge of the SAM is –4; mutation of a single acidic patch neutralizes the SAM net charge and mutation of both patches inverts it to +4 (the combination of D16K/D17R and D33K/D34R is referred to as SAMsurf) (**Figure 3A**). We first determined the effects of the SAM surface mutations on SAM-SAM interactions using multichain simulations of the SAM. While our CG model did not capture canonical SAM oligomerization, we observed non-canonical clustering of WT SAM chains (**Figure S3A**). Neutralizing the SAM charge by introducing single pairs of mutations increased clustering of SAM chains, which is likely a result of reduced electrostatic repulsion (**Figure S3A**). Importantly, introduction of both pairs of mutations abolished SAM clustering completely (**Figure S3A**). These non-canonical SAM interactions, which depend on acidic surface residues, may contribute to protein assembly and phase separation *in silico* and possibly in experiments.

**Figure 3:**
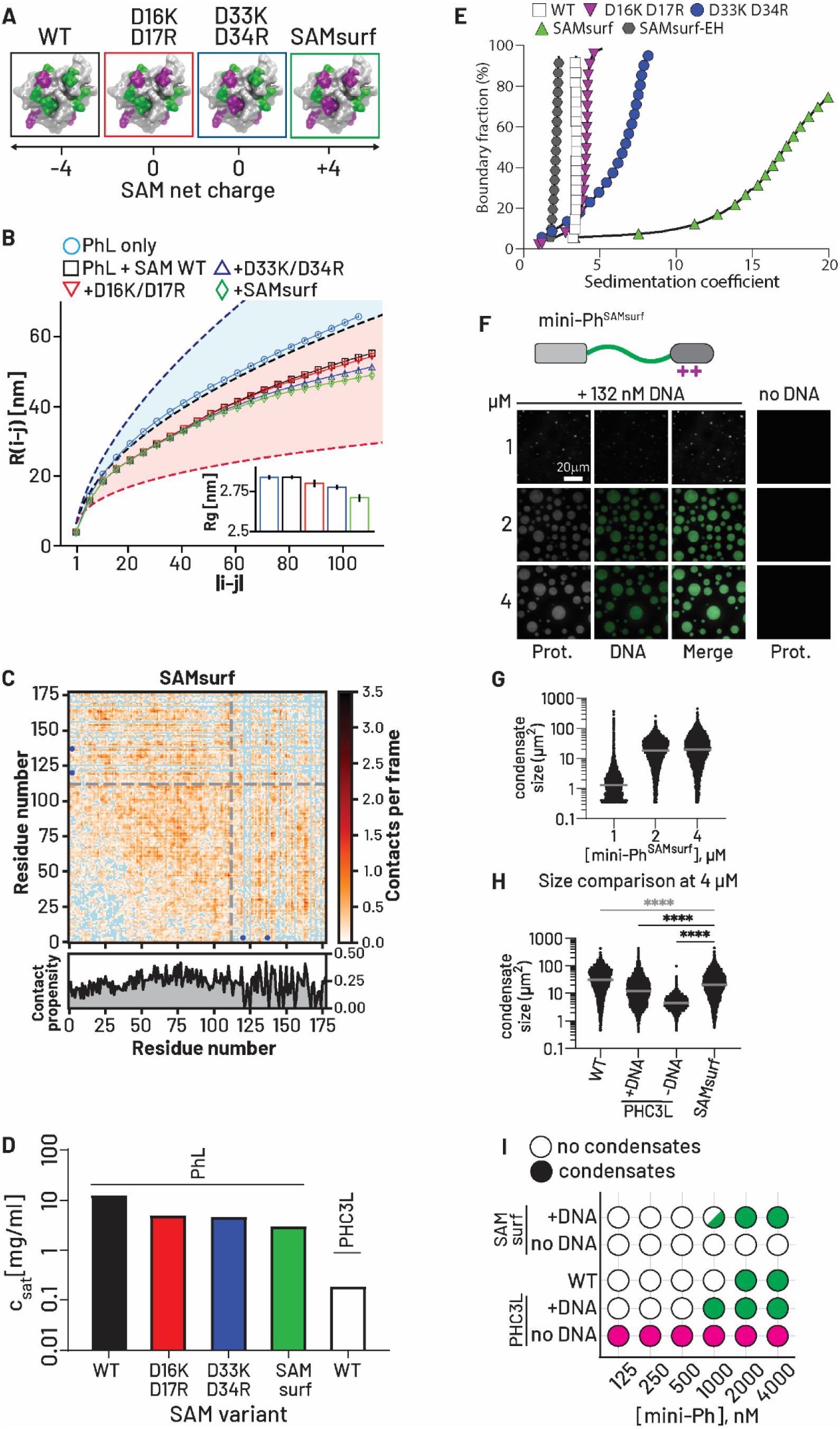
Introducing charge complementarity between PhL and SAM promotes linker-SAM contacts, oligomerization and phase separation. A. SAM mutants were designed to change acidic residues (green) on the SAM surface to basic ones (magenta). B. Intrachain distance distributions (i.e. distribution of distances between ith and jth residue on the linker sequence) and radius of gyration (inset) calculated from single chain simulations show PhL with SAM surface charge mutants have a higher compaction compared to PhL with WT SAM. Blue dashed line represents the expanded chain limit, black dashed line represents the ideal chain limit and the red dashed line represents the collapsed or globular chain limit. C. Intermolecular contact maps from multi chain simulations of PhL with SAMsurf show overall contact frequencies increase compared to WT SAM (Figure 2B) (non-interacting residues colored in blue). Grey lines mark linker-SAM boundary; blue dots indicate mutated sites. D. Computed saturation concentrations from multichain simulation is lowest for PHC3L-WT SAM, intermediate for PhL-SAM surface charge mutants and highest for PhL-WT SAM chains. E. Analytical ultracentrifugation (AUC) of PhL-SAM with different SAM mutations. F. Phase separation assays of mini-Ph^SAMsurf^ after overnight incubation. G. Quantification of condensate areas from three experiments. Grey bars indicate median. H. Comparison between condensate sizes of mini-Ph variants at 4 µM (with data from Figure 1). ****: p<0.0001. I. Summary of titration experiments with comparing mini-Ph^SAMsurf^ (WT and PHC3L data from Figure 1). Data for condensate size comparison were filtered to remove the smallest structures that include artifacts (see Methods for details).

To dissect how the SAM surface mutations affect interactions with linkers, we first conducted single chain simulations. We measured the impact of each SAM mutant on the polymer scaling and R_g_ of PhL. We observed increasing linker collapse as the overall SAM charge increases (**Figure 3B** and inset). We also observed increasing linker-SAM interactions in intermolecular contact maps from multichain simulations, which correlate with increases in the overall SAM charge (**Figure 3C; Figure S3B, C**). These contacts were primarily mediated by acidic residues in PhL, and the designed basic patches on the SAM (**Figure S3B**), suggesting that charge complementarity governs PhL-SAM interactions.

Multichain simulations of PhL with SAM surface mutants predict a small increase in phase separation propensity (lower c_sat_) relative to WT (**Figure 3D, S3C**), although c_sat_ is still higher than for PHC3L-SAM. This gap may be explained by the additional contribution to phase separation through the linker-linker interactions of PHC3L (**Figure 2J, K**), and loss of SAM-SAM clustering in the case of the 4-residue mutant, SAMsurf (**Figure S3A**). Taken together, simulations confirm new PhL-SAM interactions occur upon introducing basic patches to the SAM, without altering the sequence of PhL. To test the importance of linker-SAM contacts for SAM oligomerization, we used Sedimentation Velocity Analytical Ultra Centrifugation (SV-AUC), which was previously used to measure SAM polymerization, including the effects of PhL versus PHC3L^45^. We carried out SV-AUC with WT PhL-SAM, and PhL with each of the three SAM mutants. We used van Holde-Weischet analysis to calculate the corrected sedimentation coefficients and plotted them against their corresponding boundary segments (**Figure 3E**). For WT PhL-SAM, we observed a nearly vertical line on the van Holde-Weischet plot indicating a single species of oligomers. Mutation of either residues D16K/D17R or D33K/D34R results in more sigmoidal shaped curves.

These curves cover a wider range of sedimentation coefficients with a drop in the lower bound as well as an increase in the upper bound indicating the presence of both smaller and larger oligomers compared to the WT. The D33K/D34R variant also has species with higher sedimentation coefficients than the D16K/D17R variant indicating stronger enhancement of oligomerization (**Figure 3E**, blue circles vs. purple triangles). The largest shift in oligomerization was observed after introducing both pairwise mutations (SAMsurf), with an extreme increase in the maximum range of sedimentation coefficients (**Figure 3E**, green triangles). This recapitulates the open-ended oligomerization observed with the SAM alone or attached to PHC3L^45^. Importantly, these effects are strictly dependent on the SAM polymerization interface, because mutating the SAM-SAM interface (end-helix (EH) mutation, L1565R in Ph) in the SAMsurf mutant reverts the protein to low S-values and homogeneous behavior (**Figure 3E**, grey hexagons). Thus, introducing charge complementarity between PhL and the SAM allows extensive oligomerization, similar to the effect of replacing PhL with PHC3L. The increase in oligomerization for variants D16K/D17R and D33K/D34R could be because of the increase in both linker-SAM and SAM-SAM interactions (**Figure S3A**). Because SAM-SAM clustering is not observed for the SAMsurf mutant (**Figure S3A**), increased oligomerization of this protein can be attributed to linker-SAM interactions.

To determine whether linker-SAM interactions can also contribute to phase separation, we prepared mini-Ph with the SAMsurf mutant (**Figure S1A**) and tested it in phase separation assays *in vitro*. Titrations indicate that phase separation of mini-Ph^SAMsurf^ with DNA occurs at concentrations between those for mini-Ph and mini-Ph^PHC3L^ (**Figure 3F-I, S2A**). mini-Ph^SAMsurf^ does not phase separate without DNA at the concentrations we could achieve. This result suggests linker-SAM interactions can enhance phase separation and also indicate that SAM clustering (**Figure S3A**), which is eliminated by the SAMsurf mutations, is not required for phase separation. Because the SAMsurf mutations introduce positive charges, we wondered if they would allow the SAM to bind DNA, which might contribute to phase separation. We measured DNA binding of PhL-SAM, PhL-SAMsurf, and PHC3L-SAM by EMSA. We did not detect DNA binding by PhL-SAM or PhL-SAMsurf, but unexpectedly, PHC3L-SAM binds DNA, an effect which must be mediated by the PHC3L (**Figure S3D**).

### The canonical SAM polymerization interface is not required for regulation of phase separation by the linker or by engineered linker-SAM interactions

The effects of the PHC3L and the SAMsurf mutations on oligomerization (as measured by AUC) are strictly dependent on the canonical SAM-SAM interface because mutating this interface blocks oligomerization (**Figure 3E**). We previously found that phase separation of mini-Ph is not dependent on the polymerization interface^31^. To determine whether the effects of the SAMsurf mutations or the PHC3L on phase separation depend on SAM polymerization activity, we introduced the polymerization-disrupting EH mutation to each of the three mini-Ph variants (**Figure S1A**). We tested these proteins in phase separation assays with DNA. All three proteins form salt-reversible condensates in the presence of DNA (**Figure 4A, S2C-E**). mini-Ph^PHC3L-EH^ phase separates at the lowest concentration, followed by mini-Ph^SAMsurf-EH^ and finally mini-Ph^EH^ (**Figure 4B, S2B**), the same order of concentration dependence observed with the polymerization-competent SAM (**Figure 3I, S2A**). Condensate size at equivalent concentrations also follows the same order as for polymerization competent proteins, with mini-Ph^EH^ forming the largest, and mini-Ph^PHC3L-EH^ the smallest condensates (**Figure 4C**).

**Figure 4:**
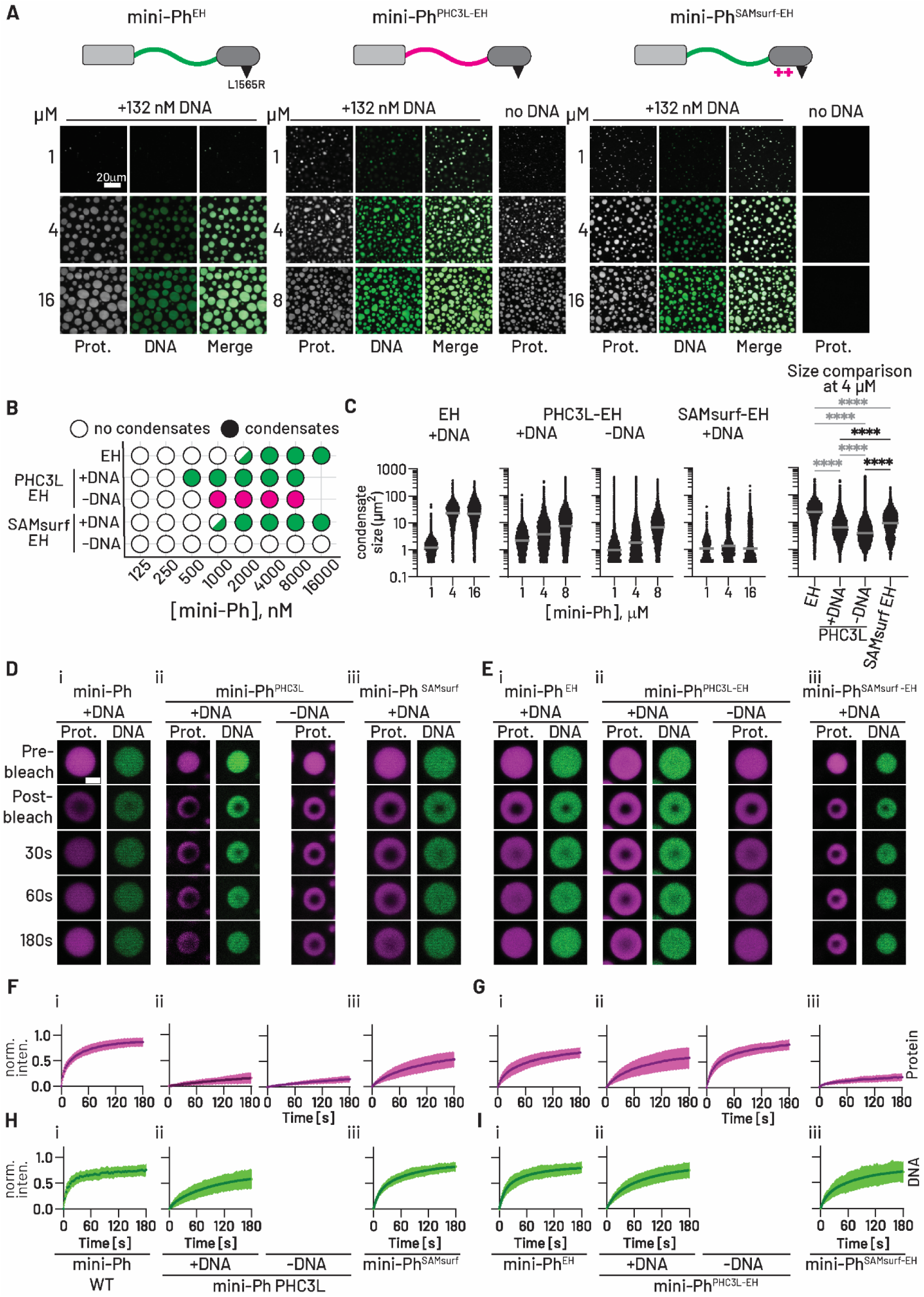
Effect of SAM oligomerization on mini-Ph phase separation and internal condensate dynamics. A. Images from phase separation assays with mini-Ph^EH^, mini-Ph^PHC3L-EH^, or mini-Ph^SAMsurf-EH^ after overnight incubation. B. Summary of titration experiments of SAM oligomerization mutants (EH). C. Quantification of condensate areas from three experiments. Grey bars indicate median. ****: p<0.0001. Data for condensate size analysis were filtered to remove the smallest structures that include artifacts (see Methods for details). D, E. Representative images from FRAP of mini-Ph (D), or mini-Ph^EH^ (E) variants. Proteins (Cy3 labelled) were used at 4 µM and DNA (YOYO1 labelled) at 132 nM. Scale bar: 2 µm. F-I. Summary of FRAP experiments for protein (F, G) and DNA (H, I) for mini-Ph (F, H), and mini-Ph^EH^ (G, I) variants. The curves represent fits of the combined data for each condition (n≥34) with standard deviations indicated by shading. See also **Figure S4.**

We also tested the proteins without DNA. This revealed a striking difference in the behavior of mini-Ph^PHC3L –EH^ (**Figure 4A**). While mini-Ph^PHC3L-EH^ still forms condensates without DNA, it requires approximately 2x higher protein concentration than with DNA (**Figure 4A, B, S2B**). mini-Ph^PHC3L^ with the intact SAM forms condensates without DNA at approximately 4x lower concentrations than with DNA (**Figure 1C, D**). This suggests that in the absence of DNA, SAM oligomerization strongly promotes phase separation and that DNA binding may inhibit SAM oligomerization in the context of mini-Ph^PHC3L^. The DNA binding of PHC3L that we observe (**Figure S3D**) might contribute to this inhibition.

Pre-formed oligomers could promote phase separation, especially in the case of mini-Ph^PHC3L^ without DNA, and might explain differences among proteins. To test this possibility and to confirm the quality of the proteins used in this work, we analyzed the oligomerization state of each protein in storage buffer conditions using size exclusion chromatography coupled to multi-angle light scattering (SEC-MALS) and mass photometry (**Figure S1D-L**). All three proteins with the SAM polymerization interface intact form small oligomers (5, 6, 7-8 for mini-Ph, mini-Ph^PHC3L^, and mini-Ph^SAMsurf^, **Figure S1D, F**). In each case, introducing the polymerization interface mutation (EH) reduces the oligomeric state to a mix of monomers and dimers (**Figure S1E, F**). We then characterized each protein using mass photometry. Mass photometry also shows small oligomers for each protein that decrease in size at lower concentrations (**Figure S4G-L)**. These experiments are not consistent with large pre-formed SAM polymers explaining differences among the different proteins. We conclude that phase separation and its enhancement by PHC3L or SAMsurf in the presence of DNA have little dependence on SAM oligomerization. This is consistent with our previous results with mini-Ph and chromatin^31^, and with the results with mammalian PRC1 and chromatin^39^.

### Linker and SAM polymerization activity both modulate protein and DNA dynamics in condensates

While the canonical SAM polymerization is not required for phase separation, our previous work on mini-Ph and the study by Niekamp *et al.* on mammalian PRC1 demonstrated its influence on condensate component dynamics^31,39^.

To understand how the PHC3L, SAMsurf mutations, and the EH polymerization mutation affect protein and DNA dynamics in condensates, we performed Fluorescence Recovery After Photobleaching (FRAP) assays (**Figure 4D-I**). Most FRAP data were fitted with a double exponential function to determine parameters for both slow and fast recovering populations (**Figure S4, Table S2**). However, mini-Ph^PHC3L^ and mini-Ph^SAMsurf^ proteins, lacking a fast-recovering population, were best fit with a single exponential. In mini-Ph condensates, both protein and DNA exhibited high mobility, with mobile fractions of 92 and 78%, respectively (**Figure 4Di, Fi, Hi, Table S2**). In contrast, mini-Ph^PHC3L^-DNA condensates showed low protein mobility (34% mobile fraction), despite maintaining high DNA mobility (92%) (**Figure 4Dii, Fii, Hii, Table S2**). In the absence of DNA, mini-Ph^PHC3L^ condensates showed even lower protein mobility (18%, **Table S2**). Notably, mini-Ph^PHC3L^, both with and without DNA, exhibited variable mobility patterns, with some condensates showing virtually no protein recovery, as evident when all fit values are plotted (**Figure S4A-D**). Mini-Ph^SAMsurf^ demonstrated intermediate behaviors: reduced protein mobility (64%) and slower recovery compared to mini-Ph, but higher mobility and faster recovery than mini-Ph^PHC3L^, while maintaining high DNA mobility (90%)(**Figure 4Diii, Fiii, Hiii, S4, Table S2**). While mini-Ph^SAMsurf^ protein showed a wide range of mobility and recovery patterns, DNA recovery remained consistent within the same condensates (**Figure S4A-D**). Together with MD simulations, these results suggest that PHC3L and the SAMsurf mutations enhance intermolecular interactions and reduce condensate dynamics, with PHC3L showing a more pronounced effect.

We then tested the same protein series with the EH polymerization mutant (**Figure 4E-H**). Introduction of the EH mutation increases the mobility of mini-Ph^PHC3L^ both with and without DNA (77%, 92% mobile, respectively), while mini-Ph remains mobile (74%) (**Figure 4Ei,ii, Gi,ii, Table S2**). The heterogeneity among condensates persists for mini-Ph^PHC3L-EH^, particularly with DNA (**Figure S4E-H**). Surprisingly, mini-Ph^SAMsurf-EH^ has low mobility in condensates (29% mobile) (**Figure 4Eiii, Giii**). We note that Niekamp *et al.* also found that while condensates formed with polymerization mutants of PHC2 in PRC1 are more dynamic, the morphologies of condensates formed with PRC1 containing PHC1 with the same mutation are consistent with slower dynamics^39^. This is consistent with a balance of interactions determining dynamics. It should also be noted that the EH mutation is L→R; while the wild-type SAM-EH remains net negatively charged (–3), SAMsurf-EH shifts from +4 to +5, which might increase interactions with PhL. DNA is mobile in condensates formed with all three EH mutant proteins, although recovery is slightly slower in mini-Ph^PHC3L-EH^ and mini-Ph^SAMsurf-EH^ than in mini-Ph^EH^ (**Figure 4E, I, Table S2**). Proteins with the polymerization interface intact were prepared in Sf9 cells and carry an N-terminal FLAG tag (DYKDDDDK), while proteins with the EH mutation were prepared in *E. coli* and have a C-terminal 6X-His tag. Because epitope tags have been shown to affect protein dynamics in phase separation assays in some cases^56^, we have not directly compared FRAP parameters from the two sets of proteins. We conclude that although the SAM polymerization interface is not required for condensate formation or the effects of PHC3L and the SAMsurf mutations on it, it has strong effects on protein mobility, especially for the PHC3L. This could indicate that SAM polymers form preferentially in condensates, although this would need to be directly tested in future studies.

### MD simulations of mini-Ph predict additional interactions between HD1/FCS and both linker and SAM

To determine whether the interactions predicted for linker-SAM constructs are likely to occur in the context of mini-Ph, we extended the MD simulations to the entire mini-Ph protein and full-length (FL) Ph (**Figure 5A, B, S5A)**. We plotted the van der Waals (vDW) contact probabilities per residue of the PhL-SAM region in the context of mini-Ph and FL Ph against those for the corresponding linker-SAM and calculated Pearson correlation coefficients (**Figure 5G, S5B**). The Pearson correlation coefficients in mini-Ph and FL Ph are approximately 0.80 and 0.75 respectively, suggesting a strong positive linear correlation in the contact propensity between the simulations of PhL-SAM and larger contexts of mini-Ph and Ph. We then carried out simulations of mini-Ph^PHC3L^ and mini-Ph^SAMsurf^ (**Figure 5C-F**) to examine the correlations between contacts in the PHC3L-SAM and PhL-SAMsurf versus their mini-Ph^PHC3L^ or mini-Ph-^SAMsurf^ contexts. We found strong positive correlations in both cases, with Pearson correlation coefficients of approximately 0.94 between the PHC3L-SAM and mini-Ph^PHC3L^ contexts (**Figure 5G**), and 0.89 between PhL-SAMsurf and mini-Ph^SAMsurf^. Thus, interactions predicted using linker-SAM are likely to be highly relevant within the mini-Ph context.

**Figure 5:**
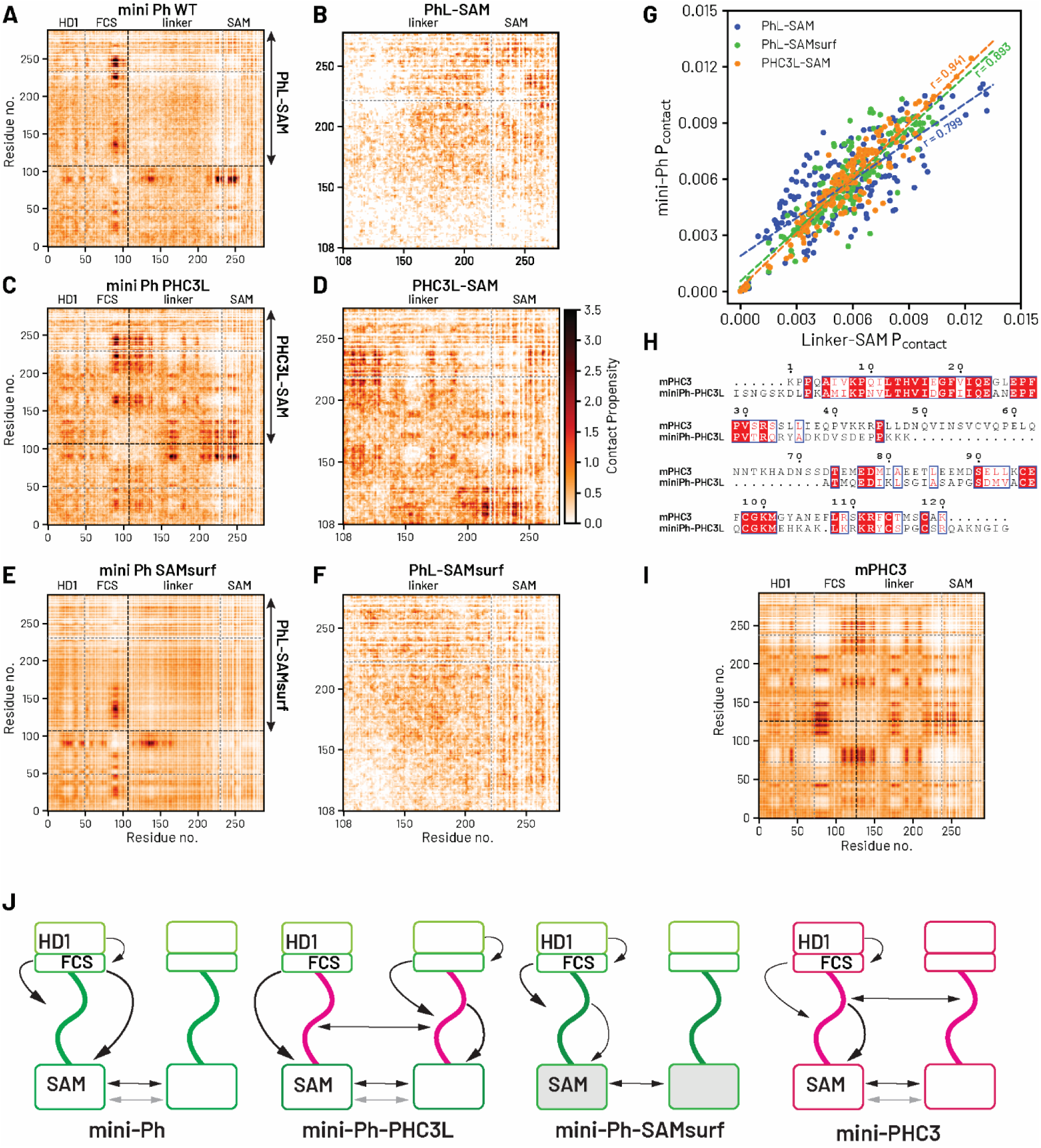
MD simulations predict that linker-SAM interactions are largely preserved in mini-Ph and reveal HD1/FCS contacts. A-F. Intermolecular contact maps of the three mini-Ph constructs (A, C, E) and their corresponding linker-SAM versions (B, D, F) within the dense phase of the CG coexistence phase separation. Light and dark colors present unfavorable and favorable interactions, respectively. G. The correlations between the contact probability per residue formed in the three linker-SAM variants (x-axis) — PhL-SAM, PhL-SAMsurf, and PHC3L-SAM — and their correspondence in the mini-Ph (y-axis) in the CG coexistence phase simulations. H. Multiple sequence alignment between the HD1/FCS region of PHC3 and dPh. Red boxes with white letters show identical amino acids, while white boxes with red letters indicate amino acids with similar properties. I. Intermolecular contact map within the dense phase of mini-PHC3 in the CG coexistence phase separation. J. Schematic summarized the interaction networks predicted by MD simulations.

Simulations of mini-Ph also revealed additional interactions made by the HD1/FCS region, that are distinct in each protein variant (**Figure 5, S5**). In mini-Ph (**Figure 5A, S5C**), MD simulation predicts prominent interactions between FCS and HD1, FCS and PhL, and FCS-SAM. For mini-Ph^PHC3L^ (**Figure 5C**), FCS-HD1 and FCS-SAM (along with PHC3L-PHC3L and PHC3L-SAM) interactions are still present, but the FCS is predicted to make different interactions with the linker (as expected since the linker sequences are different). The FCS is highly basic and is predicted to interact with the SAM via the acidic residues mutated in mini-Ph^SAMsurf^. Indeed, in mini-Ph^SAMsurf^, FCS-SAM interactions are not present (**Figure 5E**). Although experimental validation will be needed in future work, these data indicate that the mini-Ph protein undergoes a complex array of interactions among its domains and with the linker. The changes in contact maps when PHC3L is introduced indicate that the effect of PHC3L is not restricted to linker-SAM and linker-linker, but also affects HD1/FCS interactions. Similarly, loss of FCS-SAM interactions in mini-Ph^SAMsurf^ may counter the addition of linker-SAM interactions, perhaps explaining the relatively subtle effect of these mutations on phase separation (**Figure 3I**). The main interactions predicted for each protein are summarized in **Figure 5J**.

The mini-Ph structured domains are conserved but not identical (**Figure 5H**). We therefore carried out simulations of mini-PHC3 (all sequences derived from PHC3) (**Figure 5I**). The pattern of predicted contacts is distinct from mini-Ph^PHC3L^ in that the FCS does not interact with the SAM, but instead with positively charged regions of PHC3L (summarized in **Figure 5J**). Inspection of the HD1/FCS regions of Ph and PHC3 reveals that there is an additional sequence between the HD1 and FCS in PHC3 that contains several acidic amino acids (**Figure 5H**). Thus, while the main interactions in mini-Ph are HD1-FCS; FCS-linker; FCS-SAM; SAM-SAM, those in mini-PHC3 are HD1-FCS; FCS-linker; linker-linker; linker-SAM; SAM-SAM. Together, the simulations reveal extensive coupling between the domains in mini-Ph, with the linker playing a key role. Coupling is present in both human and *Drosophila* proteins, but with compensatory sequence changes altering the precise nature of the interactions.

### Linker interactions affect condensate formation in cells

Our *in vitro* and *in silico* experiments suggest that the linker-SAM interactions present with PHC3L and SAMsurf promote SAM oligomerization and phase separation and alter condensate size and dynamics. To relate these mechanistic observations to condensate formation in cells, we prepared stable *Drosophila* Kc167 cell lines with copper-inducible expression of Venus tagged mini-Ph, mini-Ph^PHC3L^, or mini-Ph^SAMsurf^. We used live imaging to analyze condensate formation (**Figure 6A**). Previously, we observed that Venus-mini-Ph forms condensates in a small fraction of cells after transient transfections into *Drosophila* S2 cells, and it nearly always forms a single large condensate that excludes chromatin^31^. This pattern also occurs in the Kc167 cell line. In striking contrast, both mini-Ph^PHC3L^ and mini-Ph^SAMsurf^ form several small condensates in nuclei, more similar to FL Ph (compare **Figure 6A** to **6F**). To assess the relationship between protein concentration and condensate formation, we quantified the mean nuclear intensity of Venus signal (as a proxy for Venus-mini-Ph concentration) and stratified cells by the presence of condensates (**Figure 6B**). As expected, cells with condensates exhibit higher mean intensities. While mini-Ph is expressed at higher levels than either mutant, the range of expression levels in cells with condensate for mini-Ph^PHC3L^ or and mini-Ph^SAMsurf^ are clearly below those for mini-Ph and overlap mini-Ph expression levels in cells without condensates. This is consistent with PHC3L and SAMsurf mutations lowering the threshold for condensate formation in cells, mirroring their effects *in vitro* (**Figure 1E, 3I**). We further quantified condensate size and plotted it against mean nuclear intensity (**Figure S6Ai**). As expected, the two parameters are positively correlated (**Figure S6Ai**). To compare condensate sizes in cells with similar expression levels, we selected a window of intensities shared among all three proteins, spanning the highest expressing mini-Ph^PHC3L^ and mini-Ph^SAMsurf^ cells, and the lowest expressing mini-Ph cells with condensates (**Figure S6Aii**). We also compared the numbers of condensates per nucleus for all cells (**Figure 6C**). These data indicate that mini-Ph^PHC3L^ and mini-Ph^SAMsurf^ form small condensates at low concentrations, while mini-Ph forms a large condensate at high expression levels (**Figure 6A-C, S6A**). Ph functions in cells as part of the PRC1 complex. To test whether mini-Ph proteins assemble into PRC1 in cells, and if the PHC3L or SAMsurf mutations affect PRC1 assembly, we carried out co-immunoprecipitation (co-IP) with each cell line with anti-GFP beads (**Figure 6D**). We find that all three proteins co-precipitate with Pc, a core PRC1 component, at equivalent levels (**Figure 6E**).

**Figure 6:**
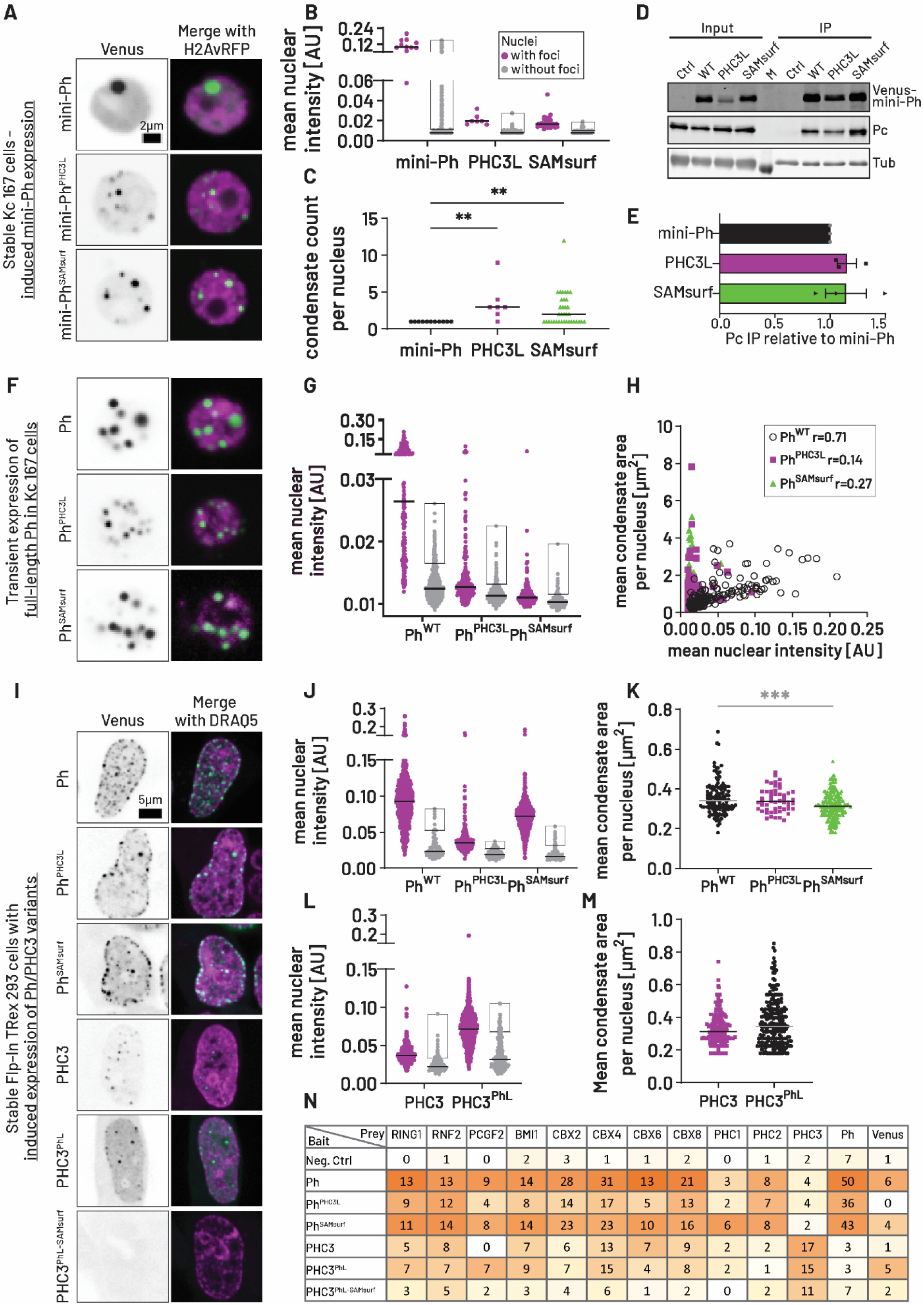
Altering charge complementarity and linker-SAM interactions affects condensate formation in cells. A. Representative images of induced Venus-mini-Ph variants in stable Kc167 cell lines (individual z-slice, H2Av-RFP is a nuclear marker). B. To compare thresholds for condensate formation, cells were stratified with (magenta) and without (gray) condensates and the mean intensity of Venus per nucleus plotted. Line indicates median and gray box the 10% highest expressing cells without condensates. Data are compiled from three experiments. mini-Ph WT n=2796 cells (11 with condensates); PHC3L n=2295 (7 with condensates); SAMsurf n=1369 cells (36 with condensates). C. Quantification of condensates per nucleus. Comparisons are to mini-Ph WT; **: p<0.005. D. Representative Western Blot for anti-Venus co-IP from Kc167 lines (control: H2Av-RFP, no Venus transgene). Pc is a PRC1 subunit; α-Tubulin (Tub) was included as a loading control and remains associated to the anti-Venus resin despite stringent washing. E. Quantification of co-IP experiments (n=3). F. Representative images of overexpressed full-length Venus-Ph variants in transiently transfected Kc167 cells (individual z-slice). Same scale as A. G. Comparison of threshold for condensate formation by full-length Ph variants as in B. Ph WT n=1238 cells (211 with condensates); PHC3L n=1044 (223 with condensates); SAMsurf n=868 cells (309 with condensates). H. Quantification of condensate size dependence on Venus-Ph concentration as in C. I. Representative images of overexpressed fly Ph variants and human PHC3 variants in stable Flp-In TRex 293 cell lines (individual z-slice, with DRAQ5 staining as nuclear marker) induced with 1 µg/ml Dox for 1 day. All constructs except PHC3^PhL-SAMsurf^ form condensates. J. Comparison of thresholds for condensate formation by Ph variants in Flp-In cells as in B. Ph n=1276 cells (1175 with condensates); Ph^PHC3L^ n=702 (234 with condensates); Ph^SAMsurf^ n=1371 cells (1177 with condensates). K. Mean condensate sizes per nucleus from cells with similar Ph expression levels. Cells were selected based on a window of nuclear mean intensities spanning the 5^th^ to 25^th^ percentile in the Ph^SAMsurf^ dataset. The datasets were compared against Ph^WT^; ***: p<0.0005. L. Comparison of thresholds for condensate formation by PHC3 variants in Flp-In cells as in B. PHC3 n=1241 cells (403 with condensates); PhL n=1482 (1157 with condensates). M. Mean condensate sizes per nucleus from cells with similar PHC3 expression levels. Same as in K except the expression window (5^th^ to 25^th^ percentile) is from the PHC3^PhL^ dataset. Differences are not significant (Mann-Whitney test). N. Summary of Co-IP followed by mass spectrometry from Flp-In cells showing averaged unique peptides for PcG proteins identified in two replicates. Colour code represents the fraction of all unique peptides identified across samples for each protein in each sample (white =0%, orange=100%).

We previously showed that the large, disordered N-terminal region of Ph, which is not present in mini-Ph (**Figure 1A**) affects the number and properties of condensates that Ph forms in cells^31,57^. Niekamp and colleagues confirmed that the N-terminal region of PHC1 affects condensate formation by PRC1 *in vitro*^39^. To determine whether effects of the PHC3L and SAMsurf mutation remain relevant in the context of FL Ph with the N-terminal IDR, we introduced FL Venus-tagged Ph variants into Kc167 cells by transient transfection (**Figure 6F**), and carried out the same analysis described above. Again Ph is expressed at higher levels than either mutant (**Figure 6G**). The Venus intensity levels for cells with condensates overlap the range where Ph does not form condensates, consistent with a lower threshold (**Figure 6G**). There is substantial overlap between Venus intensities in cells with and without condensates, indicating that factors other than protein concentration influence condensate formation. Plotting mean condensate size against mean nuclear Venus intensity indicates positive correlations between the parameters for all three proteins (**Figure 6H**). However, the correlation is strongest for Ph, for which condensates steadily increase in size with expression levels. In contrast, for Ph^PHC3L^ and Ph^SAMsurf^, a wide range of condensate sizes is observed over a small range of protein concentrations (**Figure 6H**). Comparison of condensate counts and sizes indicates that Ph forms the smallest condensates (at equivalent expression levels) (**Figure S6Bi**) but slightly more condensates compared to Ph^PHC3L^ and Ph^SAMsurf^ (**Figure S6Bii**). We conclude that the threshold for condensate formation is highest for Ph, and that both PHC3L and SAMsurf change how condensate size is regulated. These effects are consistent with those observed in mini-Ph constructs, indicating that the influence of PHC3L and SAMsurf on phase separation is maintained in the context of FL Ph.

To test the role of polymerization in condensate formation by the various mini-Ph proteins, we used transient transfection in Kc167 cells with CuSO_4_-mediated induction of expression (**Figure S6Ci, Di**). We compared mini-Ph, mini-Ph^PHC3L^, and mini-Ph^SAMsurf^ to the same proteins with the EH polymerization mutation. As observed in stable lines (**Figure 6A**), the PHC3L and SAMsurf mutations form condensates at lower expression levels than mini-Ph (**Figure S6Cii**). With the EH mutation in the mini-Ph variants, only the highest expressing cells formed condensates, particularly for the SAMsurf mutations (**Figure S6Dii**). This could reflect a shifted threshold for condensate formation, distinct from what is observed *in vitro*, or lower stability of condensates (consistent with previous *in vitro* results^31,39^). mini-Ph variants with the EH mutation were also expressed at lower levels (assessed by Western blot, **Figure S7A**), which may explain the small number of cells with condensates. We also tested FL Ph, Ph^PHC3L^ and Ph^SAMsurf^ with the EH mutation using transient transfection (**Figure S6Ei**). Although Ph^PHC3L^ and Ph^SAMsurf^ have lower expression levels than Ph (for both WT and EH mutants), many cells form multiple condensates, and Ph^PHC3L-EH^ forms condensates at the lowest expression levels, followed by Ph^SAMsurf^ (**Figure S6Eii**). Comparing condensates in cells within the same expression window, the size is similar, and the numbers of condensates in the whole population of cells is also similar (**Figure S6Eiv**). However, as for the same proteins with the intact SAM, size and expression level are well correlated for Ph^EH^ but less so for Ph^PHC3L-EH^ or Ph^SAMsurf-EH^ (**Figure S6Eiii**). This indicates that altered condensate size control due to PHC3L or SAMsurf mutations does not depend on SAM polymerization, consistent with our *in vitro* findings.

To determine if the differences in condensates caused by changing the linker or mutating the SAM surface are specific to *Drosophila* cells or can also occur in human cells where the PHC3L normally functions, we created stable (Flp-IN) human cell lines for inducible expression of (*Drosophila*) Ph, Ph^PHC3L^ and Ph^SAMsurf^. We also tested chimeric proteins in which the PhL is introduced into PHC3 (PHC3, PHC3^PhL^, or PHC3^PhL-PHC3-SAMsurf^), all with an N-terminal Venus tag. We used live imaging to characterize the condensates formed as above (**Figure 6I**). The pattern of condensates formed for all proteins in human cells is distinct from that in *Drosophila* cells in that Venus signal and condensates often accumulate at the nuclear periphery.

All of the proteins formed nuclear condensates when induced, with the exception of PHC3^PhL-PHC3-SAMsurf^, which was diffusely localized in both nucleus and cytoplasm. This result can be understood from the expected interactions in this protein (**Figure 5J)**—in PHC3, the FCS interacts strongly with the linker, and the linker in turn with the SAM. In PHC3^PhL-PHC3-SAMsurf^, PhL should interact with the PHC3 SAM, but the PHC3 FCS will likely not interact with PhL so that the coupling from FCS to SAM mediated by the linker is absent. We measured nuclear Venus intensity and condensate size as above and plotted nuclear Venus intensities for cells as with or without condensates as above. As observed in *Drosophila* cells, Ph is expressed at higher levels than Ph^PHC3L^ or Ph^SAMsurf^, and Ph^PHC3L^ forms condensates at lower expression levels than Ph (**Figure 6J**). For Ph^SAMsurf^, which is expressed at higher levels than Ph^PHC3L^, the range of intensities over which condensates are observed is similar to that for Ph (**Figure 6J**). The correlation between condensate size and expression levels were similar and positive for all three proteins (**Figure S6Fi**). Ph forms larger condensates than Ph^SAMsurf^ in cells in a matched expression range (**Figure 6K**), and more condensates than either mutant protein (**Figure S6Fii**). We also compared PHC3 with PHC3^PhL^; intriguingly, PHC3^PhL^ is expressed at higher levels than PHC3. PHC3 forms condensates at lower expression levels; condensate size in cells from a matched expression window is similar although PHC3^PhL^ forms more condensates (**Figure 6L, M**, **S6Gii**). Although condensate size and expression level are positively correlated, the correlations are not strong for either PHC3 or PHC3^PhL^, with a wide range of condensate sizes over a narrow concentration range (**Figure S6Gi**). We conclude that the linker and SAMsurf mutations affect condensate formation and properties in mammalian cells, although additional mechanisms regulating condensate size may be used by PHC3, which requires further study.

To determine if ectopically expressed Ph proteins assemble into PRC1 in human cells, we carried out immunoprecipitations with anti-GFP resin followed by mass spectrometry. We find that all Ph proteins co-IP PRC1 components (**Figure 6N**). PHC3 variants co-precipitated fewer PRC1 peptides than Ph variants, which may reflect their lower expression. We conclude that in both *Drosophila* and human cells, the PHC3L and PhL+SAMsurf promote condensate formation at lower concentrations than PhL, and regulate condensate size. The finding that proteins containing the SAMsurf mutations and the PHC3L tend to be expressed at lower levels than those with PhL in both human and *Drosophila* cells suggests a possible unexpected role for condensates in regulating protein levels.

### The PHC3 linker and SAM surface mutations affect condensate formation by endogenous Ph

To determine if the changes in interactions induced by PHC3L and SAMsurf mutations affect condensates formed at endogenous Ph levels, we introduced these changes into endogenous Ph. We used CRISPaint editing, which depends on non-homologous end joining (NHEJ) and is highly efficient in *Drosophila* tissue culture cells^58,59^, in *Drosophila* Kc167 cells. There are two tandem *ph* genes in the *Drosophila* genome, *ph-p* and *ph-d*. All of our work uses the protein encoded by *ph-p*. *ph-p* and *ph-d* are highly similar at both the DNA and protein levels, with the main difference being that *ph-d* has a shorter N-terminal disordered region. Both *ph-p* and *ph-d* express FL proteins (predicted 167 and 162 kDa) and smaller isoforms predicted to be ∼140kDa (Flybase, FB2024_06). Thus, tagging the N-terminus of *ph* will not result in full protein tagging. The SAM is present at the C-terminus of *ph*, and previous work showed that a C-terminal GFP tag can interfere with condensate formation in mammalian cells^21^. We therefore inserted an internal Venus tag along with the sequence changes to the linker or SAM to be able to image condensates. The Venus tag was placed between the N-terminal disordered region and the HD1 (i.e. the start of mini-Ph) (**Figure S7C**). This is the same placement of Venus in our mini-Ph constructs used for imaging (**Figure 6A**). We obtained one single cell clone for each of Ph, Ph^PHC3L^, and Ph^SAMsurf^. For the Ph line, we detected one in-frame edited allele with Venus inserted into the *ph-p* gene as well as *ph-p* and *ph-d* truncated near the editing site (thus not producing protein with the mini-Ph region) (**Table S3**). For the Ph^PHC3L^, line we detected one in-frame edited allele of *ph-p* and one with a deletion removing amino acids 1287-1561 containing the HD1 but remaining in frame (**Table S3**). The HD1 is required for assembly into PRC1^60^. For Ph^SAMsurf.^, we only detected perfectly edited *ph-p* (**Table S3**). We did not detect either intact or truncated *ph-p*, and were not able to amplify *ph-d* at all. Thus, in these cell lines, the only FL Ph that should be expressed is Venus tagged *ph-p*. We measured Venus-Ph protein levels in the cell lines and find that protein levels of Ph^PHC3L^ are modestly reduced relative to Ph and Ph^SAMsurf^ consistent with the behavior of ectopically expressed proteins (**Figure 7B, S7F**). We also used co-IP to test if Venus-Ph proteins assemble into PRC1 and find that they do (**Figure S7F, G**). These cell lines provide a suitable system to analyze Ph condensates at endogenous levels or slightly reduced protein levels while maintaining PRC1 assembly.

**Figure 7:**
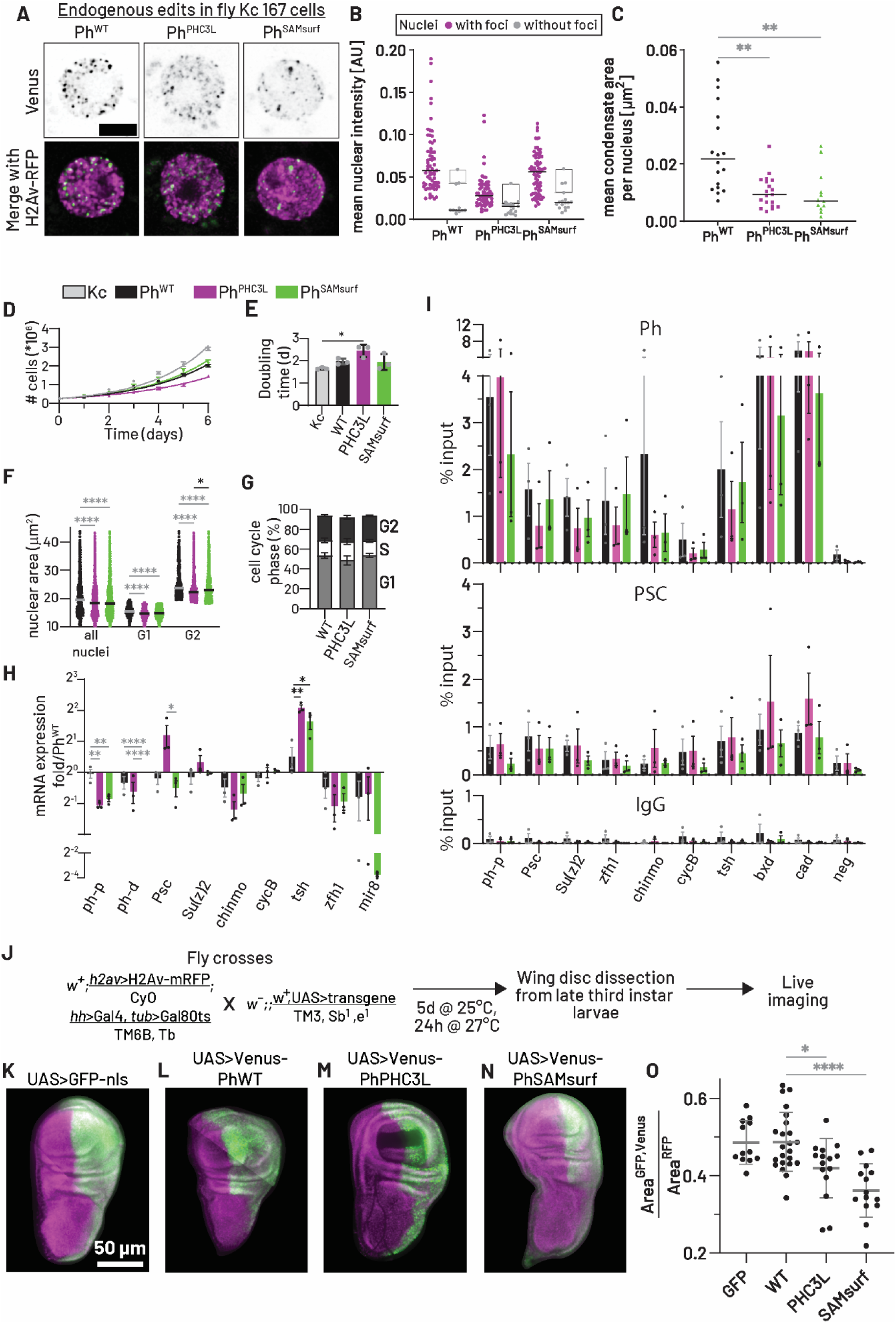
Effects of altering charge complementarity and linker-SAM interactions in endogenous Ph in cells and ectopically expressed Ph in *Drosophila* wing imaginal discs. A. Representative images of Kc167 cells with Venus-tagged endogenous Ph variants transfected with H2Av-RFP as nuclear marker (maximum intensity Z-projection). Scale bar: 2 µm. B. To compare thresholds for condensate formation, cells were stratified into with (magenta) and without (grey) condensates and mean nuclear Venus intensity plotted. Line indicates median and gray box the highest expressing cells without condensates (top 25%). Ph WT n=77 cells (66 with condensates); PHC3L n=81 (63 with condensates); SAMsurf n=93 cells (76 with condensates). ****: p<0.0001. C. Mean condensate sizes per nucleus from cells with similar Ph expression levels (window from 20^th^ to 40^th^ percentile in the Ph^SAMsurf^ dataset). Comparisons are vs. Ph^WT^ **: p<0.005. D. Growth curve of edited and parental Kc167 cells. n=3 experiments; fits are simple exponential growth curves,. E. Doubling times from data in (D). *: p<0.05. F. Quantification of nuclear area from DAPI-stained edited Kc167 cell lines (n=3, colours as D). To compare sizes of nuclei in G1 and G2 the full populations were split at the median. *: p<0.05, ****: p<0.0001 G. Cell cycle profiles of edited Kc167 cells from flow cytometry with Hoechst staining (n=3). H. Gene expression analysis of selected targets using RT-qPCR. *: p<0.05, **: p<0.01, ****: p<0.0001 I. ChIP-qPCR of PRC1 subunits Ph and PSC. Bars are mean +/-SEM. Colours as in D. J. Schematic of experiments to express Ph variants in wing imaginal discs. K-N. Representative wing discs from flies expressing Venus-Ph transgenes. Images are summed z-projections from stitched tiles. The wing discs were cropped and placed on a black background to aid visualization. O. Quantification of the ratio of posterior compartment to total disc area. GFP: n=12; WT: n=23; PHC3L: n=14; SAMsurf: n=15. *: p<0.05, **: p<0.01, ****: p<0.0001

We used live cell imaging of Venus Ph and transfected H2Av-RFP as a nuclear marker. Because the intensity of the signal is low, we used a Stellaris confocal microscope in “Lightening” mode to attain super-resolution. We find that most cells form Venus-Ph condensates for all three cell lines (**Figure 7A**). Comparing intensities for cells that form condensates, Ph^PHC3L^ forms condensates in cells with lower expression levels than Ph, although most Ph expressing cells form condensates and have higher expression levels than Ph^PHC3L^ (**Figure 7B**). We plotted mean condensate area against mean nuclear intensity and observe a positive but low correlation between the two parameters for Ph^PHC3L^ and Ph^SAMsurf,^ (**Figure S7D**) similar to what is observed in transfections (**Figure 6H**). Ph cells form two populations—one of which shows the steady increase in condensate size with expression level observed in transfections, and the other that is more similar to Ph^PHC3L^ and Ph^SAMsurf^ in having a range of condensate sizes over a narrow expression window (**Figure S7D**). As is evident from the scatter plot (**Figure S7D**), Ph forms more numerous condensates, and larger condensates than Ph^PHC3L^ or Ph^SAMsurf^ in a matched expression window (**Figure 7C, S7E**). We conclude that as observed in the other cell-based assays, Ph^PHC3L^ can form condensates at lower expression levels, and both PHC3L and SAMsurf affect condensate size regulation.

### PHC3L and SAMsurf mutations affect nuclear size and PHC3L affects cell growth

Ph is implicated in growth regulation in *Drosophila* imaginal discs so that when it is removed, cell overgrowth and tumour formation is observed^61^. We noticed that Ph^PHC3L^ cells grow slowly and therefore measured the doubling time for each of the Ph lines (**Figure 7D**). We found that Ph^PHC3L^ cells grow more slowly than the parental Kc167 cell line (**Figure 7E**). This is consistent with enhanced Ph function in growth control imparted by the PHC3L. Disruption of long range interactions mediated by PRC1 was shown to result in larger nuclear size in mouse embryonic stem cells^37^. If the PHC3L or the SAMsurf mutations enhance Ph function in long range interactions, it might result in smaller nuclei. We therefore measured nuclear sizes of the Ph lines and found that Ph^PHC3L^ and Ph^SAMsurf^ cells have smaller nuclei than Ph cells (**Figure 7F**). This was true whether we considered all nuclei, or split them into G1 and G2 based on DNA content. This effect cannot be explained by changes in the cell cycle as Ph^PHC3L^ cells have a slightly higher fraction of cells in S-phase and G2 (when nuclei are larger) than Ph cells, while Ph^SAMsurf^ cells have a similar cell cycle profile to Ph cells (**Figure 7G**).

To determine how changing condensates affects Ph function, we measured expression of a small number of target genes in the three cell lines by qRT-PCR (**Figure 7H**). We included *ph-p*, *ph-d* and the PRC1 components *Psc* and *su(z)2*, since these genes are strongly bound by PRC1 and contain functional Polycomb Response Elements^62,63^. *ph-p* and *ph-d* are both downregulated in Ph^PHC3L^ and Ph^SAMsurf^, consistent with the lower levels of protein (**Figure 7H**), while *Psc* is upregulated in Ph^PHC3L^. We also tested several targets previously implicated in growth control by the PcG or misexpressed when PcG genes are deleted in imaginal disc cells^64–68^ (**Figure 7H, Table S4**). The only significant change detected was for the developmental regulator *tsh*, which is upregulated in Ph^PHC3L^ and (slightly less) in Ph^SAMsurf^ cells (**Figure 7H**).

We also tested if Venus Ph proteins are recruited to target genes using ChIP-qPCR (**Figure 7I**). We tested Ph and PSC binding at the *ph-p*, *Psc*, and *su(z)2* promoters, a *Psc* upstream region, and other strong binding sites, including genes implicated in growth control. We did not observe significant changes in binding of Ph or PSC at any sites tested, including *Psc* and *tsh* whose expression is increased. This indicates that PRC1 is recruited to chromatin in all three cell lines (**Figure 7I, Table S5**). Although we cannot rule out that binding is affected at non-tested sites, our data suggest the changes in condensates and gene expression caused by the PHC3L and SAMsurf mutations do not reflect changes in PRC1 recruitment to chromatin.

### PHC3L and SAM surface mutations affect Ph function in *Drosophila* imaginal discs

As noted above, previous work demonstrated that *Drosophila* imaginal discs are sensitive to *ph* levels and activity^46,69^. Overexpressing wild type Ph or Ph with a synthetic linker that enhances polymerization suppresses growth in discs^46^. In contrast, deleting *ph* or impairing SAM polymerization results in tumorous overgrowth^46,69^. To test the effect of the PHC3L and SAMsurf mutations on Ph function, we ectopically expressed wild type Ph, Ph^PHC3L^, Ph^SAMsurf^ or GFP only. We used *hedgehog*-Gal4 and *tubulin*-Gal80ts, which drives expression in the posterior compartment of wing imaginal discs at the permissive temperature (**Figure 7J**). We induced protein expression by temperature shift for 24 hours before harvesting imaginal discs for live imaging (**Figure 7K-N**). All three Ph proteins formed condensates in wing disc cells (**Figure S7H-J**). To assess the effect of the Ph proteins on growth, we measured the ratio of posterior compartment to total disc size (**Figure 7O**). For discs expressing Ph or GFP, this ratio is ∼0.5 (**Figure 7K, L, O**). Discs expressing Ph^PHC3L^, or Ph^SAMsurf^ have reduced ratios (**Figure 7M, N, O**). This is consistent with growth suppression in cells expressing these proteins resulting in a smaller posterior compartment. We measured Venus intensity normalized to the H2Av-RFP nuclear marker in discs, which shows that PhPHC3L is expressed at slightly lower levels than Ph (**Figure S7K**). The integrity of the Venus fusion proteins was confirmed by Western blotting (**Figure S7L**). Thus, in a developing *Drosophila* tissue, the PHC3L and SAMsurf mutations which alter condensate formation and properties, decrease growth.

## Discussion

We investigated the role of a disordered linker in regulating condensate formation driven by the oligomerizing SAM domain of the Polycomb protein Polyhomeotic by comparing *Drosophila* Ph and human PHC3 linkers in the context of the functional core of Ph (mini-Ph). MD simulations revealed predominant interactions between PHC3L and the SAM domain, as well as PHC3L self-interactions, which are predicted to promote phase separation. Using reconstituted condensates of mini-Ph with either PhL or PHC3L and DNA, we found that the PHC3L lowered the threshold for phase separation with DNA, enabled DNA-independent phase separation, and produced smaller condensates. We confirmed the role of PHC3L-SAM interactions by engineering the SAM surface to interact with PhL (mini-Ph^SAMsurf^). Both simulations and phase separation assays showed that engineered interactions qualitatively mimicked the effect of PHC3L. Importantly, the influence of linker-SAM interactions was independent of SAM oligomerization as phase separation persisted even when the SAM polymerization interface was disrupted by mutation. Instead, oligomerization primarily regulated condensate dynamics. To assess the importance of the linker and linker-SAM interactions *in vivo*, we edited endogenous Ph to introduce PHC3L or the SAM surface mutations. Both modifications lowered the apparent threshold for condensate formation and altered condensate sizes without disrupting Ph assembly into PRC1 or its recruitment to target genes. Notably, Ph^PHC3L^ and Ph^SAMsurf^ cells exhibited smaller nuclei, and Ph^PHC3L^ cells showed reduced growth rate. When expressed in developing Drosophila imaginal discs, both variants formed condensates and restricted disc growth. Together, our findings demonstrate that the role of the linker and its predicted interactions in controlling condensate formation showed remarkable consistency *in silico*, in reconstituted condensates, and in cells, further supported by their effects in developing *Drosophila*.

### The role of SAM oligomerization in condensates

Our findings, along with previous studies^39^, suggest that SAM oligomerization primarily influences condensates dynamics and stability rather than their initial formation. mini-Ph proteins with the EH oligomerization mutation formed reconstituted condensates at similar concentration ranges as the wild-type (**Figure 4**) but FRAP assays revealed striking dynamics differences: mini-PH^PHC3L^ showed little recovery (median mobile fraction 0.3, **Table S2**), while mini-PH^PHC3L-EH^ exhibited high mobility (median mobile fraction 0.8, **Table S2**). In our previous work with large chromatin or DNA substrates, mini-Ph^EH^ condensates demonstrated faster FRAP recovery and lower salt resistance compared to mini-Ph condensates^31^. Furthermore, mini-Ph^EH^-chromatin condensates, unlike their mini-Ph counterpart, failed to maintain structural integrity in nuclear extracts^31^. While proteins with the EH mutation can form condensates in cells, particularly in FL form (**Figure S6C-E**)^31^, their stability required further investigation in future studies.

Our data align with the *in vitro* study by Niekamp and colleagues on mammalian PRC1, where the EH mutation in PHC2_short_ or PHC1 did not prevent condensate formation but altered exchange dynamics and morphologies^39^. Most strikingly, PRC1 formed with PHC2_short_ resisted fusion when mixed, while those with PHC2_short_ carrying the EH mutation readily combined^39^. In mouse embryonic stem cells, PHC2_short_ with the EH mutation disrupted CBX2 condensates, while wild-type PHC2_short_ incorporated into them^39^. In contrast, we found that Ph^EH^ still formed condensates in *Drosophila* cells (**Figure S6E**), possibly due to the role of the N-terminal IDR in promoting condensates in cells^57^, an aspect not tested in Niekamp *et al.* Intriguingly, a similar role for SAM oligomerization in restricting condensate dynamics was described for the *C. elegans* protein SOP-2^70^, a PcG system component^71^. SOP-2 comprises of an N-terminal RNA-binding IDR and a SAM, which shares distant homology with that of Ph^49,71^. While the SOP-2 RNA binding IDR drives condensate formation *in vivo* and *in vitro*, the SAM domain is essential only *in vivo*^70,71^. However, mutating the SAM polymerization interface increases SOP-2 mobility within *in vitro* condensates, mirroring the behavior of mini-Ph^EH 31^.

These observations suggest that oligomerization may occur after condensate formation across these systems, potentially facilitated by increased protein concentration within condensates. This hypothesis challenges traditional models of Ph/PHC function, which propose that SAM-mediated oligomerization initiates PRC1 clustering^18,21,72^. It is also distinct from oligomerization-dependent phase separation observed in other systems^73,74^. However, pre-formed oligomers may be relevant for mini-Ph^PHC3L^ condensates formed without DNA, since their formation is strongly disrupted by the EH mutation (**Figure 4A-C**). In contrast, condensates with DNA form at similar concentrations irrespective of the EH mutation, arguing against a strict requirement for pre-formed oligomers.

Regulation of condensate stability through oligomerization could serve as a mechanism to reinforce gene repression. In this regard, SUMOylation has been implicated in the regulation of PcG bodies across multiple species, including mammals, *Drosophila* and *C. elegans* ^70,75–77^. Consistent with this, a SUMO tag affects condensates formed by PHC3L-SAM (**Figure S2J**). Additionally, Ph is heavily modified by O-linked glycosylation of its N-terminal IDR, which was shown to restrict Ph aggregation in a polymerization-dependent manner^24^. Deletion of the O-GlcNac transferase gene (*ogt*) or the modified region of Ph results in similar defects in gene repression^24^, further suggesting that post-translational modifications fine-tune condensate stability through the SAM to regulate gene expression.

### Interactions of flexible linker underly phase separation by the functional core of Ph

The distinct properties of condensates formed by mini-Ph, mini-Ph^PHC3L,^ and mini-Ph^SAMsurf^ with DNA arises from their unique networks of molecular interactions (**Figure 5J**), despite sharing FCS-DNA interactions and SAM-SAM mediated oligomerization. Since all three proteins can undergo phase separation with DNA in the absence of SAM oligomerization (**Figure 4**), and FCS-DNA interactions are insufficient for phase separation^31^, additional interactions must be required. In mini-Ph, these additional interactions include non-canonical SAM-SAM interactions (**Figure S3A**), and FCS-SAM interactions (**Figure 5A**). Neither of these interactions is present in mini-Ph^SAMsurf^ (**Figure 3C**, **Figure 5E, J**), yet mini-Ph^SAMsurf^ still phase separates, suggesting that linker-SAM interactions compensate for the absence of SAM-SAM and FCS-SAM interactions. The mini-Ph^SAMsurf-EH^ presents the most minimal interaction network, with phase separation likely governed by FCS-linker, linker-SAM, and FCS-DNA interactions.

While mini-Ph^PHC3L^ shares a similar overall interaction pattern with mini-Ph, it exhibits distinct predicted interactions in the HD1/FCS region. Additionally, mini-Ph^PHC3L^ features unique PHC3L-PHC3L, and PHC3L-SAM interactions, and PHC3 itself can bind DNA (**Figure S3D**). All of these interactions may contribute to phase separation. Our simulations of (human) mini-PHC3 reveal an interaction network most closely resembling mini-Ph^SAMsurf^, with the addition of PHC3L-PHC3L interactions. This change is due to the HD1-FCS region in PHC3 that remove FCS-SAM contacts present in the *Drosophila* protein (**Figure 5H, I, S5F**).

As noted in the Results, the only protein we tested that did not form condensates in cells (PHC3^PhL-PHC3-SAMsurf^, **Figure 6I**) may lack effective coupling among domains. Importantly, the interactions in mini-Ph are maintained in simulations of the FL protein (**Figure S5A**). The agreement between *in vitro* and *in vivo* condensate formation with different sequences further supports the relevance of this molecular interaction model for Ph function in PRC1. Together, our computational and experimental findings highlight the functional core of Ph as highly interconnected and sensitive to sequence variations, features that have likely been exploited through evolution.

### Linker control of condensate size

In both reconstituted condensates and cells, we consistently observed differences in the sizes of condensates formed with PhL, PHC3L, or the SAM surface mutations. Many mechanisms have been described for condensate size control and the coexistence of multiple condensates within cells. These include saturation of valencies^78^, sub-diffusive movement in the cellular environment coupled to periodic dissolution and reformation of condensates during mitosis^79^, and condensate substructure. Substructure-based size control can involve components with surfactant properties ^80^, proteins that bind to condensate surfaces acting as surfactants^81^ or Pickering particles^82^ that block condensate fusion. Under certain conditions, condensates can also undergo kinetic arrest that impairs fusion, passing rapidly through a liquid phase to a gel or even solid, but remaining reversible^83^. This may be relevant for condensates that form and dissolve each cell cycle but are not dynamic (like centrosomes)^11^. If SAM oligomerization is triggered by condensate formation, it could contribute to kinetic arrest, an idea that is consistent with the non-fusing PRC1-chromatin condensates observed by Niekamp *et al.*^39^ that depend on the SAM polymerization interface. *In vivo*, condensates likely have a complex architecture with multiple components that contribute to size regulation.

*In vitro*, we did not observe obvious substructure in mini-Ph-DNA condensates. Valency saturation is a possible explanation for the observed size differences, where condensate growth halts once most available interaction sites are occupied. In this model, condensate fusion depends on interactions between molecules in each droplets and strong interactions with slower dissociation kinetics reduce fusion likelihood^78,84^. Since both PHC3L and SAM surface mutations enhance protein-protein interactions, they may promote valency saturation and slower dissociation kinetics, ultimately influencing condensate size. However, we must emphasize that the precise mechanism of size control remains an open question for future studies as neither our experiments nor simulations directly address this issue.

In addition to the intrinsic size control observed in reconstituted condensates, the chromatin environment is likely an important determinant of size control in cells. Chromatin has been shown to mechanically restrict condensate growth^85^, and stronger chromatin binding correlates with smaller condensate sizes in other systems^86,87^. Consistent with this, our previous work demonstrated that Ph without its N-terminal IDR (i.e., mini-Ph) exhibited weaker association with chromatin and formed larger condensates^57^, whereas the glutamine-rich subregion of the IDR promoted strong chromatin binding and smaller condensate formation^57^.

A more intriguing question is why condensate size could matter for function. Recent studies in *Ashbya gossypii* have shown that the same regulatory protein can differentially affect RNA translation in small versus large condensates^88^, likely due to distinct properties between condensate interfaces versus interiors^88^. Addressing similar question in Ph condensates will require the development of new tools to analyze condensate architecture in cells and investigate chromatin locus positioning, and the establishment *in vitro* activity assays where condensate size and activity can be directly compared. The ability to modulate condensate size through the linker provides a valuable approach for investigating the functions of condensate size variation.

### Regulation of growth by Ph

Introducing PHC3L into Ph alters growth in tissue culture cells (**Figure 7D, E**), and both PHC3L and the SAM surface mutations influence growth control in wing discs (**Figure 7 J-O**). It is well established that Ph has an important function in growth regulation in imaginal discs, a system that is highly sensitive to Ph function. Both increased and decreased Ph activity affects growth homeostasis^46,67–69,89–92^, with even transient loss of Ph leading to tumorous overgrowth^61,65^. However, identifying key target genes regulated by PRC1 in growth control has proven challenging, as different genes and pathways were identified in different discs and experimental paradigms^92–95^. To investigate potential transcriptional effects, we tested expression of several genes implicated in growth regulation^64–68^ did not identify significant differences in expression in cell lines (**Figure 7H, Table S4**). Similarly, we found no major changes in chromatin binding of Ph mutants, suggesting their effects may occur downstream of chromatin binding (**Figure 7I, Table S5**). Future genome wide analysis of these lines may clarify the mechanisms underlying Ph-mediated growth regulation.

Ph^PHC3L^ and Ph^SAMsurf^ restrict growth, whereas loss of Ph or expression of Ph with a mutated polymerization interface leads to overgrowth^46,61,90^. Therefore, we predict that Ph^PHC3L^ and Ph^SAMsurf^ act as stronger repressors than wild-type Ph, though this prediction remains to be tested. Additionally, condensates may play a role in coordinating expression of multiple growth-regulatory genes, leading to subtle, network-wide effects rather than strong repression of a small set of genes. Notably, the upregulation of two genes (*Psc* and *tsh*) in Ph^PHC3L^ lines (and *tsh* in Ph^SAMsurf^) (**Figure 7H**), suggests *reduced* Ph function at these loci, despite retained PRC1 binding (**Figure 7I**), highlighting complex effects of condensates on gene expression. The smaller nuclei observed in Ph^PHC3L^ and Ph^SAMsurf^ cells (**Figure 7F)** may reflect stronger or more extensive Ph-mediated long-range interactions, as previous observation with Ph overexpression^22^. In mammals, while PcG proteins are implicated in certain cancers, specifical roles for PHC proteins in cancer progression remain largely unexplored^94,96^. Intriguingly, PHC3 has been identified as a tumor suppressor in osteosarcoma^97,98^ and a PHC1 SAM mutation has been linked to microcephaly^99^. These observations highlight the broader significance of Ph proteins and underscore the importance of understanding how condensate formation contributes to growth regulation.

In summary, our work reveals a mechanism by which Ph SAM activity has evolved while maintaining overall protein architecture and functional coupling within the Ph core domains. This adaptation occurs through sequence changes in both the linker (an IDR) and the HD1/FCS region, which modulate condensate formation and properties. These findings highlight how regulated condensates contribute to Ph function in chromatin organization. More broadly, this work demonstrates how small sequence variations can fine-tune condensate behavior, allowing the same macroscopic event—condensate formation—to emerge from distinct molecular interactions, resulting in condensates with diverse properties.

### Limitations

Our biochemical experiments use the functional core of Ph, but do not include its large N-terminal IDR or its binding partners in PRC1. We also used small DNA substrates, but the physiological substrate for Ph is chromatin. While our minimal system clearly captures relevant interactions, developing more complex reconstituted systems to mimic the biological context more closely is an important future goal. While our MD simulations capture aspects of native SAM oligomers, they may not fully represent the relative populations of non-native transient complexes^100^. As a result, we were unable to directly assess the effects of PHC3L on oligomerization or determine the role of oligomerization in phase separation. Our assessment of Ph^PHC3L^ and Ph^SAMsurf^ binding to chromatin and regulation of gene expression was restricted to a small number of targets; genome-wide analysis could clarify how the PHC3L affects growth, and whether these proteins are more repressive than wild-type. We also have not analyzed endogenous Ph with a SAM oligomerization mutation. Finally, in cells, the Ph SAM is believed to interact with two other SAM containing Polycomb proteins, Sex Comb on the Midleg (SCM) and Scm-related gene containing four mbt domains (Sfmbt). Specifically, Ph SAM can form a heteropolymer with the SAM of SCM^101^, and the SCM SAM can also bind to the non-polymerizing SAM of Sfmbt. Sfmbt is part of the Pho-RC complex that is important for recruitment of PRC1 to chromatin^102–105^ so that a SAM network is hypothesized to anchor PRC1 to specific sites^104^. It will be fascinating to consider the role of SCM and Sfmbt SAMs in condensates.

## Acknowledgments

The authors thank Dr. François Robert for critical reading of the manuscript, and Dr. David Hipfner and his lab for resources and advice on *Drosophila* experiments. We thank the IRCM Microscopy, Flow Cytometry and Mass Spectrometry Platforms for their assistance, Drs. Marlene Oeffinger and “Sonny” Pudchalaluck Panichnantakul for input on mass spectrometry and mammalian cell experiments, and Drs. Stefan Niekamp and Robert Kingston for sharing unpublished work. We are grateful for the computation resources provided by Texas A&M High Performance Research (HPRC).

This work was supported by: the Canada 150 Research Chairs program (C150-2017-00015, BD), the CFI (CFI-37589, BD), the NIH (1R01GM120600, BD), (R01GM114338 to CAK & NJF), (R15CA26780, CAK), (R35GM153388, JM), NSERC (DG-RGPIN-2019-05637, BD), (DG-RGPIN –2019-05040, NJF), the CIHR (469023, NJF) and the Welch Foundation (Grant A-2113-20220331, JM). UltraScan supercomputer calculations were supported through NSF/XSEDE grant TG-MCB070039N, and University of Texas grant TG457201 to BD. AH was supported by a scholarship grant from NSERC and TMG by doctoral fellowships from FRQS and the IRCM (Jacques-Gauthier Tribute Scholarship). YCK is supported by the Office of Naval Research through the U.S. Naval Research Laboratory base program. This work was partially conducted at the Structural Biology Platform of the Université de Montréal (RRID:SCR_022303). The Platform was funded by a grant from the Canada Foundation for Innovation (#30574) and is currently supported by the Centre d’Innovation Biomédicale of the Université de Montréal.

## Author contributions

TMG: conceptualization, investigation, formal analysis, visualization, writing—original draft, writing—review & editing; RMR: conceptualization, investigation, visualization, formal analysis, writing—original draft; TMP: investigation, visualization, formal analysis, writing—review & editing; NP: investigation; JS: investigation; OS, AJM, HJL, AH: investigation; DL: investigation, formal analysis, visualization; UK: investigation; AB: investigation; DS: methodology; YCK: investigation, formal analysis; DJ: supervision, resources; BD & CAK: conceptualization, resources, supervision, funding acquisition; JM: conceptualization, writing-original draft, writing—review & editing, funding acquisition, supervision; NJF: conceptualization, writing—original draft, writing—review & editing, funding acquisition, supervision.

## Materials & methods

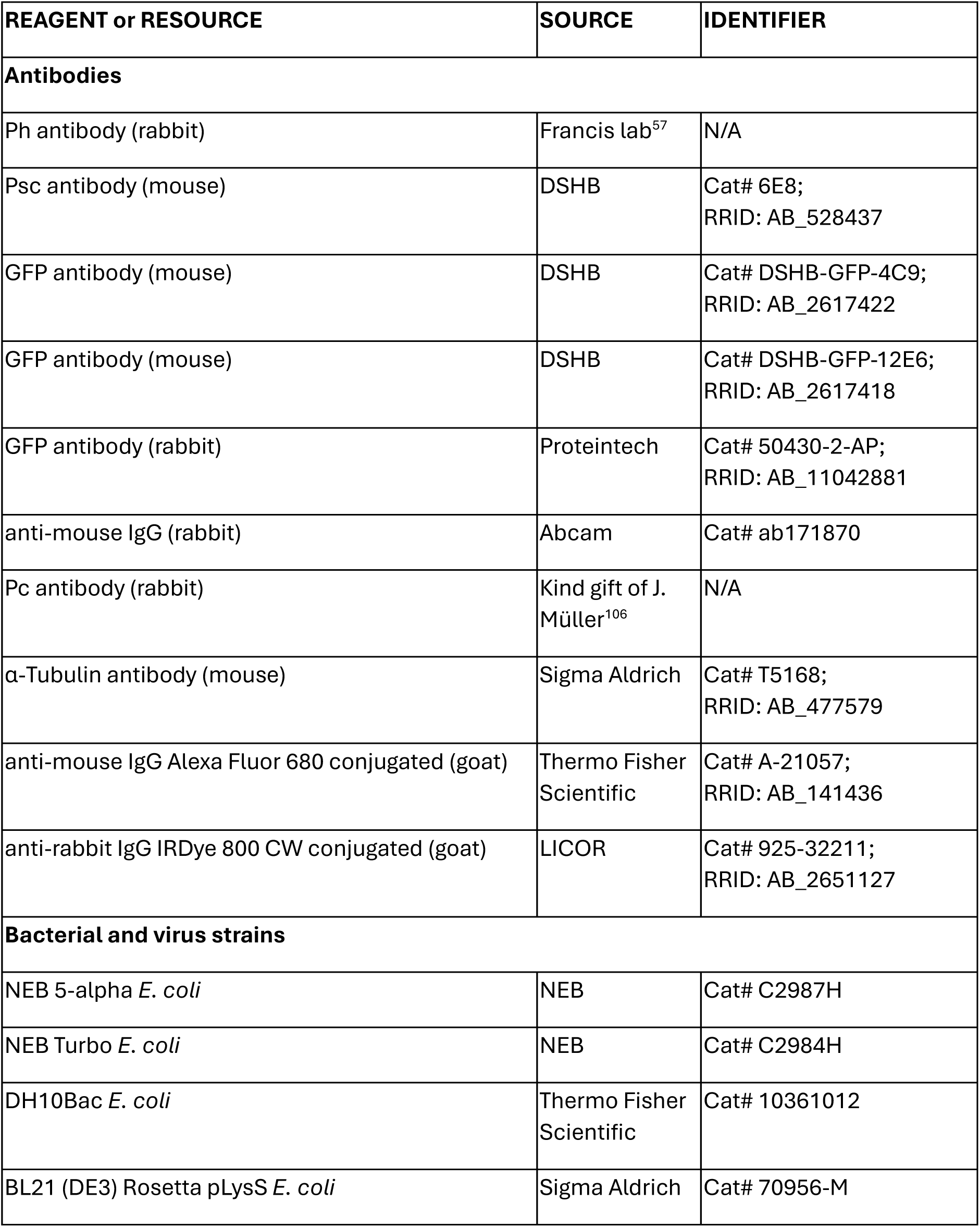
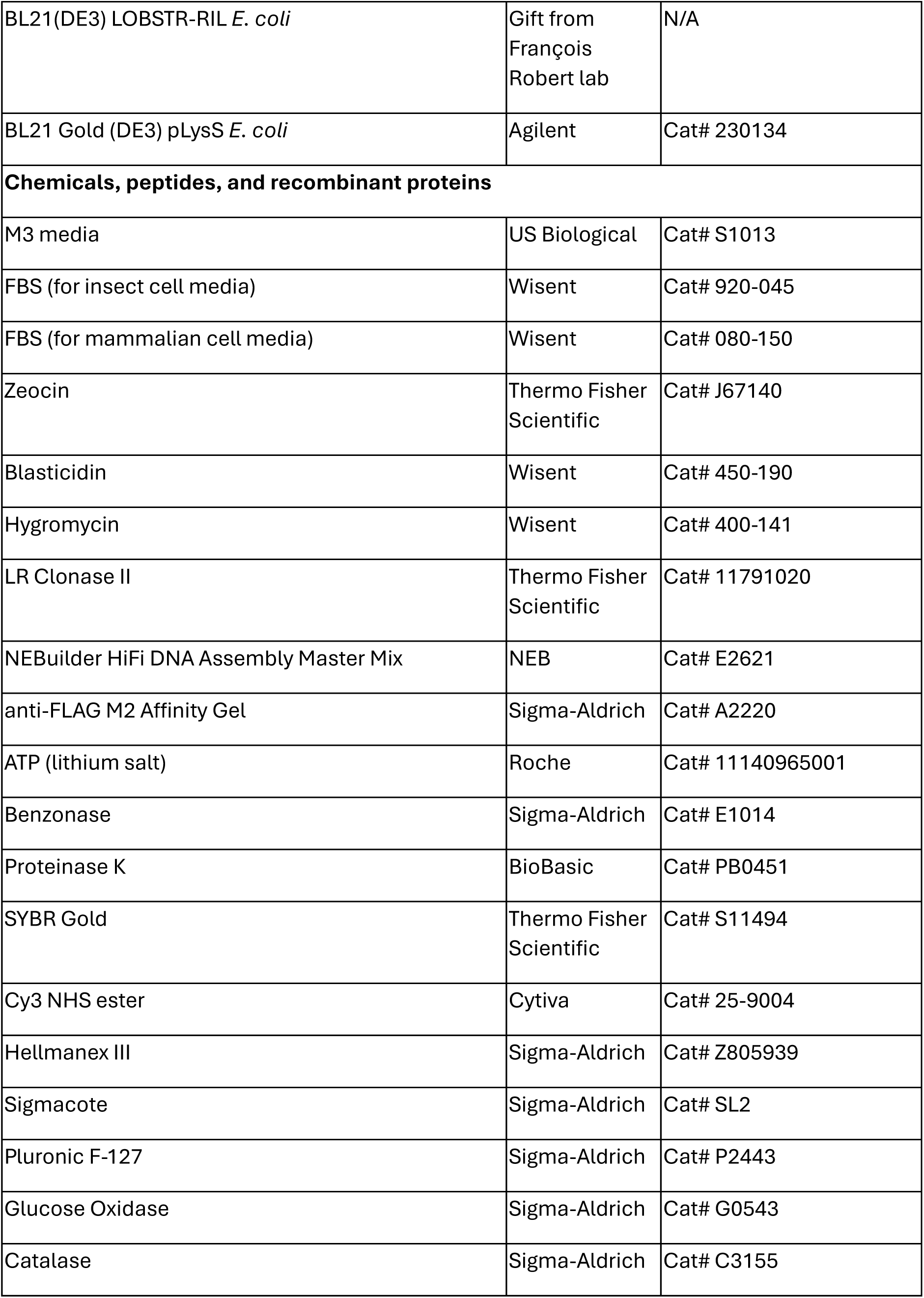
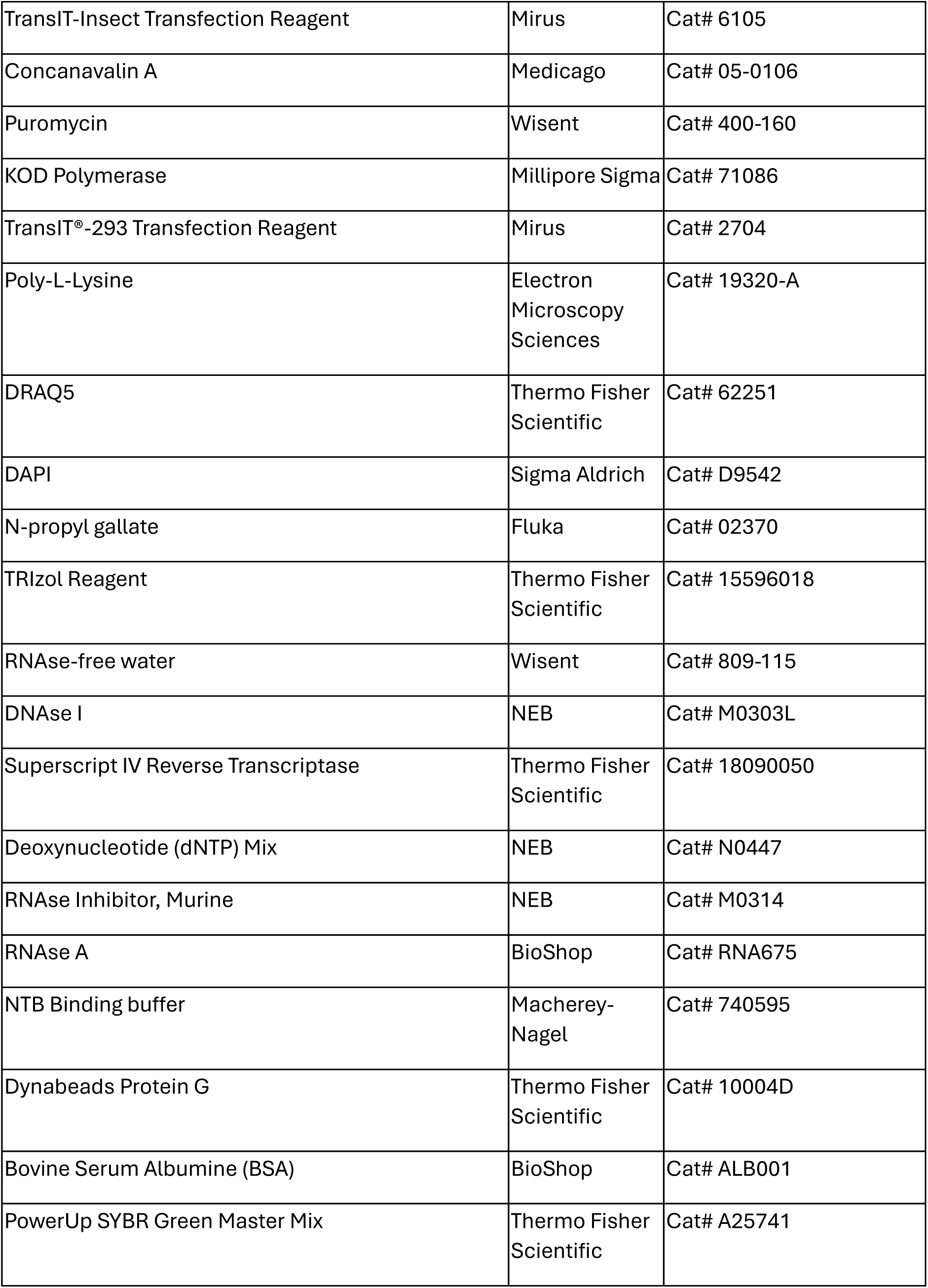
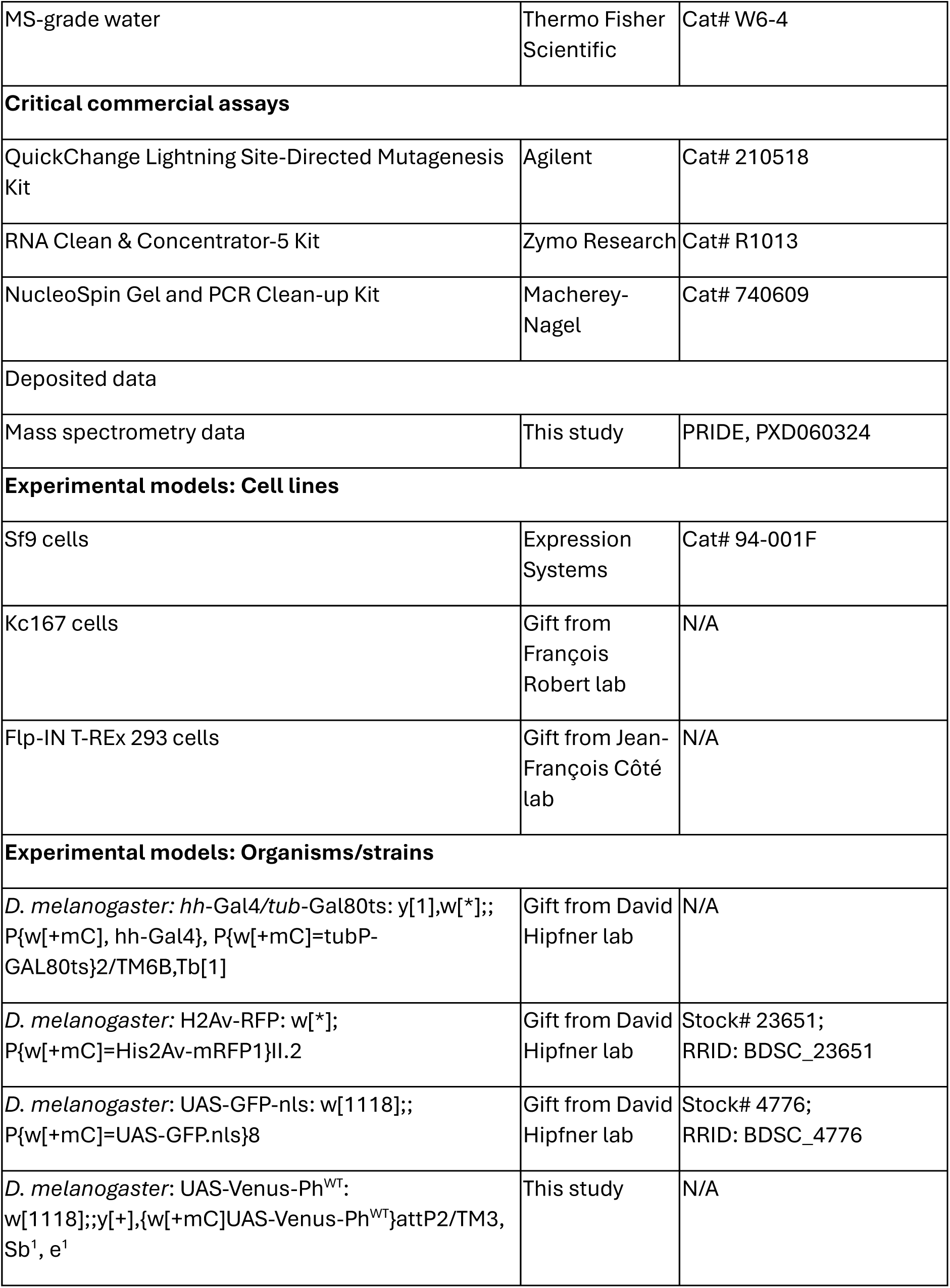
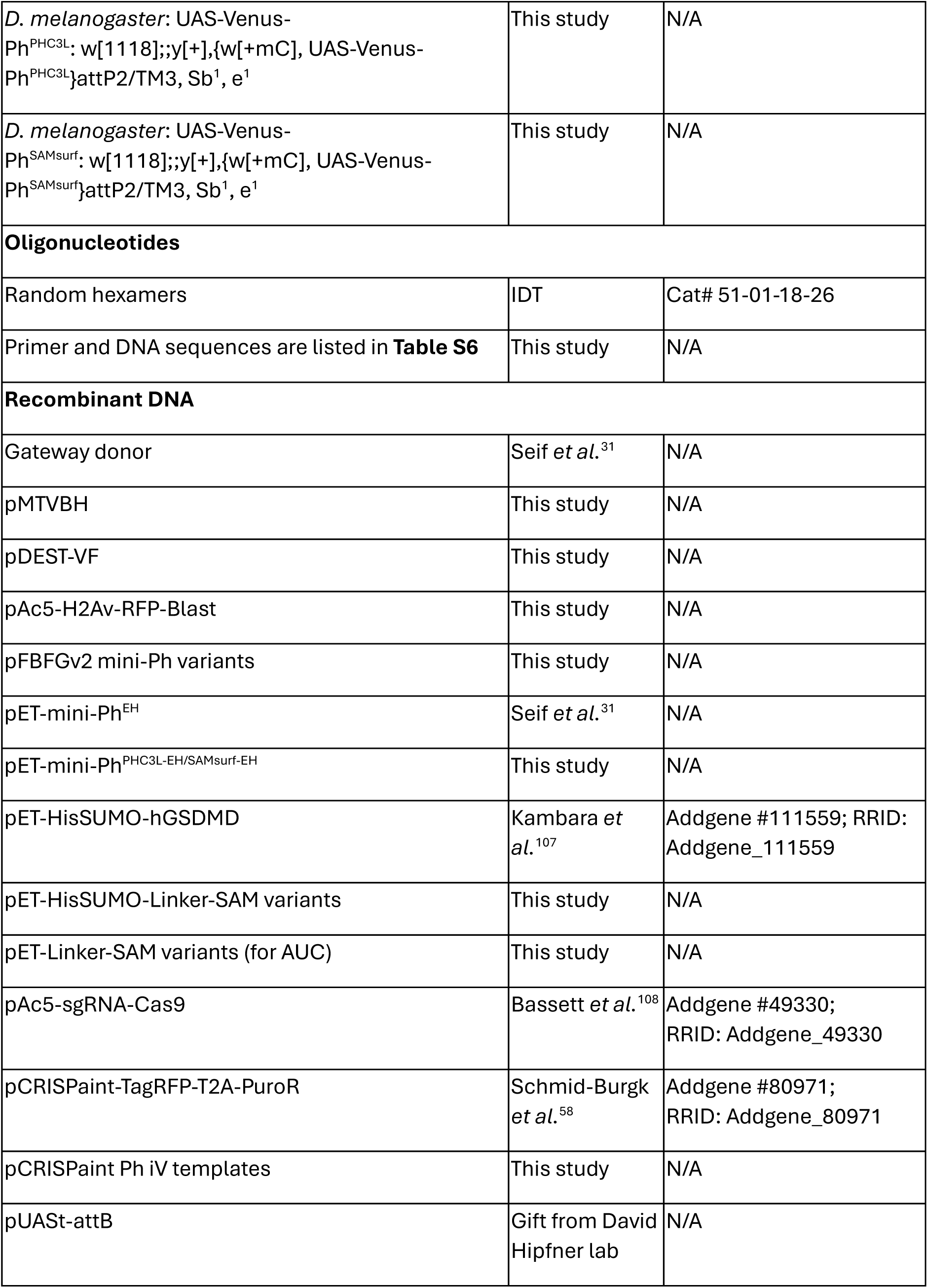
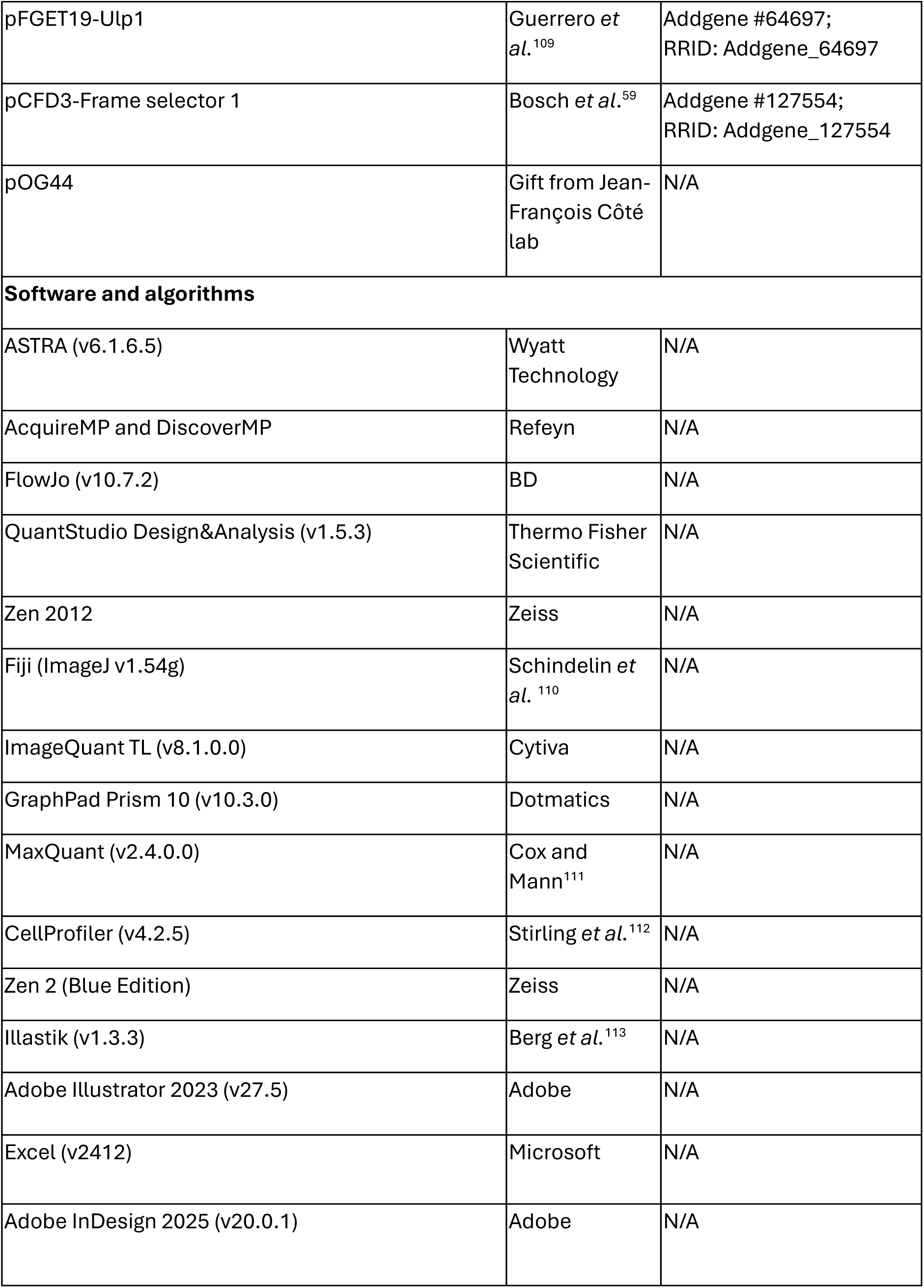
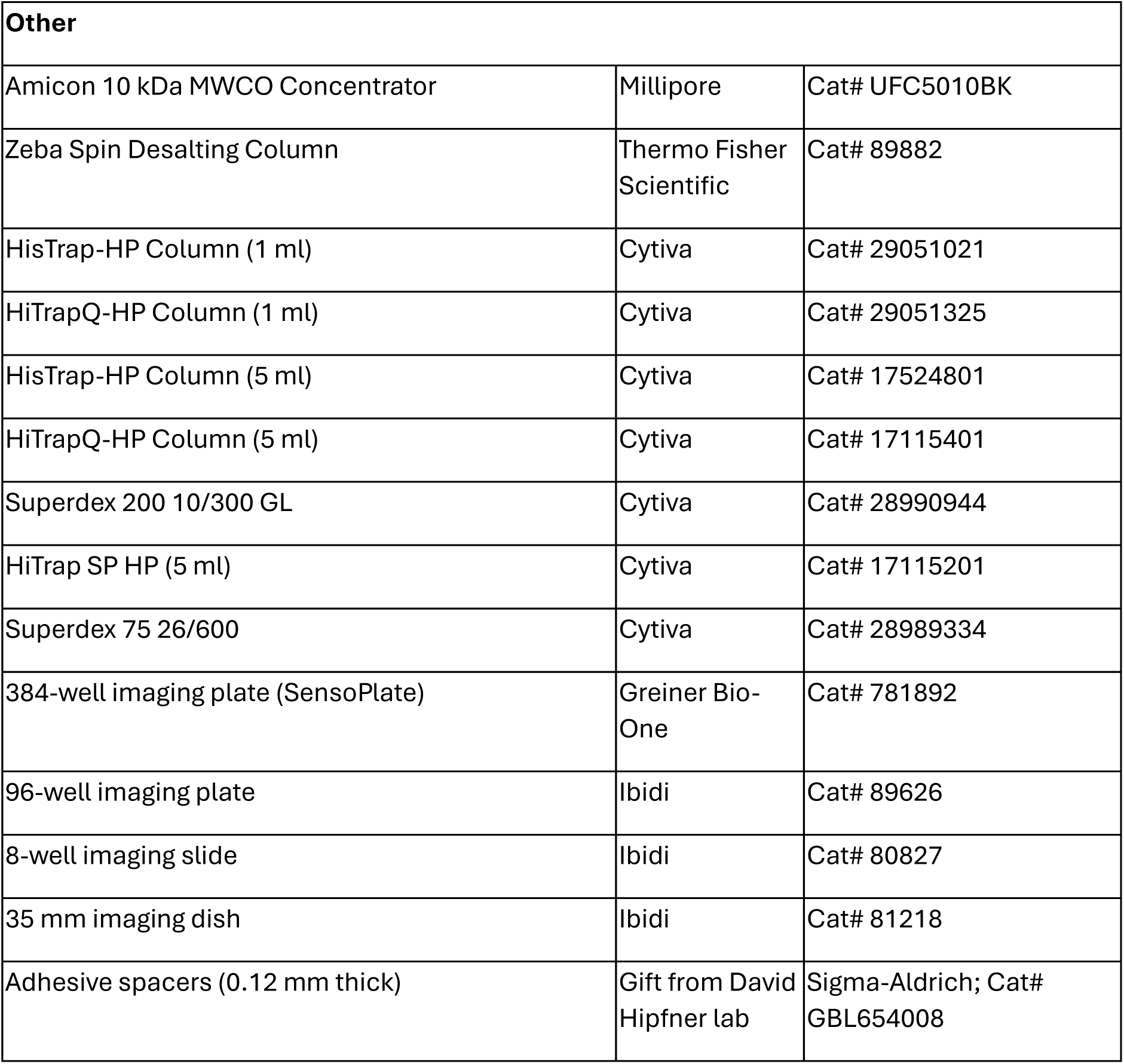
Key resources.

## Key resources

## Resource availability

### Lead contact

Requests for information, resources or reagents should be directed to the lead contact, Nicole Francis

### Materials availability

Materials generated in this study (plasmids, fly strains, cell lines) are available on reasonable request to the lead contact.

## Data and code availability

● Mass spectrometry data generated in this work has been deposited on PRIDE, PXD060324
● This paper does not report original code.
● Data from this paper and information required to reanalyze it are available from the lead contact on reasonable request.

## Experimental model details

### Sf9 cells

Sf9 cells (*Spodoptera frugiperda* cell line) (Expression Systems) were cultured in ESF 921 medium (Expression Systems) in a shaking incubator at 27°C.

### Kc167 cells

*Drosophila* Kc167 cells were cultured in M3 media (US Biological) with 5% (v/v) FBS (Wisent, Cat# 920-045) and 1% (v/v) Penicillin/Streptomycin (Wisent) in a room temperature incubator.

### Flp-IN T-REx 293 cells

Flp-IN T-REx 293 cells were cultured in DMEM (Wisent) with 10% (v/v) FBS (Wisent, Cat# 080-150) and 1% (v/v) Penicillin/Streptomycin at 37°C with 5% CO2. Unmodified cells were maintained with 50 µg/ml Zeocin (Thermo Fisher Scientific) and 6 µg/ml Blasticidin (Wisent), while cell lines with transgenes (details below) were maintained with 100 µg/ml Hygromycin (Wisent) and 6 µg/ml Blasticidin.

### Drosophila melanogaster

Detailed descriptions of fly strains and their origin can be found in the key resources table. Flies were raised at 25°C on standard food. The driver line used in this study was obtained by crossing *hh*-Gal4/*tub* Gal80 flies with H2Av-RFP flies. The generation of transgenic flies with UAS-Venus Ph variants is described in the method details.

## Method details

### Cloning

NEB 5-alpha *E. coli* or NEB Turbo *E. coli* were used for most cloning procedures. To express Venus-tagged proteins in Kc167 cells or Flp-IN T-REx 293 cells, mini-Ph, full length Ph or PHC3 variants were first cloned into a house-modified Gateway donor vector^31^ using NEBuilder HiFi DNA Assembly cloning (NEB) or restriction-ligation cloning. Donor vector sequences were confirmed before Gateway LR recombination (LR Clonase II, Thermo Fisher) was used either with house-modified Gateway destination vector pMTVBH (harboring metallothionein promoter, pMT, for copper-inducible transgene expression in *Drosophila* cells, N-terminal Venus tag, Blasticidin resistance gene and H2Av-RFP under control of the constitutive Ac5 promoter) or with a modified vector pDEST-VF (harboring CMV promoter with tetO sites for Doxycycline-inducible transgene expression in Flp-IN T-REx 293 cells, N-terminal Venus-FLAG tag and FRT site for stable integration). Additionally, pMTVBH was also used to generate a Blasticidin-selectable control vector (pAc5-H2Av-RFP-Blast) with H2Av-RFP but without pMT-Venus cassette using HiFi Assembly cloning. For baculovirus-mediated expression in Sf9 cells, mini-Ph variants with WT SAM with an N-terminal 1X FLAG tag (DYKDDDDK) were cloned into a modified pFastBac1 using HiFi Assembly cloning and then used for transposition into DH10Bac *E. coli* (Thermo Fisher Scientific) to generate bacmids. For expression of mini-Ph variants with the EH mutant SAM in *E. coli*, mini-Ph^EH^ in a modified pET-3c vector was modified to replace PhL with PHC3L using HiFi Assembly cloning to generate mini-Ph^PHC3L-EH^, while mini-Ph^SAMsurf-EH^ was assembled using restriction-ligation cloning. Linker SAM variants with N terminal HisSUMO-tag for expression in *E. coli* were assembled in a pET vector (starting from Addgene plasmid #111559) using HiFi Assembly cloning. For AUC experiments, the DNA sequence matching Ph residues 1397 – 1577 comprising the Ph linker + SAM was cloned into a modified pET-3c plasmid following an N-terminal leader sequence of MHHHHHHAMKGVDSPSAELDKKAENLYFQGTR. Mutations were introduced using the QuickChange Lightning Site-Directed Mutagenesis Kit (Agilent). For CRISPaint editing of Kc167 cells, the *ph* targeting guide-RNA (CGTCGAGTATCAGTAATGGA) was introduced into pAc-sgRNA-Cas9 (Addgene #49330) using restriction-ligation cloning. The template pCRISPaint TagRFP with T2A-puromycin N-acetyltransferase (Addgene #80971) was modified by replacing TagRFP with Venus followed by diglycine linker and the DNA sequence matching Ph-p residues 1288-1577 or mutated variants (PHC3L and SAMsurf) to generate pCRISPaint Ph internal Venus (iV) templates. N-terminally Venus-tagged *ph-p* transgenes were subcloned into pUASt-attB vector (a gift from David Hipfner lab) using restriction-ligation cloning for PhiC31 mediated generation of transgenic *Drosophila*. Sequences were confirmed using Sanger sequencing or Whole Plasmid Sequencing (Plasmidsaurus). Protein sequences were compiled in **Table S7**.

### Protein purification

Mini-Ph variants with WT SAM were purified from whole cell extracts of 1L of Sf9 cells infected with baculoviruses for 3 days essentially as described^114^ with modifications as indicated. Briefly, the following steps were performed at 4°C if not otherwise indicated. Cells were pelleted and washed once with 1X PBS with 0.2mM PMSF and resuspended in 2-3 volumes of Buffer F (20 mM Tris pH 8, 500 mM NaCl, 20% glycerol, 10 mM MgCl_2_, 0.4 mM EDTA, 2 mM DTT, 0.1% NP 40, 10 µM ZnCl_2_) with protease inhibitors (0.4 mM PMSF, 10 μg/ml Aprotinin, 10 μg/ml Leupeptin,2 μg/ml Pepstatin, 50 μg/ml N-α-tosyl-L-lysine chloromethyl ketone hydrochloride (TLCK), 1.6 μg/ml Benzamidine and 10 μg/ml phenanthroline). Cells were incubated on ice and homogenized with 3*10 strokes in a Dounce homogenizer over the course of 30 min followed by centrifugation for 20 min at 48,400*g. Nucleic acids were depleted from the supernatant by adjusting to a final concentration of 0.1% polyethyleneimine (from a 5% stock at pH 8) followed by immediate centrifugation for 10 min at 15,000*g. The supernatant was incubated overnight with M2 anti-FLAG agarose beads (Sigma-Aldrich) pre-equilibrated with Buffer F, and then washed by gravity flow (20 bead volumes per wash step) with increasingly stringent BC buffers (20 mM HEPES pH 7.9, 20% glycerol, 10 mM MgCl_2_ 0.4 mM EDTA, 1 mM DTT, 0.05% NP 40, 10 µM ZnCl_2_) containing 300 mM KCl (BC300N), 600 mM KCl and 1200 mM KCl with protease inhibitors (same for all wash buffers, 0.2 mM PMSF, 4 μg/ml Aprotinin, 4 μg/m Leupeptin l, 0.8 μg/ml Pepstatin, 20 μg/ml TLCK, 0.64 μg/ml Benzamidine and 4 μg/ml phenanthroline). After a stringent wash consisting of BC2000N + 1 M Urea (20 mM Hepes, pH 7.9, 0.4 mM EDTA, 2 M KCl, 1 M deionized urea, 0.05% NP40, no glycerol), the salt concentration was reduced to 300 mM KCl using a descending series of BC buffers. anti-FLAG beads were incubated with 5 volumes of BC300N with 4 mM ATP (Millipore Sigma, A3377) + 4 mM MgCl_2_ for 30 min at room temperature to reduce the amount of HSC-70 that may otherwise copurify. Following washes with BC300N and BC300 (without NP40) without MgCl_2_, the protein was eluted with 0.1 M Glycine, pH 3, 0.5 M NaCl, 1 mM DTT, 10 µM ZnCl_2_. Collected elution fractions were immediately neutralized with 1 M Tris pH 8.5, 0.5 M NaCl (added at 1:20), and pooled protein containing fractions were dialyzed through three changes of BC300 (with 1 mM DTT, no protease inhibitors except 0.2 mM PMSF) before concentrating at room temperature using an Amicon 10 kDa cut off microconcentrator (Millipore) to 1-2 mg/ml and storing at –70°C. Aliquots of mini-Ph variants were buffer-exchanged into standard BC300 for *in vitro* phase separation assays using Zeba columns (Thermo Fisher Scientific) to remove DTT.

mini-Ph variants with the EH SAM mutation were expressed in BL21 (DE3) Rosetta pLysS *E. coli* (for mini-Ph^EH^ and mini-Ph^SAMsurf-EH^) or BL21(DE3) LOBSTR-RIL *E. coli* (mini-Ph^PHC3L-EH^) following previously published conditions with modifications as indicated^3^. mini-Ph^EH^ cultures were grown at 37°C to an OD of 0.8–1.0 and then shifted to 15°C for overnight induction with 1 mM IPTG. Cells were pelleted, flash-frozen, and stored at –70°C. For mini-Ph^EH^, cells were resuspended in 2 ml/g lysis buffer consisting of 50 mM Tris, pH 8.5, 200 mM NaCl, 10 mM β-ME, 100 µM ZnCl_2_, 0.2 mM PMSF, and 0.5 mM benzamidine. After incubation on ice for 10 min, cells were flash-frozen in liquid nitrogen, thawed at 37°C, and sonicated 6*30 s at 30% intensity. Freeze–thaw and sonication were repeated, and the lysate was cleared by centrifugation at 4°C. The cleared lysate was sonicated again for 6*30″ at 40% intensity and filtered through a 22-µm filter. Lysate from 1 l culture was applied to a 1-ml His-Trap column (Cytiva) using an ÄKTA FPLC system and eluted with an imidazole gradient from 10 to 300 mM in lysis buffer. Fractions with mini-Ph^EH^ were dialyzed overnight against 1 l of 20 mM Tris, pH 8.5, 50 mM NaCl, 100 µM ZnCl_2_, and 10 mM β-ME. Dialyzed fractions were centrifuged at 4°C to remove potential aggregates, and the supernatant was loaded on a 1 ml HiTrapQ-HP column (Cytiva) and eluted with a gradient from 50 mM to 1 M NaCl in lysis buffer. Fractions containing mini-Ph^EH^ were pooled and dialyzed overnight into 20 mM Tris, pH 8, 50 mM NaCl, 10 µM ZnCl_2_, and 1 mM β-ME, aliquoted, and stored at –70°C. The purification for mini-Ph^PHC3L-EH^ had the following modifications. Cultures were grown at 37°C to OD 0.3, and then shifted to 18°C before overnight induction with 1 mM IPTG was started at OD 0.8. Lysis buffer was 50 mM Tris, pH 8.5, 200 mM NaCl, 5 mM β-ME, 0.2 mM PMSF. The cleared lysate was filtered but not sonicated. The lysate was applied to a 5 ml His-Trap column (Cytiva) and protein was eluted with an imidazole gradient from 0 to 500 mM. Dialyzed fractions (in 50 mM Tris, pH 8, 200 mM NaCl, 5 mM β-ME, 0.2 mM PMSF) were loaded on a 5 ml HiTrapQ-HP column (Cytiva). The column was washed with 15 ml of buffer A (50 mM Tris, pH 8, 50 mM NaCl, 5 mM β-ME, 0.2 mM PMSF) and elution was performed with a stepwise gradient of buffer A with NaCl as follows: 50-170 mM NaCl for 10 ml, 170 mM NaCl for 15 ml, 170-300 mM NaCl for 10 ml, 300 mM NaCl for 20 ml (main elution peak observed here), 300-420 mM NaCl for 10 ml, 420 mM NaCl for 15 ml, 420-540 mM NaCl for 10 ml, 540 mM NaCl for 15 ml, 540-1000 mM NaCl for 20 ml and 1000 mM NaCl for 20 ml. The pooled fractions containing protein were concentrated and size-exclusion chromatography using a Superdex 200 10/300 GL column (Cytiva) was performed in 50 mM Tris, pH 8.5, 500 mM NaCl, 5 mM β-ME, 0.2 mM PMSF. The final protein fractions were pooled, concentrated and aliquoted, before being stored at –70°C. The purification for mini-Ph^SAMsurf-EH^ had the following modifications. Cultures were grown at 37°C to OD 0.3, and then shifted to 15°C before overnight induction with 1 mM IPTG was started at OD 0.8. Lysis buffer A1 was 50 mM Tris, pH 8.0, 200 mM NaCl, 0.2 mM PMSF, 5 mM β-ME, 30 mM imidazole. Cells were sonicated 5*1 min (4 s on/4 s off pulses) at 50% amplitude. Cleared lysate was added to pre-equilibrated Ni-NTA beads in bulk and rotated at room temperature for 1 h. After centrifugation, the Ni-NTA binding was repeated twice with fresh beads using the supernatant, while the beads with bound protein were washed twice with lysis buffer and then transferred to columns. The protein was eluted in a stepwise manner using 300 mM imidazole in buffer A1, while the protein concentration was followed using Bradford assay. Fractions with eluted protein from all batches were pooled and dialyzed through two changes of buffer A2 (50 mM Tris, pH 7.0, 0.1 mM PMSF, 5 mM β-ME). The dialyzed sample, which was slightly hazy, was clarified by centrifugation and loaded on a 5 ml HiTrap SP column (Cytiva) pre-equilibrated with buffer A2. The column was then washed with 15 ml buffer A2 after loading, and elution was performed with a stepwise gradient of buffer A2 with NaCl as follows: 0-130 mM NaCl for 20 ml, 130 mM NaCl for 12 ml, then 130-250 mM NaCl for 15 ml and 250 mM NaCl for 8 ml, then 250-330 mM NaCl for 8 ml and 330 mM NaCl for 20 ml (main elution peak observed here), then 330-500 mM NaCl for 20 ml and 1 M NaCl for 14 ml. Pooled protein containing fractions were concentrated and loaded on a Superdex 75 26/600 column (Cytiva) for size-exclusion chromatography in buffer A3 (50 mM Tris, pH 8.0, 200 mM NaCl, 0.1 mM PMSF, 5 mM β-ME). Since the chromatogram indicated heterogenous oligomeric states, pooled fractions with protein were subsequently incubated overnight at 4°C in buffer A3 with the addition of high salt (∼867 mM NaCl final) and β-ME (15 mM final). The sample was then concentrated and loaded again on a Superdex 75 26/600 column for size-exclusion chromatography in buffer A4 (50 mM Tris, pH 8.0, 500 mM NaCl, 0.1 mM PMSF, 5 mM β-ME) giving rise to a more homogenous chromatogram. The high salt, high β-ME (1 M NaCl, 45 mM β-ME final) overnight incubation was repeated on the concentrated pooled fractions from the second size-exclusion chromatography, before a final size-exclusion chromatography in buffer A4 was performed. The pooled protein containing fractions from this run were concentrated and stored at –70°C. Aliquots of mini-Ph EH variants were buffer-exchanged into BC300 for *in vitro* assays using Zeba columns, except for titration experiments (details below).

HisSUMO-tagged Linker-SAM proteins were expressed in BL21 (DE3) LOBSTR-RIL cells. Cultures were grown at 37°C to an OD of 0.6 and then shifted to 18°C for overnight induction with 1 mM IPTG. Cells were pelleted, washed once in ice cold PBS with 1 mM PMSF, flash-frozen, and stored at −70 °C. Cells were resuspended in 2 ml/g lysis buffer (50 mM Tris, pH 8.5, 500 mM NaCl, 10 mM β-ME, 1 mM PMSF, 1 mM EDTA). 1 mg/ml lysozyme was added, and cells were incubated for 1 h at 4°C on a nutator. Afterwards, cells were flash-frozen in liquid nitrogen, thawed at 37°C, and sonicated in an ethanol ice bath for 5×30s (30 s off cycle) at 40% intensity. Freeze–thaw and sonication were repeated three times, and the lysate was centrifuged for 1 h at 48400*g and 4°C. Cleared lysate was filtered through a 0.2-µm filter. Filtered lysate was applied to a 1 ml His-Trap column using an ÄKTA FPLC system and eluted with a gradient of imidazole (from 20 mM to 1 M) in binding buffer (50 mM Tris, pH 8.5, 500 mM NaCl for PHC3L-SAM/1 M for PhL-SAM and PhL-SAMsurf, 10 mM β-ME, 1 mM PMSF). Fractions with recombinant protein were pooled and dialyzed overnight against binding buffer. The eluted fractions were further purified by size-exclusion chromatography using a Superdex 200 10/300 GL column, then concentrated to at least 1 mg/ml, flash frozen and stored at −70 C.

For cleaving HisSUMO-tag, His-tagged SUMO protease Ulp1 (pFGET19-Ulp1, Addgene #64697) was expressed in BL21 (DE3) Rosetta cells. Cultures were grown at 37°C to an OD of 0.5 and protein expression was induced with 1 mM IPTG for 3 h. Cells were pelleted, resuspended in 4 ml/g lysis buffer (50 mM NaH_2_PO_4_, 300 mM NaCl, 10 mM imidazole, pH 7.5). 1 mg/ml lysozyme, 1 mM MgCl_2_ and Benzonase (Sigma-Aldrich) were added, and cells were incubated for 30 min on ice. Cells were sonicated for 10×30 s (30 s OFF cycle) in an ethanol ice bath at 40% intensity. The lysate was cleared for 25 min at 48,400*g and 4°C, before it was applied on a Ni NTA agarose beads equilibrated in lysis buffer. Lysate was passed over the beads by gravity flow four times, then washed with 10 bead volumes wash buffer (50 mM NaH_2_PO_4_, 300 mM NaCl, 20 mM imidazole, pH 7.5) before the protein was eluted with 3 volumes elution buffer (50 mM NaH_2_PO_4_, 300 mM NaCl, 250 mM imidazole, pH 7.5). Fractions containing Ulp1 were pooled, flash frozen and stored at −70°C.

For proteins used for AUC analysis, the DNAs were transformed into BL21 Gold (DE3) pre-transformed with pLysS RARE2 (Agilent) and induced overnight at 15°C using 1 mM IPTG. Bacterial cells collected from 1 l of culture were resuspended with 10 ml of lysis buffer (50 mM Tris pH 8.0, 150 – 200 mM NaCl, 20 mM imidazole, 5 mM β-ME, 1 mM PMSF). For the D16K D17R D33K D34R (SAMsurf) protein with the oligomer interface (EH) mutant, 20 mM ADA pH 6.0, 50 mM NaCl, 1 mM β-ME was used as the lysis buffer. Cells were lysed via sonication and the soluble lysate introduced to 1 ml of bulk Ni-sepharose beads. The beads were exposed to the soluble lysate with gentle agitation for 30 – 60 min at room temperature, then washed several times with the lysis buffer (without PMSF) using a total volume 30 – 50X the Ni-sepharose volume. The bound proteins were eluted with 300 mM imidazole pH 6.9 – 7.5, 150 mM NaCl, 5 mM β-ME. For the SAMsurf EH mutant, the elution buffer was 20 mM ADA pH 6.0, 50 mM NaCl, 1 mM β-ME. After the bulk Ni-affinity purification, all proteins except the two having the SAMsurf mutations were further purified using anion exchange chromatography (HiTrap Q) running a gradient from 0% (i.e. 25 mM Tris pH 8.0 – 8.5, 50 mM NaCl, 5 mM β-ME) to 100% of 25 mM Tris pH 8.0, 1 M NaCl, 5 mM β-ME over 60 min. Cation exchange chromatography was used for the SAMsurf proteins, either with or without the oligomerization interface mutation, using a gradient of 0% (i.e. 20 mM MES or ADA pH 6.0, 50 mM NaCl, 1 mM β-ME) to 100% 20 mM Tris pH 8.0, 1 M NaCl, 1 mM β-ME over 60 min. The SAMsurf protein with the oligomer interface mutant was further purified over a HisTrap HP column using a gradient of 0% 20 mM Tris pH 8.0, 200 mM NaCl, 5 mM β-ME to 100% 500 mM imidazole pH 7.5, 200 mM NaCl, 5 mM β-ME over 60 min. For the AUC experiments, all proteins were diluted to 37 µM into a buffer of 10 mM Tris pH 8.0, 50 mM NaCl, 1 mM TCEP.

### Test for DNA contamination in protein preparations

Protein preparations used in phase separation assays were tested for contamination with nucleic acid. Protein samples were digested with Proteinase K (BioBasic) overnight at 37°C. The samples were loaded on a 1% agarose gel and run for 30 min at 100 V. The gel was stained with highly sensitive SYBR Gold dye (Thermo Fisher Scientific) before imaging on a Typhoon imager (GE Healthcare).

### Size-exclusion chromatography with multi-angle light scattering (SEC-MALS) analysis

Size exclusion chromatography (SEC) was performed using a Superdex 200 10/300 GL column pre-equilibrated with protein storage buffer BC300. 60 to 384 µL samples of mini-Ph variants were injected into the column (to reach an equal loading of 500 µg per sample). For mini-Ph EH SAM variants buffer exchange into BC300 was performed using Zeba columns prior to injection (not for other mini-Ph variants). Chromatographic separation was carried out at room temperature using an ÄKTAmicro chromatography system (GE Healthcare) at a flow rate of 0.5 ml/min. The column effluent was analyzed using an inline DAWN HELEOS II multi-angle light scattering (MALS) detector and an Optilab T-rEX refractive index (RI) detector (Wyatt Technology), enabling simultaneous measurements of UV absorbance, light scattering, and refractive index. Data were processed using ASTRA software (v6.1.6.5, Wyatt Technology) to calculate the molecular weight of the proteins. The system was calibrated using a bovine serum albumin (BSA) standard.

### Mass photometry analysis

Mass photometry (MP) data were measured from mini-PH variant samples (WT, PHC3L, SAMsurf, EH, PHC3L-EH, SAMsurf-EH) using a Refeyn OneMP instrument at the New York Structural Biology Center. The instrument light source was allowed to warm up for the recommended 60 min to enable precise molecular mass analyses. The glass slides (Refeyn) used in the measurements were cleaned with 100% isopropyl alcohol followed by MilliQ Water (Millipore) rinse and placed into six-well cassettes (Refeyn) for measurements. Autofocus on the sample drops was established using the manufacturer’s recommendation. Once autofocus and zero signal intensity are established, differences in laser signal intensity due to the presence of protein are read as contrast values. MP data were taken over a 1 min period, followed by a second 1 min read for a lower incident count using the instrument’s default settings. A standard curve generated with apoferritin at different concentrations was used to convert contrast values into kDa values and fit them to a Gaussian curve. MP data were taken with the Refeyn AcquireMP and analyzed with DiscoverMP.

Mini-PH proteins concentrations were measured using a NanoDrop spectrophotometer (Thermofisher) at 280 nm (extinction coefficients are listed in **Table S7**). Protein samples were diluted to 1000 nM using BC300 storage buffer without glycerol (20 mM HEPES, pH 7.9, 300 mM KCl, 0.2 mM EDTA, pH 8) before MP data collection. The instrument was blanked against a 10 µL drop of BC300 storage buffer, 0.22 μm filtered (Millipore). No contaminants (typically dust) were observed in the storage buffer blanks. Mini-PH proteins were added to the well containing the buffer blank to concentrations of 500 nM, 250 nM, and 125 nM. Sample reads were performed in storage buffer and compared against each other for protein oligomerization.

### Fluorescent labelling of mini-Ph variants

NHS-ester-Cy3 (Cytiva) was used to randomly label lysines in mini-Ph proteins essentially as described^3^ with the exception that proteins were not buffer exchanged before labeling. Labeling was carried out at a ratio of 0.5:1 (dye:protein) for 1 h at room temperature in protein storage buffer) and quenched by addition of a lysine solution to 10 mM. Free dye was removed using a Zeba column pre-equilibrated with protein storage buffer. For imaging experiments, labeled and unlabeled protein was mixed at ratios between 1:5 and 1:10 (labelled:unlabelled), depending on the labeling efficiency.

### Phase separation assays

Proteins and DNA templates (“TPT”^115^, prepared by PCR followed by extraction from acrylamide gels, **Table S6**) were routinely centrifuged full speed in a microfuge for 2–5 min at 4°C to remove aggregates before setting up phase-separation assays. Reactions (typically 15 µl) were assembled in a 384-well glass-bottom imaging plate (Greiner Bio-One). Wells were pre-treated following a previously published protocol^116^. Briefly, the imaging plate was cleaned by submerging in 1 l of 5% Hellmanex III (Sigma-Aldrich) solution for 1 h, followed by extensive washes with tap water and milliQ H_2_O. Each well was then etched by adding 1 M KOH and incubating for 1h before extensive washes with tap water and milliQ were performed. The wells were then treated with Sigmacote (Sigma-Aldrich), washed twice with isopropanol and the dried plate was sealed with adhesive plastic film. Just before setting up phase separation reactions, wells were coated by adding 0.5% (w/v) Pluronic F127 (Sigma-Aldrich) in BC0 buffer without glycerol (20 mM HEPES, pH 7.9, 0.4 mM EDTA) and incubating for at least 15 min. Wells were then washed 5 times with salt-free phase separation buffer (either BC0 without glycerol, or 50 mM Tris, pH 8 for mini-Ph^EH^ titrations) without draining wells completely. After the last wash, each well was quickly drained completely and 10 µl buffer were added (final BC0 was supplemented with 25 µM ZnCl_2_). YOYO-1 (Thermo Fisher Scientific, 100 nM final concentration) and DNA (132 nM, if any) were added before phase separation reactions were initiated by addition of the protein and mixing the reaction by gently pipetting up and down three times without introducing air. Standard reaction conditions were 20 mM HEPES, pH 7.9, 60 mM KCl, 0.4 mM EDTA, 4% glycerol, 20 µM ZnCl_2_, except for EH titrations, which had 50 mM Tris, pH 8, 50 mM NaCl, 20 µM ZnCl_2_. Reactions without labelled protein and YOYO-1, and reactions to test reversibility of condensate formation were set up under standard reaction conditions with 4 µM protein. While titrations were imaged after overnight incubation in the dark, unlabelled reactions were incubated for approximately 1 h before imaging. Reversibility tests were performed by spiking in 3 µl of high salt BC1500 buffer without glycerol to the 15 µl reaction after ∼30 min to reach 300 mM KCl (same salt concentration as under storage conditions). The final conditions for the reversibility assays were thus 17 mM HEPES, pH 7.9, 300 mM KCl, 0.33 mM EDTA, 3.3% glycerol, 17 µM ZnCl_2_ with 3.3 µM protein and 110 nM DNA (if any). Condensates immediately began to dissolve, and wells were imaged after ∼15 min.

Phase separation reactions of HisSUMO-Ph linker SAM or HisSUMO-PHC3 linker-SAM (diluted to NaCl concentration of 300 mM) in untreated glass-bottom wells were initiated by addition of 1:10 molar ratio of Ulp1 diluted in 50 mM Tris, pH 8 resulting in cleavage of the solubility tag as assessed with SDS PAGE. Ulp1 was omitted for reactions with uncleaved HisSUMO-linker-SAM proteins. Reaction conditions were 60 mM NaCl, 50 mM Tris, pH 8.5 and 6 µM protein were used.

For FRAP experiments, BC300-buffer exchanged mini-Ph samples were used and phase separation reactions were set up in coated glass-bottom wells as described above with protein/DNA concentrations of 4 µM and 132 nM (if any), respectively. The reactions were supplemented with an oxygen scavenger system composed of DTT, glucose, Glucose Oxidase and Catalase (as described in^117^). Final reaction conditions were 20 mM HEPES, pH 7.9, 60 mM KCl, 0.4 mM EDTA, 4% glycerol, 20 µM ZnCl_2_, 10 mM DTT, 40 mM glucose, 20 µg/ml Glucose Oxidase (Sigma-Aldrich) and 3.5 µg/ml Catalase (Sigma-Aldrich). In the absence of oxygen scavengers, we did not observe protein recovery, and DNA recovery was variable, consistent with photodamage from the bleaching step. FRAP imaging typically started within the first 20 min of condensate formation (for condensates to be big enough for bleaching) and data acquisition was stopped after ∼1 h, when condensate dynamics started to change.

### Electrophoretic mobility shift assay (EMSA)

HisSUMO-Linker-SAM proteins were diluted to appropriate concentrations for the titration series in 50 mM Tris pH 8.5 buffer and the salt concentration was adjusted to 300 mM NaCl for all samples. DNA binding reactions with 15 nM dsDNA were started by addition of protein to reactions. The final composition of the reaction buffer was 50 mM Tris pH 8.5, 60 mM KCl, 5 mM DTT in 10 µl (8 µl reaction mix plus 2 µl diluted protein), and reactions were incubated at room temperature for 1 h. In the meantime, a 5% native polyacrylamide gel prepared with 0.5X TBE buffer was pre-run at 4°C for 15 min. After the incubation of DNA binding reactions was completed, 2 µl 50% glycerol were added and 9 µl of each sample were loaded onto the gel, which was run for 1 h at 4°C. The gel was then stained with SYBR Gold and imaged on a Typhoon imager.

### Analytical ultracentrifugation

Sedimentation velocity experiments were performed on a Beckman Coulter Optima AUC at the Canadian Center for Hydrodynamics at the University of Lethbridge. All samples were measured at 37 µM in 10 mM Tris, pH 8.0, 50 mM NaCl, and 1 mM TCEP using 2-channel epon-charcoal centerpieces fitted with quartz windows. Data was collected in intensity mode at 280 nm for all samples except Ph 1397-1577 SAMsurf L1565R (EH mutant), which was measured at a higher concentration using 296 nm detection, so that the absorbance was within the dynamic range of the detector. Despite the elevated concentration of L1565R mutant used in the experiment, no oligomerization was detected, indicating the mutation prevents all polymerization. All data were analyzed with UltraScan III version 4.0^118^ and fitted with an iterative two-dimensional spectrum analysis^119^ to fit the meniscus position and time– and radially-invariant noises. Diffusion-corrected integral sedimentation coefficient distributions were generated using the enhanced van Holde-Weischet analysis methods^120^. UltraScan calculated the buffer density and viscosity to be 1.000580 g/cm and 1.00536 cP, respectively.

### Cell culture, transfection and generation of cell lines

For imaging of transient transfections, 1 million cells were plated in 24-well plates the night before transfection. Transfection of pMTVBH constructs with copper-inducible Venus tagged (mini-)Ph transgenes and H2Av-RFP nuclear was carried out using TransIT-Insect Transfection Reagent (Mirus), according to the manufacturer’s protocol. The day after the transfection, transgene expression was induced by addition of 300 mM CuSO4. On the second day after transfection, cells were replated in CuSO_4_-containing media on ConA-coated 96-well imaging plates (Ibidi). For the ConA coating, the glass surface was treated by adding 0.5 mg/ml ConA (Medicago) in sterile water and incubating for 1 h at room temperature, which was followed by rinsing several times with milliQ H_2_0 and a final wash with 100% ethanol before drying the glass (all steps performed under sterile conditions). Cells were imaged on the third day after two days of transgene induction (details below). Remaining cells from the 24-well plate were harvested for Western Blot by spinning them down and washing 1X in PBS. Pellets were stored at 70°C.

To generate stable cell lines with copper-inducible mini-Ph expression, Kc167 cells were transfected with pMTVBH constructs as above. A control cell line with H2Av-RFP but without Ph transgene was generated similarly using pAc5-H2Av-RFP-Blast. Three days after transfection, 6 µg/ml Blasticidin were added. Selection was maintained for three weeks by regular addition of fresh Blasticidin-containing media. Cells with integrated transgenes were isolated by FACS based on H2Av-RFP expression onto irradiated wild type Kc167 cells (24 kR gamma irradiation as described^121^) in 6-well plates. Once cells had recovered and started to grow normally, they were maintained in standard M3 media with 6 µg/ml Blasticidin. For live imaging, cells were analyzed on ConA-coated 8-well imaging slides (Ibidi) after two days of induction in media with 300 mM CuSO_4_ but without Blasticidin. Western Blot samples from these cells were collected as described above.

For generating endogenous edits of *ph*, we used the previously published CRISPaint approach^58,59^. Kc167 cells were transfected with pAc-sgRNA-Cas9 targeting all *ph* alleles just upstream of the mini-Ph region, the pCRISPaint Ph iV templates and pCFD3-Frame selector 1 (Addgene #127554) targeting the template. This strategy allowed us to insert an internal Venus tag into *ph* alleles along with mutations in the mini-Ph region, effectively replacing the endogenous sequence with the sequences in pCRISPaint. Three days after transfection, 5 µg/ml puromycin (Wisent) was added to the cells to start selection. Only those cells in which *ph* alleles were repaired in frame by integrating the pCRISPaint template can express the T2A-puromycin N-acetyltransferase, thus being resistant to puromycin. Selection was maintained for three weeks by regular addition of fresh puromycin containing medium. FACS was used to isolate Venus expressing cells, which were sorted onto 96-well plates containing standard media with irradiated wild-type Kc167 cells. Once colonies had formed, cells were expanded for confirmation of correct edits. We screened cells by microscopy to confirm condensate formation in the nucleus and by PCR across the expected junction. To genotype the clones, we prepared genomic DNA by standard Proteinase K digest and phenol-chloroform extraction followed by ethanol precipitation. We carried out PCR with primers specific to ph-p (616-2012) and ph-d (616-2013) to assess alleles without Venus, and with primers that amplify either ph-p or ph-d (2057-2011)—Venus fusions (see **Table S6** for primer sequences) using KOD polymerase (Millipore Sigma). PCR products were purified using PCR cleanup columns (Macherey-Nagel) and then through a house-made G-25 or G-50 Sephadex spin column equilibrated in H_2_O followed by sequencing at Plasmidsaurus (“premium PCR”). To confirm that multiple targeting events, if present, could be detected, we sequenced a mixture of DNA amplified from ph-p—Venus and ph-d—Venus alleles. The results from genotyping were compiled in **Table S3**. For live imaging, cells were first transfected with pAc5-H2Av-RFP-Blast and maintained in 6 µg/ml Blasticidin-containing media for up to three weeks. Transfected cells with nuclear H2Av-RFP marker were plated and grown on ConA-coated 8-well imaging slides for at least two days in the absence of Blasticidin before imaging.

For generation of cell lines with doxycycline-inducible Ph/PHC3 expression, Flp-IN T-REx 293 (in Zeocin-free media) were co-transfected with pDEST-VF constructs and pOG44 using TransIT-293 Transfection Reagent (Mirus). pOG44 encodes Flp recombinase, which allows integration of pDEST-VF at the genomic FRT site of the Flp-IN cells. On the third day after transfection, selection of cells with pDEST-VF was started by addition of media with 200 µg/ml Hygromycin and 6 µg/ml Blasticidin. The selection was maintained for around three weeks by regularly replacing media with fresh Hygromycin and Blasticidin-containing media. During this time, colonies started to form, indicating successful integration of inducible Ph/PHC3 transgenes. For live imaging, cells were plated on Poly-L-lysine-coated 8-well imaging slides. For the Poly-L-lysine coating, the glass surface was treated like for ConA coating but by adding 0.1% (w/v) aqueous Poly-L-lysine solution (Electron Microscopy Sciences). Protein expression was induced for one day in standard media with 1 µg/ml Doxycycline but without selection antibiotics before image acquisition, for which nuclei were stained by addition of DRAQ5 (Thermo Fisher Scientific) to a final concentration of 2.5 µM (from a 125 µM dilution in PBS). Western Blot samples were collected as above.

### DAPI staining for nuclear size measurement

CRISPaint cell lines were plated on a ConA-coated 12-well imaging slides (Ibidi) and fixed the next day by adding 4% formaldehyde diluted in PBS for 10 min. Cells were washed once with PBS, before being permeabilized in PBS with 1% triton-X for 15 min. After another wash with PBS, nuclei were stained by incubating with 0.5 µg/ml DAPI (Sigma Aldrich) in PBS for 15 min. Following another wash with PBS and addition of mounting media (20 mM Tris, pH 8, 0.5% (w/v) N-propyl gallate [Fluka], 90% (v/v) glycerol), the slide was covered with a coverslip and sealed with nail polish. Slides were stored at 4°C until imaging with a spinning disc microscope.

### Growth assay

250,000 cells were plated in 1 ml media in 24-well plates. For counting each day over the next six days, cells were stained with trypan blue (no excessive cell death was observed) and mounted on a hemocytometer. Brightfield images were taken on a DM6 microscope (Leica) and live cells were counted manually from images. The data of three replicates was fit with a exponential growth equation in Graphpad Prism 10 (v10.3.0, Dotmatics) to extract the doubling time, which was statistically compared using Kruskal-Wallis test followed by Dunn’s test for multiple comparison (p values are from multiple comparison test).

### Hoechst cell cycle staining

4 million cells in standard M3 media were fixed with formaldehyde at a final concentration of 1% on a nutator at room temperature for 10 min. Fixation was quenched by addition of glycine pH 7.9 to 130 mM and cells were kept on ice. Cells were washed once in PBS, resuspended in permeabilization buffer (0.015% Triton X-100 in PBS) and incubated on ice for 15 min. Cells were washed once with FACS Wash Buffer (1% BSA [BioShop], 0.1% Triton 100X in PBS) and DNA stained with 1 µg/ml Hoechst for 45 min on ice in the dark. Cells were washed twice with FACS wash buffer and resuspended in PBS and kept on ice. Cell cycle profiles were analyzed using a LSRFortessa flow cytometer (BD Biosciences) equipped with 405 nm laser. Data were analyzed in FlowJo (v10.7.2, BD) and cell cycle profiles were fitted with the built-in functions.

### RNA extraction and reverse transcription (RT) reactions

RNA was extracted from ∼20 million cells using TRIzol Reagent (Thermo Fisher Scientific) and resuspended in RNAse-free water (Wisent) and quantified on a NanoDrop spectrophotometer. 10 µg RNA were treated with DNAseI (NEB) for 15 min at 37°C. DNAse-treated RNA was then purified using the RNA Clean & Concentrator-5 Kit (Zymo Research) and eluted in RNAse-free water. After NanoDrop quantification, 2 µg RNA were used for reverse transcription (RT) reactions using SuperScript IV Reverse Transcriptase (Thermo Fisher Scientific). Reactions were performed according to manufacturer instructions with fresh dNTP aliquots (NEB), random hexamer primers (IDT) and murine RNAse inhibitor (NEB). Controls (no reverse transcription) were carried out for each cell line to verify absence of genomic DNA contamination (specifically for non-exon spanning primers). The cDNA was stored at –20°C before qPCR analysis.

### Chromatin immunoprecipitation (ChIP)

For each cell line, 20 million cells were fixed in media by adding formaldehyde to a final concentration of 1% for 10 min on a nutator at room temperature. The fixation was quenched by addition of glycine pH 7.9 to a final concentration of 130 mM and incubation for 5 min on a nutator at room temperature before transferring to ice. Cells were collected by centrifugation for 4 min at 1300*g and 4°C (this centrifugation was used between all subsequent washes), washed 1X with PBS with 0.3 mM glycine, pH 7.9 and 1X with PBS. Pellets were resuspended in wash buffer I (10 mM Hepes, pH 7.6, 10 mM EDTA, 0.5 mM EGTA and 0.25% (v/v) Triton-X100) and incubated 10 min at 4°C on a nutator, centrifuged and the pellet resuspended in wash buffer II (10 mM Hepes, pH 7.6, 200 mM NaCl, 1 mM EDTA, 0.5 mM EGTA, 0.01% Triton-X-100) and incubated as above followed by centrifugation. Pellets were resuspended in PBS with protease inhibitor mix (0.2 mM PMSF, 10 μg/ml aprotinin, 10 μg/ml Leupeptin, 2 μg/ml Pepstatin, 50 μg/ml TLCK, 1.6 μg/ml Benzamidine and 10 μg/ml phenanthroline, 1 mM DTT), centrifuged, and the pellet was frozen in liquid nitrogen and stored at –70°C before further processing. The samples were resuspended in 1 ml of sonication buffer (50 mM Hepes, pH 7.5, 500 mM NaCl, 1 mM EDTA, 1% Triton X-100, 0.1% sodium deoxycholate, 0.1% SDS), incubated for 10 min on ice and sonicated in a 15 ml polypropylene tube in an ice-water-ethanol bath with 8*30 s pulses (with 59 s between pulses) at an amplitude of 40%. After centrifugation, 100 µl supernatant for each cell line were kept aside as input. SDS was added to input samples to 1% final concentration, after which samples were incubated at 65°C overnight to reverse crosslinking. RNAse A (BioShop) was added to 180 µg/ml followed by incubation for 30 min at 37°C. The buffer was then supplemented with 34 mM Tris, pH 7, 8 mM EDTA and 170 µg/ml Proteinase K and samples were incubated at 50°C for 1 h. Chromatin fragments from input samples were purified using NucleoSpin Gel and PCR Clean-up Kit (Macherey-Nagel) with NTB buffer (Macherey-Nagel) for SDS-containing samples. For immunoprecipitation of the crosslinked chromatin samples after sonication, Protein G dynabeads (Thermo Fisher Scientific) were prepared by first washing twice with cold, filtered PBS containing 5 mg/ml BSA) on a magnetic rack. Beads were kept in PBS-BSA and antibodies were added followed by overnight incubation on a rotator at 4°C. The following antibodies and quantities were used: rabbit anti-Ph (20 µl serum per sample, Francis lab, mouse anti-Psc (60 µl per sample, ∼4 µg antibody, DSHB), mouse anti-GFP (1:1 mix of two antibodies from DHSB, 50 µl each per sample, ∼4 µg antibody total) and control IgG (4 µl per sample, ∼4 µg antibody, Abcam). After the incubation, beads were washed twice with PBS-BSA. Chromatin samples (200 µl per ChIP) supplemented with 2.5X fresh protease inhibitors as above, without DTT, were diluted 1:2.5 in ChIP dilution buffer (16.7 mM Tris, pH 8, 167 mM NaCl, 1.2 mM EDTA, 1.1% Triton X-100 and 0.01% SDS) and NP40 was added to a final concentration of 0.1% (v/v). Pre-coupled protein G beads were added (equivalent of ∼4 µg of antibody per ChIP) and samples were incubated overnight at 4°C with rotation. The beads were then washed with the following series of buffers with 5 min incubation at 4°C on a rotator between washes: 1X with RIPA buffer (10 mM Tris, pH 8.0, 140 mM NaCl, 1 mM EDTA, 1% Triton-X100, 0.1% sodium deoxycholate, 0.1% SDS), 3X with RIPA containing 500 mM NaCl, 1X with LiCl wash (10 mM Tris, pH 8.0, 250 mM LiCl, 1 mM EDTA, 1% NP40 and 1% sodium deoxycholate) and 2X with TE buffer (10 mM Tris, pH 8.0, 1 mM EDTA). After the last wash, freshly prepared elution buffer (100 mM NaHCO_3_, 500 mM NaCl, 1%SDS) was added to the beads, and samples were incubated overnight at 65°C (which also reverses crosslinking). The next day, the supernatant was removed and another elution for 10 min at 65°C was performed. The combined eluates were then treated with RNAse A and Proteinase K, and chromatin fragments were purified in the same way as the input samples. DNA from input and ChIP samples was stored at 4°C before qPCR analysis.

### Quantitative PCR (qPCR)

qPCR was performed using a QuantStudio 5 system (Thermo Fisher Scientific) in a 384-well plate format. PowerUp SYBR Green Master Mix (Thermo Fisher Scientific) was used to set up 5 µl reactions with 120 nM primers. For RT-qPCR, cDNA was diluted to 5 ng/µl (or an equivalent for no RT control) and 1 µl was added per 5 µl reaction. The recommended PCR program for the utilized qPCR mix was used and threshold values (Ct) were exported using QuantStudio Design & Analysis software (v1.5.3, Thermo Fisher Scientific) for downstream ΔΔCt analysis. The resulting fold changes represent values normalized to α-Tubulin (ΔCt) and then normalized to one Ph WT replicate (ΔΔCt) to emphasize the inherent variations. The statistical comparisons were calculated using ANOVA followed by Tukey’s multiple comparison test (normality of data was verified with in-built functions; p-values are from multiple comparison test) and graphs were plotted with GraphPad Prism 10. For ChIP-qPCR, input samples were diluted 1:20 and ChIP samples were diluted 1:2 and 1 µl was added per 5 µl reaction. Here, the standard curve mode was used with the recommended PCR program using Kc167 genomic DNA as reference, and DNA amount per reaction was exported using the Design & Analyze software to quantify the percent of input DNA in the ChIP samples. The resulting data was statistically compared using Kruskal-Wallis test followed by Dunn’s multiple comparison test (the data was not consistently normally distributed as verified with in-built functions; p-values are from multiple comparison test) and graphs were plotted with GraphPad Prism 10.

### Transgenic *Drosophila* experiments

Venus-tagged *ph* transgenes subcloned into pUASt-attB vector were integrated into y[1] w[*] P{y[+t7.7]=nos-phiC31\int.NLS}X; P{y[+t7.7]=CaryP} attP2 flies by microinjection into early syncytial-stage blastoderm embryos (Genome ProLab, Sherbrooke, QC, Canada). This background contains both PhiC31 integrase on the X chromosome and an attP integration site on chromosome 3. Stable transformants were selected by integration of the *white* gene into *white* mutant background. Transgenic constructs of *ph* are controlled by the UAS promoter. Expression of Ph variants (or GFP-nls, a gift from David Hipfner lab) was driven by mating UAS transgenic flies with the *hedgehog*-Gal4/*tubulin*-Gal80ts driver line harboring H2Av-RFP as a nuclear marker for live imaging. Crosses were kept at 25°C for 5 days before shifting to 27°C for 24 h. Larvae were collected in the mid to late L3 stage and wing imaginal discs were dissected and mounted in M3 medium for live imaging. For imaging of whole wing discs, discs were mounted on 35 mm imaging dishes (Ibidi) with 0.12 mm thick adhesive spacers (a gift from David Hipfner lab, Sigma-Aldrich) to avoid damage to the disc when a coverslip was put on top to keep the tissue in place. Spacers were omitted for condensate imaging at high magnification.

### Imaging

All images from mini-Ph phase separation assays (except FRAP experiments) as well as z-stacks from live-cell imaging (except CRISPaint lines with endogenous edits), fixed cells for nuclear area measurements and wing discs, were collected on a Zeiss AxioObserver Z1 microscope, equipped with a Yokogawa CSU-1 spinning-disk confocal head and an Evolve EMCCD camera from Photometrics. Zen 2012 software (Zeiss) was used for image acquisition with a 63X oil objective for phase separation assays and cell imaging, 20X air objective for full wing discs. The excitation wavelengths for YOYO-1/Venus, Cy3/RFP and DRAQ5 were 488 nm, 561 nm, and 639 nm, respectively. Phase contrast images of mini-Ph condensates were also acquired on this system. For linker-SAM proteins, phase contrast images were collected on an LSM 710 AxioObserver Confocal Microscope (Zeiss) with a 40X oil objective.

FRAP imaging was conducted using an LSM980 NLO laser scanning confocal microscope (Zeiss) with a 63X oil objective and Definite Focus Z-stabilization mode. Excitation wavelengths for YOYO-1 and Cy3 were 488 nm and 543 nm, respectively. Upon selecting a condensate, a circular region of interest (ROI) was designated for bleaching resulting in a bleached area of around 1 µm. Bleaching at 488 nm for 10 iterations with 0.5 ms spot bleach duration resulted in simultaneous bleaching of all fluorophores. FRAP acquisitions were obtained within a window from ∼20 min to 1 h after starting the phase separation reactions. One image before, followed by acquisitions every 3 s after bleaching were captured for up to 180 s.

The imaging of condensates in CRISPaint lines with endogenous edits was performed on a Stellaris laser scanning confocal microscope (Leica) with a tunable white light laser (WLL) used to excite Venus at 515 nm and RFP at 554 nm. LAS X software (Leica) was used for image acquisition with a 63X oil objective and z-stacks were taken at 8X zoom in Lightning Mode. At optimized settings, this mode allows deconvolution of the data to reach super-resolution (down to 120 nm). Acquisition was performed with a pinhole size of 1 AU, a refractive index value of 1.518, and an adaptive deconvolution strategy with other settings at default.

### Western Blot and co-immunoprecipitation (Co-IP)

For Western Blots of cultured cells, cell pellets harvested as described above were processed as described^122^. Briefly, pellets were lysed in 2X Laemmli buffer (4% SDS, 20% glycerol, 120 mM Tris-HCl, pH 6.8). Lysates were boiled for 10 min, and the relative protein amount was assessed using a NanoDrop spectrophotometer. 6X Laemmli dye (0.4% (w/v) bromophenol blue, 10% β-ME, 100 mM Tris-HCl, pH 6.8) was supplemented only after the NanoDrop measurement. For Western Blots from wing disc samples, 30-50 discs were dissected in PBS and kept on ice until the collection was completed. The discs were centrifuged (1000*g, 3 min) to remove supernatant, flash-frozen in liquid nitrogen and stored at –70°C. To obtain wing disc lysate, RIPA buffer (10 mM Tris, pH 8, 140 mM NaCl, 1 mM EDTA, 0.5 mM EGTA, 1% (w/v) Triton X-100, 0.1% (w/v) SDS, 0.1% (w/v) Sodium deoxycholate, 0.2 mM PMSF) was added to the discs and the samples were sonicated on ice for 15 min (10 s ON/15 s OFF with amplitude 40%) before addition of SDS loading buffer and boiling for 10 min. Lysates from 30-50 discs were separated in 8% SDS-PAGE gels and transferred to nitrocellulose membranes overnight at 4°C with 30 V constant voltage. Following the transfer, blots were blocked in 5% (w/v) skim milk powder in PBS with 0.3% Tween-20 (PBST). The following primary antibodies were used, diluted in blocking solution supplemented with 0.02% (w/v) sodium azide: rabbit anti-GFP (1:3000, Proteintech) for detection of Venus-tagged proteins, mouse anti-α-Tubulin (1:3000, Sigma-Aldrich), rabbit anti-Pc (1:3000) and rabbit anti-Ph (1:3000). The blots were incubated with primary antibody for 2-3 h at room temperature. Following three washes with PBST, blots were incubated for 45 min at room temperature with the following fluorescent secondary antibodies diluted in blocking solution supplemented with 0.02% (w/v) sodium azide: goat anti-mouse IgG Alexa Fluor 680 conjugated (1:25000, Thermo Fisher Scientific) and goat anti-rabbit IgG IRDye 800 CW conjugated (1:25000, LICOR). Following three washes with PBST, membranes were imaged using an Odyssey CLx imager (LICOR). Western Blots on cell samples were performed in three biological replicates with similar results highlighting expression of full-length proteins. For wing discs, Western Blots were performed on four biological replicates but for technical reasons only two replicates of SAMsurf were of sufficient quality.

For Co-IP experiments, cell pellets were harvested as described above scaling up to 20 million Kc167 cells or confluent 10-cm dishes for the mammalian cell lines. The Co-IP protocol was same for all cell types except that the lysis buffer and the downstream processing for the mammalian samples were different. The lysis buffer for Kc167 cells was buffer F (20 mM Tris, pH 8, 500 mM NaCl, 20% glycerol, 10 mM MgCl_2_, 0.4 mM EDTA, 0.5 mM DTT, 0.1% NP 40) and the lysis buffer for mammalian cells was 20 mM Hepes-KOH, pH 7.5, 150 mM KCl, 10 mM MgCl2, 0.5 mM DTT, 0.5% NP 40. The following concentrations of protease inhibitors were added to the lysis buffers: 0.2 mM PMSF, aprotinin 10 μg/ml, Leupeptin 10 μg/ml, Pepstatin 2 μg/ml, TLCK 50 μg/ml, 1.6 μg/ml Benzamidine and 10 μg/ml phenanthroline. After lysis of cells for 1 h at 4°C on a rotator, the lysate was clarified by centrifugation (20000*g, 10 min at 4°C) and 10% of the supernatant was removed as an input sample (for Kc167 samples only). The remaining supernatant was added to anti-GFP resin (Abnova) (which binds Venus-tagged proteins) pre-equilibrated with lysis buffer and incubated overnight at 4°C on a rotator. The next day, for Kc167 samples, the resin was washed four times with 1 ml lysis buffer for 5 min at 4°C on a rotator, after which 2X SDS loading buffer was added directly to the beads. The samples were boiled for 10 min to elute bound protein. Input and IP samples were loaded on SDS-PAGE gels, after which transfer to nitrocellulose membrane and development of the blot were performed as described above. Bands from CoIP blots were quantified using ImageQuant TL software (v8.1.0.0, Cytiva). To calculate Pc IP relative to the relevant WT protein, the intensity of Pc was first normalized to the intensity of the Venus-tagged protein, and then normalized to WT. Statistical comparisons (ANOVA followed by Holm-Šidák test for multiple comparison; p-values are from multiple comparison test) were calculated, and graphs were plotted using Graphpad Prism 10 software. For samples from mammalian cells, after overnight binding, the resin was washed four times with 1 ml lysis buffer for 5 min at 4°C on a rotator followed by 3 washes with 50 mM ammonium bicarbonate dissolved in mass spectrometry grade H_2_O (Thermo Fisher Scientific). The beads were kept in a small volume of this solution on ice before downstream processing for mass spectrometry (details below).

### Trypsin digestion and LC-MS/MS

The on-bead proteins were first incubated in 4 M urea/100 mM ammonium bicarbonate for 10 min at room temperature, on an Eppendorf MixMate (600 rpm) and then diluted below 2 M urea for a trypsin digestion performed overnight on an Eppendorf Thermomixer at 37°C (450 rpm) using 0.25 µg Sequencing Grade Modified Trypsin (Promega). The samples were then reduced with 9 mM DTT at 37°C for 30 min and, after cooling for 10 min, alkylated with 16.7 mM iodoacetamide at room temperature for 20 min in the dark. The supernatants were acidified with trifluoroacetic acid and salts/detergents removal was performed using MCX cartridges (Waters Oasis MCX 96-well Elution Plate) following the manufacturer’s instructions. After elution in 10% ammonium hydroxide/90% methanol (v/v), samples were dried with a Speed-vac, reconstituted under agitation for 15 min in 75 µl of 1% ACN 1% FA and loaded into a 75 μm i.d. × 250 mm PicoFrit Explore C18 column (New Objective) installed in a Proxeon 1200 LC system (Thermo Scientific). The buffers used for chromatography were 0.2% formic acid (buffer A) and 85% acetonitrile/0.2% formic acid (buffer B). Peptides were eluted with a three-slope gradient at a flowrate of 300 nL/min. Solvent B first increased from 3 to 38% in 100 min, from 38 to 58% in 10 min and then from 58 to 90% B in 2 min. The LC system was coupled to an Orbitrap Fusion mass spectrometer (Thermo Scientific) through a Nanospray Flex Ion Source (Thermo Scientific). Nanospray and S-lens voltages were set to 1.3-1.8 kV and 50 V, respectively. Capillary temperature was set to 250°C. Full scan MS survey spectra (m/z 360-1500) in profile mode were acquired in the Orbitrap with a resolution of 120,000 with a target value at 8e5. The 25 most intense peptide ions were fragmented in the HCD collision cell and analyzed in the linear ion trap with a target value at 1.8e4 and a normalized collision energy at 29%. Target ions selected for fragmentation were dynamically excluded for 30 s after two MS/MS spectra.

### Mass spectrometry analysis and protein identification

Raw files (made available in the Proteomics Identification Database, PRIDE, PXD060324) were uploaded to MaxQuant (v2.4.0.0) and analyzed using default parameters. LC MS/MS data were collected in two runs due to a technical issue with the instrument. We used “match between runs” but adjusted the time window to 10 min. based on comparison of the same sample analyzed in both runs. Unique peptide counts were obtained from the “protein groups” results file.

### Molecular dynamics (MD) simulations and analysis

MD simulations were conducted using a coarse-grained (CG) protein model. We used the Hydropathy Scale (HPS) – Urry model^50^, a recently developed single bead per amino acid CG model which has been shown to capture intrinsically disordered protein phase separation in significant agreement with *in vitro* behavior. In the HPS framework^123^ the total interaction energy of the system comes from three different sources, nonbonded interactions driven by hydropathy, electrostatics, and bonded interactions between consecutive amino acids on the protein sequence,

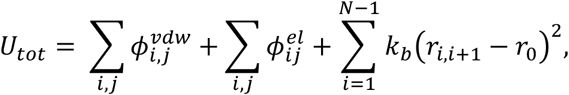

Here the first two terms are because of nonbonded interactions whereas the last term comes from the harmonic springs connecting bonded amino acids. *k*_b_ = 20 kJ/Å is the spring constant for the harmonic potential and r_0_ = 3.82 Å is the equilibrium bond length. The first term 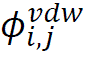 is the short-range van der Waals interaction between residues i and j, and is modeled using the Ashbaugh and Hatch^124^ functional form given by,

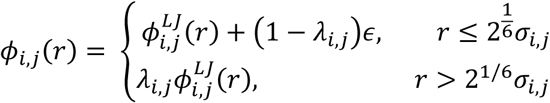

Where 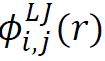 is the standard Lennard-Jones (LJ) potential,

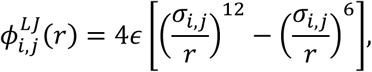

ϵ = 0.2 kcal/mol is the overall energy parameter and the 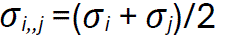 is the distance parameter that is the average van der Waals distance between residues *i* and *j*.

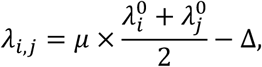

where 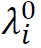 is the hydropathy parameter for amino acid *i* which is derived from the Urry hydropathy scale^125^ after it is normalized to range from 0 to 1. μ = 1.0 is the scale parameter and Δ = 0.08 is the shift parameter both which are derived via an optimization procedure to provide improved agreement with *in vitro* behavior of intrinsically disordered proteins. 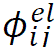 represents the contribution from the electrostatic interactions between fully charged amino acids (Lys, Arg: +1.0 and Asp, Glu: –1.0) we used the Debye-Hückel functional form,

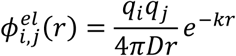

Where *q_i_* is the charge of residue of *i* and is located at the centre of the bead, D=80 (dielectric constant of water) is the dielectric constant of the medium, and *k* is the inverse Debye screening length which is a proxy for the shielding effect of salt ions (1 nm^-1^ is equivalent to roughly 100 mM of salt).

The linker-SAM model was constructed by connecting the disordered linker to the SAM domain (PDB 1KW4) using MODELLER^126^. Both mini-Ph and full-length Ph structures were modeled using AlphaFold2^127^ All single chain simulations were performed for 5 μs using LAMMPS^128^. Multichain simulations in a cubic box and coexistence phase simulations using slab geometry were conducted for 5 μs using HOOMD-Blue 2.9.3^129^, following the protocol described previously^54,130^. In these simulations, the SAM folded domain was treated as a rigid body using the hoomd.md.constrain.rigid function^131^, while other regions remain flexible. The simulation box dimensions and number of protein chains were selected to minimize the potential impact of finite-size effects, as done in our previous work^53^. All the simulations were performed at 300 K.

The simulation trajectories were analyzed after discarding the first 1 μs as equilibration period. Clustering analysis was performed using the freud package^132^ with a 7.5 Å cutoff for nearest neighbor finding. For contact map analysis, two residues were considered in contact if the distance between them was less than 1.5 times arithmetic mean of their van der Waals radii. Simulations snapshots were visualized using VMD^133^. Protein sequences were compiled in **Table S7**.

### Image analysis of condensates

Images of the Cy3 channel (protein) were analyzed in CellProfiler (v4.2.5), after channels were split and images exported using Zen 2 Blue Edition software (Zeiss). Condensates were identified using adaptive minimum cross-entropy thresholding (250 pixel adaptive window size) with a typical object diameter of 3 to 500 and a lower bound for threshold set to: 0.03 for mini-Ph WT and PHC3L with DNA, 0.04 for mini-PH^PHC3L^ without DNA, mini-Ph^SAMsurf^ and mini-Ph^EH^, 0.05 for mini-Ph^SAMsurf-EH^ and 0.08 for mini-Ph^PHC3L-EH^. Adjusting these lower bounds was important to avoid artifacts, and the values were equally or more stringent (i.e. higher) for those samples that formed condensates at lower concentration. Declumping was set to “Shape” with dividing lines based on “Intensity”. Other options were automatic, and holes were filled after thresholding and declumping. The total area covered by condensates as well as individual condensates sizes and shapes were measured and exported. Sizes were converted to µm^2^ using the factor 0.045 µm^2^/px unit. For analysis and comparison of condensate sizes at 4 µM between protein variants, data were filtered based on the form factor 1.1. The form factor measurement in CellProfiler yields values higher than 1 (perfect circles) for small, irregularly shaped structures, which may be artifacts or small fibers. Since these structures would dominate the size distributions (especially for mini-Ph^SAMsurf-EH^) the filtering improved the comparison of similar condensate populations across samples. Unfiltered data were used to calculate the total area covered and for plotting condensate sizes across concentrations. The effect of the filtering can be observed by comparing the 4 µM point in the titration and comparison (e.g. **Figure 4C**). Statistical comparisons for condensate sizes were calculated (Kruskal-Wallis test followed by Dunn’s multiple comparison tests; p-values are from multiple comparison test) and graphs were plotted using GraphPad Prism 10. For condensate size comparison across concentrations only neighbouring concentrations were compared, while for comparisons at 4 µM all pairs were compared.

### FRAP analysis

First, using Fiji (ImageJ, v1.54g) protein (Cy3) and DNA (YOYO-1) channels were split, and background was subtracted using the rolling ball algorithm with a radius of 50. Subsequent analyses were done with publicly available ImageJ plugins (developed by Jay Unruh, Stowers Institute for Medical Research, Kansas City, MO). First, “roi average subtract jru v1” and “roi average divide jru v1” with an ROI outside condensates (background subtraction) and an ROI in a non-bleached condensate (detrending) were used, respectively. Then, “create spectrum jru v1” was used on an ROI in the bleached area to obtain the FRAP trace that were saved as plot objects. Individual FRAP traces from each condition were then combined (“combine all trajectories jru v1”) and normalized from 0 to 1 (“normalize trajectories jru v1” with Min_Max option). The normalized traces were then saved as .csv files. FRAP fits were done on the combined data, or on individual traces, in GraphPad Prism 10 using single or double exponential fits depending which model fits better (Sum-of-squares F test on combined data with default settings). Fit parameters for individual traces were extracted and compared using Kruskal-Wallis test followed by Dunn’s test for multiple comparison (p-values are from multiple comparison test). The single and double exponential equations were as follows:

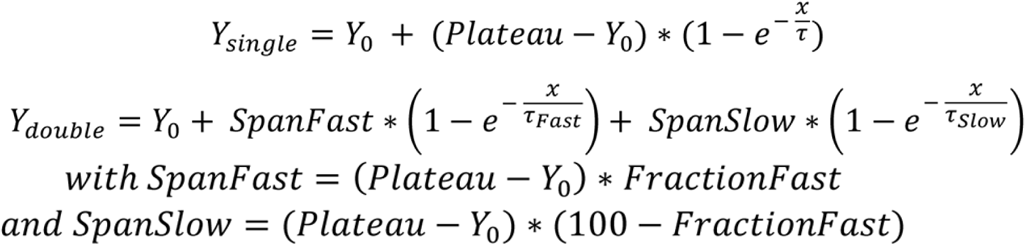

### Image analysis of live cells

All image analysis pipelines were set up with CellProfiler. If not otherwise indicated, a central z-slice was extracted from stacks for analysis. For stable Kc167 lines with inducible mini-Ph expression, the RFP channel was used to segment nuclei with global minimum cross-entropy thresholding for objects with diameter between 25 to 60 pixels and smoothing with a sigma of 2. Declumping was set to “Shape” with dividing lines set to “Shape”, while other options were set to automatic and speed-up using lower resolution image, with holes filled after both thresholding and declumping. Condensates were then identified with global three-class Otsu thresholding (middle intensity class assigned to foreground) for objects with diameter between 5 to 15 pixels and a lower bound for threshold set to: 0.4 for mini-Ph, 0.05 for mini-Ph^PHC3L^ and mini-Ph^SAMsurf^. Adjusting these lower bounds was important to avoid artifacts. For mini-Ph, which typically formed condensates at much higher concentrations than the other constructs, using smaller values for the lower threshold bound resulted in segmenting whole nuclei (featuring high overall Venus signal) instead of condensates. Declumping was set to “Intensity” with dividing lines based on “Intensity” with other options as for nuclei above. For transfected Kc167 cells with mini-Ph variants, the same pipeline was used, but with adaptive three-class Otsu thresholding (middle intensity class assigned to foreground) with an adaptive window size of 60 pixels for objects with diameter between 5 to 15 pixels and a lower bound for threshold that was set to 0.12 for all variants. For transfected Kc167 cells with full-length Ph variants, the RFP channel was first inverted followed by enhancing dark holes with feature sizes between 25 to 60 (which functions like rolling ball background subtraction). Nuclei were then segmented as described above. For condensate segmentation, the rolling ball background subtraction was performed with feature size between 2 to 15, after which condensates were identified like for transfected mini-Ph variants but without lower bound on the threshold (except for SAMsurf variants, which had a lower bound of 0.002). For mammalian cells, the DRAQ5 channel was used to segment nuclei with global two-class Otsu thresholding for objects with diameter between 40 to 200 pixels and smoothing with a sigma of 2. Declumping was set to “Intensity” with dividing lines set to “Intensity”, while other options were set to automatic, with holes filled after both thresholding and declumping. Speckles in the Venus channel were enhanced with a feature size of 8 with speed and accuracy set to slow. Condensates were then identified with adaptive minimum cross-entropy thresholding using an adaptive window size of 100 and a lower bound for threshold set to 0.05 for Ph variants and 0.04 for PHC3 variants. Declumping was set to “Intensity” with dividing lines set to “Intensity”, while other options were set to automatic and speed-up using lower resolution image, with holes filled after both thresholding and declumping. For CRISPaint lines z-stacks were flattened using maximum intensity z-projection performed in FIJI (ImageJ), before the RFP channel was used to segment nuclei with global minimum cross-entropy thresholding for objects with diameter between 75 to 500 pixels. Declumping was set to “Shape” with dividing lines set to “Intensity”, while other options were set to automatic, with holes filled after both thresholding and declumping. Nuclei with a form factor below 0.4 were filtered out to remove artifacts. Then, condensates were identified with a manual global threshold set to 0.2 for objects with a diameter between 4 to 50 (objects touching the border were removed but those outside the diameter range were not discarded). No declumping was performed. For all cell live imaging, condensate sizes per nucleus and nuclear Venus intensities were exported. Sizes were converted to µm^2^ using the factor 0.045 µm^2^/px unit for cells imaged on the spinning disc, and 6.2e-4 µm^2^/px unit for cells imaged with the Stellaris microscope (CRISPaint lines, Figure 7). Statistical comparisons were calculated (Kruskal-Wallis test followed by Dunn’s multiple comparison test; p-values are from multiple comparison test) and graphs were plotted using GraphPad Prism 10. Definitive outliers were removed from mean condensate sizes for mammalian cells and CRISPaint lines in the presented graphs using the ROUT method (Q=0.1) in GraphPad, which did not change the overall results of the statistical analysis.

### Image analysis for nuclear size measurements

Z-stacks of DAPI-stained nuclei were first flattened using maximum intensity z-projection performed in FIJI (ImageJ), before the images were processed in CellProfiler. First, nuclei signal was enhanced using the rolling ball background subtraction described above with a feature size between 1 to 30. Nuclei were identified with global minimum cross-entropy thresholding for objects with diameter between 17 to 50 pixels and smoothing with a sigma of 1. Declumping was set to “Shape” with dividing lines set to “Shape”, while other options were set to automatic and speed-up using lower resolution image, with holes filled after declumping. Nuclei with a form factor between 0.7 to 1 were filtered and their areas were exported. The nuclei were either combined or split into G1 nuclei (those below the median nuclear area) and G2 nuclei (those above the median nuclear area). Statistical comparisons were calculated (Kruskal-Wallis test followed by Dunn’s multiple comparison test; p-values from multiple comparison test) and graphs were plotted using GraphPad Prism 10. Definitive outliers were removed from the combined dataset (before splitting into G1/G2) in the presented graphs using the ROUT method (Q=0.1) in GraphPad, which did not change the overall results of the statistical analysis.

### Image analysis of wing discs

The input .czi stacks of individual wing discs were first split into RFP channel and Venus channel using CellProfiler. Split images were saved as .tif files and a custom ImageJ script was used to generate Sum slices Z-projections. The whole wing disc images (marked by nuclear H2Av-RFP) were transferred to Ilastik (v1.3.3)^113^ for training a pixel classifier. Once the classifier achieved satisfying segmentation of full wing discs, all images were processed to obtain a simple binary segmentation output. The same was repeated for each condition, GFP control, Ph^WT^, Ph^SAMsurf^ and Ph^PHC3L^, respectively. The masks of whole wing discs and corresponding protein expression domains were opened in CellProfiler, where they were resized with a factor of 0.5 (nearest neighbour interpolation). A manual threshold was used to identify wing disc and expression domain as objects, and to fill holes. After resizing the wing disc and expression domain back with a factor of 2, the size of each object was measured. Similarly, the intensities of the Venus and RFP intensities in expression domain were extracted to compare relative Venus expression levels normalized to RFP signal. Statistical comparisons were calculated and graphs were plotted using GraphPad Prism 10. Wing disc sizes were compared (after excluding artifactual values below 0.05 and above 1) using ANOVA followed by Holm-Šidák test for multiple comparison (normality of data was verified with in-built functions; p-values are from multiple comparison test) and Venus expression levels were compared using Kruskal-Wallis test followed by Dunn’s test for multiple comparison (data was not consistently normally distributed; p-values are from multiple comparison test).

## Figure and table preparation

Figures were prepared in Adobe Illustrator 2023 (v27.5, Adobe) and tables were assembled in Excel (Microsoft, v2412) and Adobe InDesign 2025 (v20.0.1, Adobe).

## Quantification and statistical analysis

Details on the exact number of samples analyzed and the number of replicates per experiment for each experiment can be found in the figure legends or in the method description. The meaning of error bars is indicated in the figure legends. Details on statistical tests can be found in the methods description or in the figure legends.

