## Supplementary_information for "How a disordered linker in the Polycomb protein Polyhomeotic tunes phase separation and oligomerization"

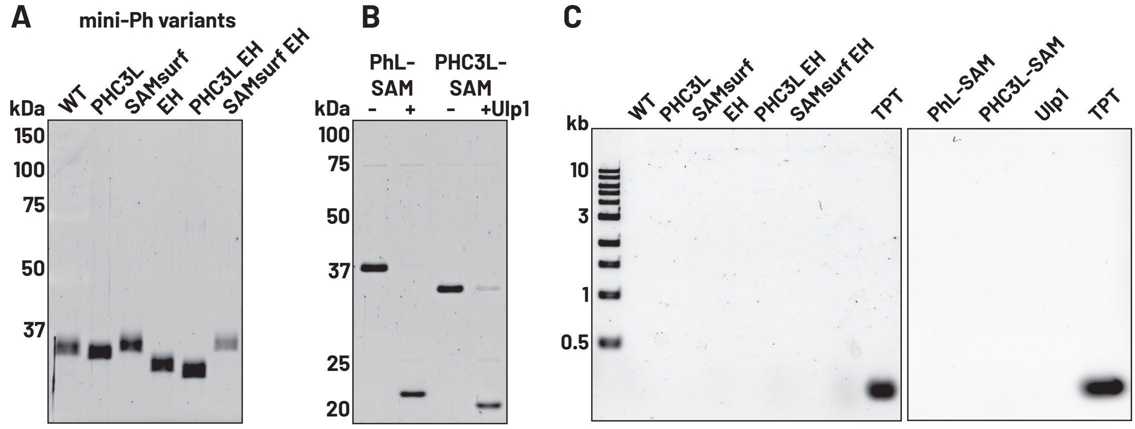


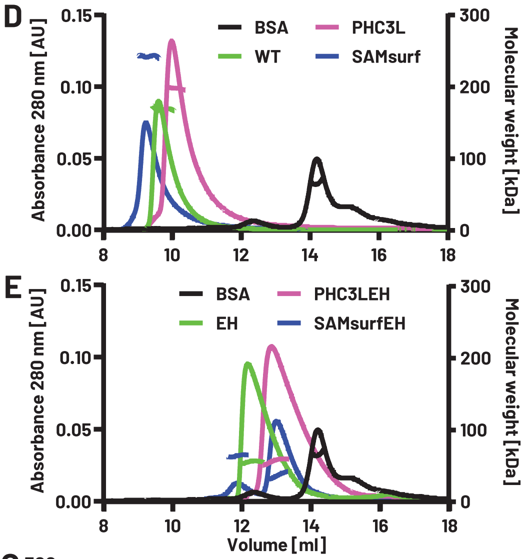


F

| Protein | Molecular weight of monomer [kDa] | Measured molecular weight [kDa] | Measured oligomeric state |
|---|---|---|---|
| mini-Ph WT | 32.8 | 168.1 | 5.1 |
| mini-Ph PHC3L | 33.0 | 197.9 | 6.0 |
| mini-Ph SAMsurf | 32.9 | 242.7 | 7.4 |
| mini-Ph EH | 32.1 | 55.8 | 1.7 |
| mini-Ph PHC3L EH | 32.4 | 57.0 | 1.8 |
| mini-Ph SAMsurf EH | 32.2 | 84.2 and 36.9 | 2.0 and 1.1 |

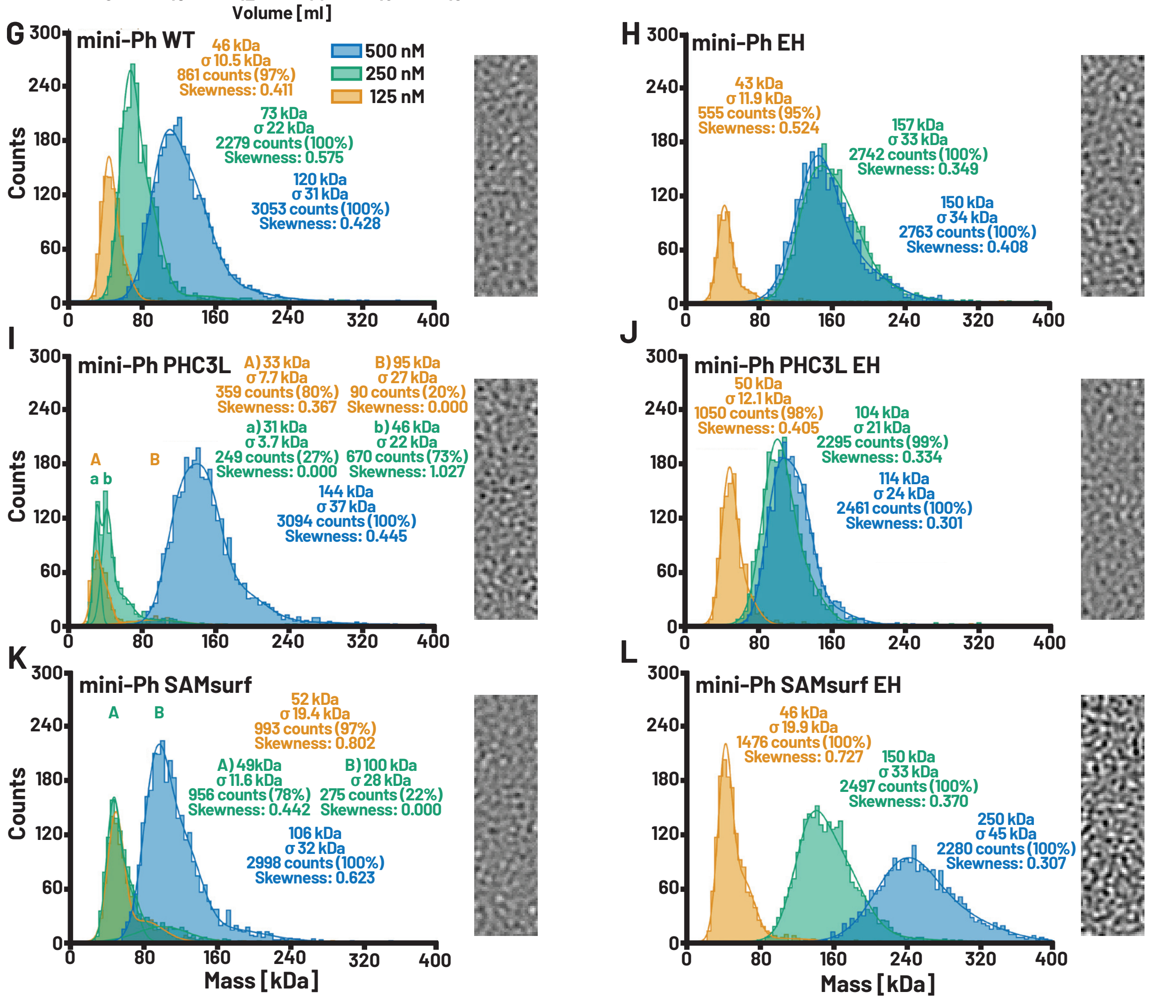

**Figure S1: Quality control of proteins used in this study – related to Figure 1 to 4.** A. SDS-PAGE of mini-Ph variants purified from Sf9 cells (WT, PHC3L, SAMsurf) and from *E. coli* (EH, PHC3L-EH, SAMsurf-EH). Note that the Sf9 and E. coli proteins differ in the presence of an N-terminal Flag tag (Sf9), or a C-terminal 6X-His tag. The altered migration of the variants is consistently observed. B. SDS-PAGE of His-SUMO-PhL-SAM and -PHC3L-SAM purified from E. coli. SUMO protease Ulp1 cleaves the His-SUMO tag from linker-SAM proteins resulting in size shift. C. DNA agarose gel of proteins in A+B after proteinase K treatment indicates absence of DNA contaminations in the purified proteins. 156 bp TPT DNA (50 ng) used for *in vitro* phase separation assays was loaded as a reference. In the left-hand gel, the lane next to TPT is not empty but contains an irrelevant protein that is slightly contaminated with nucleic acid. D, E. SEC-MALS traces and size estimates for mini-Ph variants with WT SAM (D) and EH SAM (E) measured in storage buffer (300 mM KCl, 20 mM HEPES pH 7.9, 20% glycerol). BSA was used to calibrate the size measurement and is shown as a reference. F. Summary of SEC-MALS experiments in D, E. G-L. Mass photometry results for mini-Ph variants diluted to different concentrations immediately before measuring in storage buffer without glycerol (300 mM KCl, 20 mM HEPES pH 7.9). Representative images of the slides to the right of each graph indicate the absence of large aggregates.

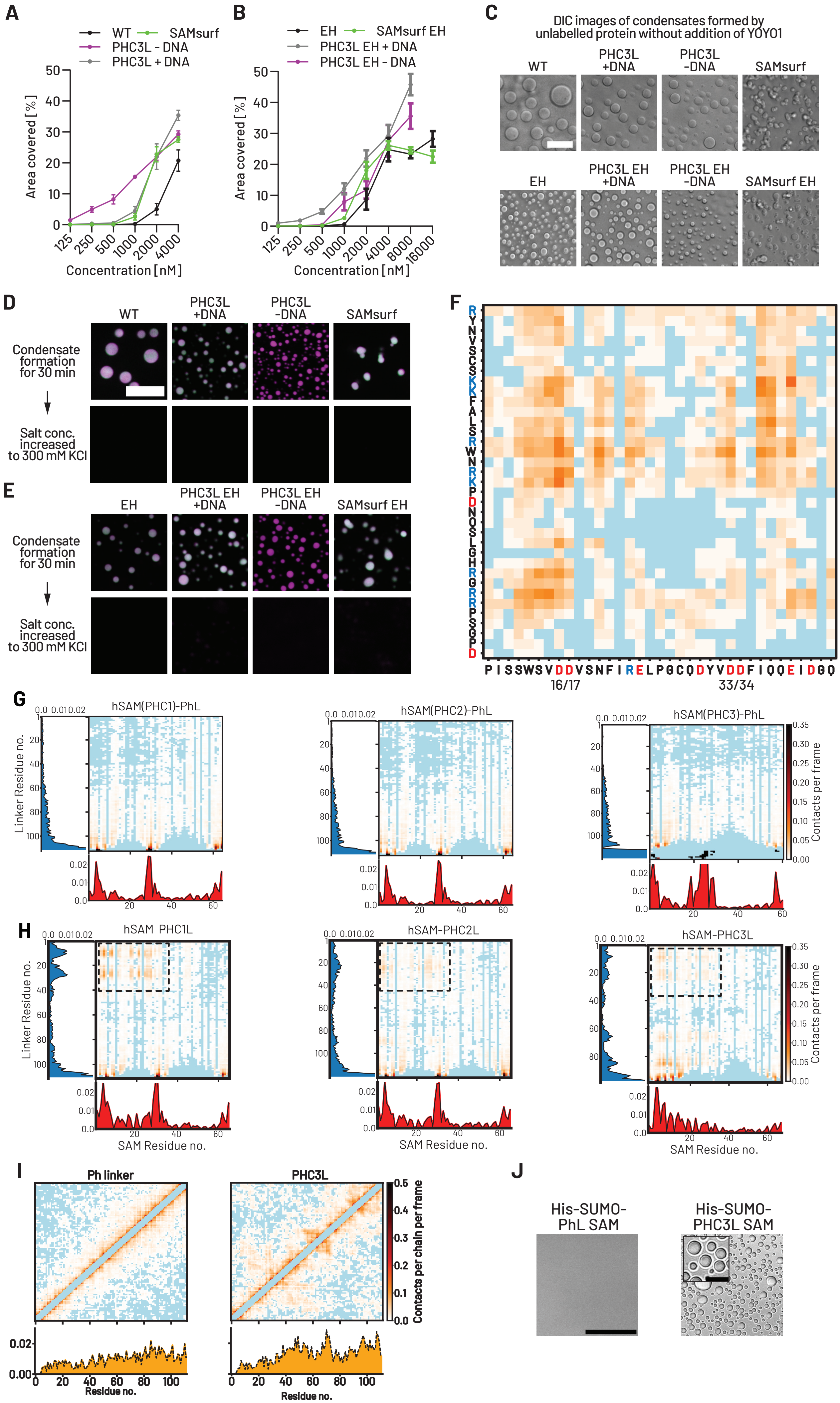

A
WT
SAMsurf
PHC3L - DNA
PHC3L + DNA
Area covered [%]
Concentration [nM]
B
EH
SAMsurf EH
PHC3L EH + DNA
PHC3L EH - DNA
Area covered [%]
Concentration [nM]
C
DIC images of condensates formed by unlabelled protein without addition of YOYO1
WT
PHC3L +DNA
PHC3L -DNA
SAMsurf
EH
PHC3L EH +DNA
PHC3L EH -DNA
SAMsurf EH
D
WT
PHC3L +DNA
PHC3L -DNA
SAMsurf
Condensate formation for 30 min
Salt conc. increased to 300 mM KCl
E
EH
PHC3L EH +DNA
PHC3L EH -DNA
SAMsurf EH
Condensate formation for 30 min
Salt conc. increased to 300 mM KCl
F
16/17
33/34
G
hSAM(PHC1)-PhL
hSAM(PHC2)-PhL
hSAM(PHC3)-PhL
Linker Residue no.
Contacts per frame
H
hSAM PHC1L
hSAM-PHC2L
hSAM-PHC3L
Linker Residue no.
Contacts per frame
SAM Residue no.
I
Ph linker
PHC3L
Contacts per chain per frame
Residue no.
J
His-SUMO-PhL SAM
His-SUMO-PHC3L SAM

**Figure S2: Condensate analysis and controls and MD of linker-SAM proteins – related to Figure 1 to 4.** A, B. Quantification of total area covered by condensates in *in vitro* phase separation assays. These informed titration summaries (qualitative phase diagrams) shown in Fig. 3F (A, includes WT and PHC3L data from Figure 1D as reference) and 4A (B). C. Representative phase contrast images of mini-Ph variants forming condensates in the absence of Cy3-labeled protein or YOYO-1 dye. Scale bar: 20 µm. D, E. Condensate formation of mini-Ph variants with both WT SAM (D) or EH SAM (E) is reversed by increasing the salt concentration to storage conditions (300 mM KCl). Scale bar: 20 µm. F. Zoom on contact map region with increased PHC3L-SAM contact frequency in Figure 2C from single chain simulations. Individual residues are indicated (PHC3L residues on y axis, SAM residues on x axis) with basic residues (K and R) in blue and acidic residues (D and E) in red. Note that simulations were conducted on PHC3L-SAM chains with additional N-terminal residues (RYNVSC) as compared with PhL-SAM. G, H. Contact maps from single chain simulations of PHC SAMs with PhL (G) or corresponding PHCL (H) show contact frequencies between linker residues and SAM residues (non-interacting residues colored in light blue). Increased long-range contacts between linker and SAM (highlighted by dashed rectangle) were observed with PHCLs (compare with Figure 2D). I. Intramolecular interactions from single chain simulations of PhL and PHC3L show increased interactions in the latter. A diagonal band of contacts between 3 neighboring residues is removed from the plot to highlight long range contacts. J. Phase contrast images of HisSUMO-PhL-SAM and HisSUMO-PHC3L-SAM without cleavage of SUMO tag with Ulp1 (compare Figure 2I). Scale bar: 50 µm and 10 µm for inset.

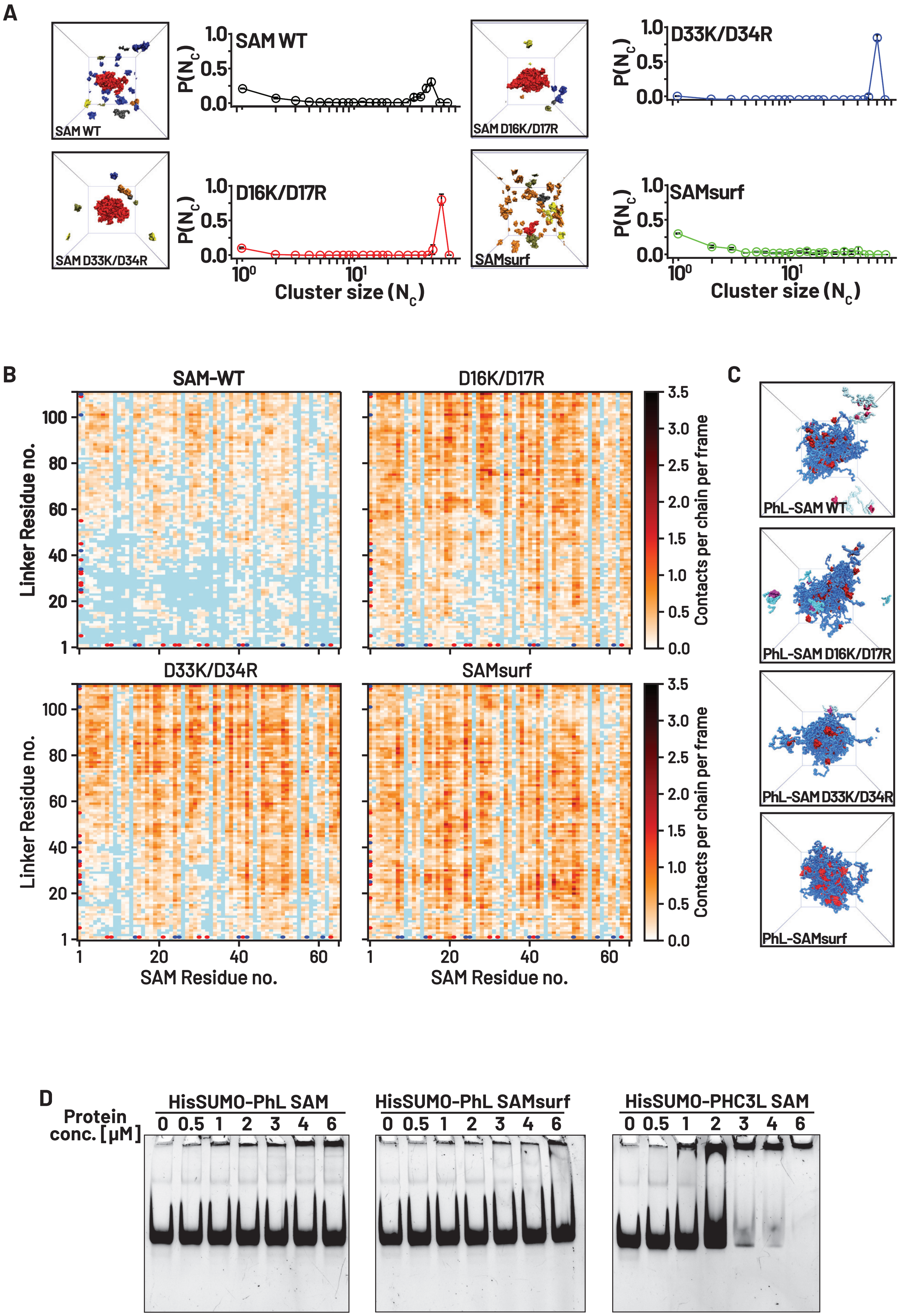
A
SAM WT
SAM D16K/D17R
SAM D33K/D34R
SAMsurf
D33K/D34R
D16K/D17R
P(N_c)
Cluster size (N_c)
B
SAM-WT
D16K/D17R
D33K/D34R
SAMsurf
Linker Residue no.
SAM Residue no.
Contacts per chain per frame
C
PhL-SAM WT
PhL-SAM D16K/D17R
PhL-SAM D33K/D34R
PhL-SAMsurf
D
Protein conc. [µM]
HisSUMO-PhL SAM
HisSUMO-PhL SAMsurf
HisSUMO-PHC3L SAM
0 0.5 1 2 3 4 6

**Figure S3: MD analysis of SAM surface mutations – related to Figure 3.** A. Quantification of SAM clustering and snapshots from multi chain simulations, where chains belonging to the largest cluster are colored red. SAM WT forms some clusters in simulations, which are increased for D16K D17R and D33K D34R but abolished when both pairs of surface mutations are introduced (D16K D17R D33K D34R, "SAMsurf"). B. Intermolecular contact maps from multi chain simulations of PhL with WT SAM, D16K/D17R SAM, D33K/D34R and SAMsurf show overall contact frequencies increase for PhL with SAM surface charge mutants compared to WT SAM (non-interacting residues colored in blue). Basic residues (K and R) are highlighted by blue dots and acidic residues (D and E) by red dots. C. Snapshots from multi chain simulations of PhL with WT SAM (same as Figure 2F) or SAM surface mutants). D. Electrophoretic mobility shift assay of His-SUMO linker-SAM constructs (without Ulp1 cleavage) with 156 bp DNA demonstrates DNA binding by the PHC3L but not by PhL, SAM or SAMsurf.

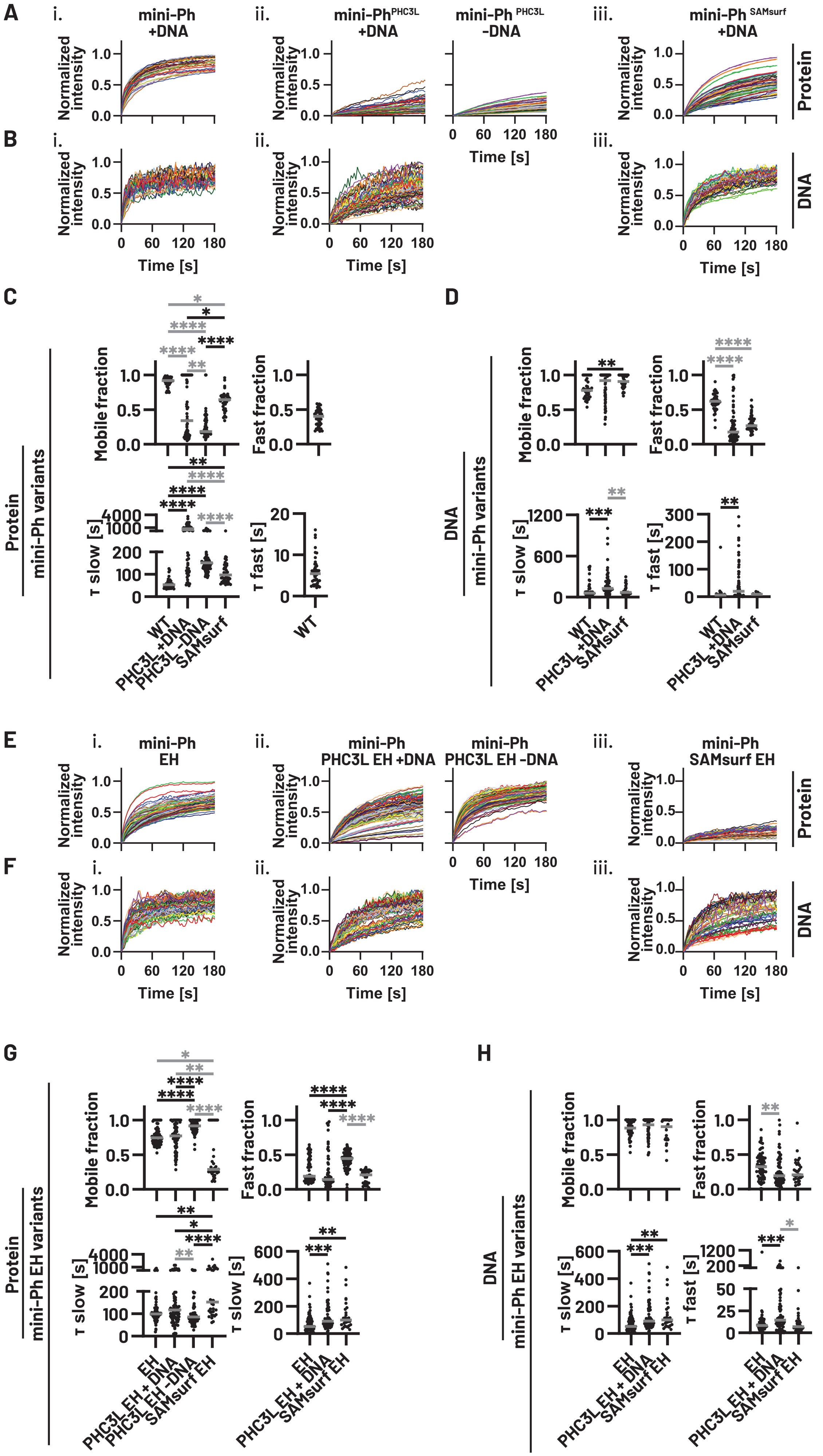
A
B
C
D
E
F
G
H
mini-Ph +DNA
mini-Ph PHC3L +DNA
mini-Ph PHC3L -DNA
mini-Ph SAMsurf +DNA
Protein
DNA
Normalized intensity
Time [s]
mini-Ph variants
Mobile fraction
Fast fraction
τ slow [s]
τ fast [s]
WT
PHC3L +DNA
PHC3L -DNA
SAMsurf
mini-Ph EH
mini-Ph PHC3L EH +DNA
mini-Ph PHC3L EH -DNA
mini-Ph SAMsurf EH
mini-Ph EH variants
EH
PHC3L EH + DNA
PHC3L EH -DNA
SAMsurf EH

**Figure S4: Analysis of individual FRAP traces – related to Figure 4** A, B. Individual protein (A) and DNA (B) FRAP traces of mini-Ph variants with WT SAM underlying the summary curves in **Figure 4D, E**. C, D. Analysis of fit parameters from individual protein FRAP traces in A (C) and DNA FRAP traces in B (D); gray bar indicates median. E, F. Individual protein (E) and DNA (F) FRAP traces of mini-Ph variants with EH SAM underlying the summary curves in **Figure 4G, H**. G,H. Analysis of fit parameters from individual protein FRAP traces in E (G) and DNA FRAP traces in F (H); gray bar indicates median. All parameters are listed in **Table S2**.

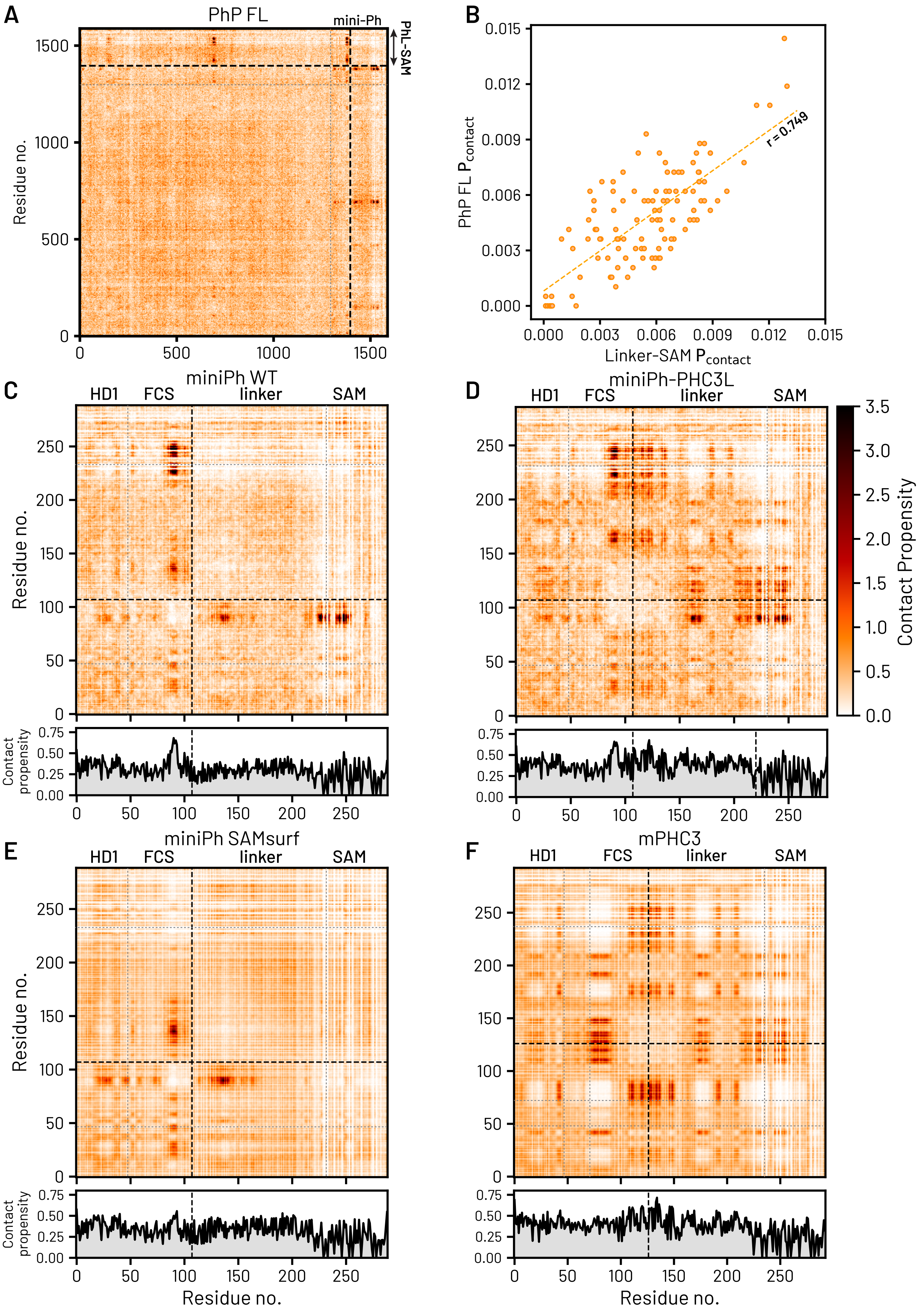

A
PhP FL
mini-Ph
PhL-SAM
Residue no.
B
PhP FL P_contact
Linker-SAM P_contact
r = 0.749
C
miniPh WT
HD1
FCS
linker
SAM
Residue no.
Contact propensity
D
miniPh-PHC3L
Contact Propensity
E
miniPh SAMsurf
F
mPHC3

**Figure S5: MD simulations of full length (FL) Ph and mini-Ph – related to Figure 5.** A. Intermolecular contact maps of FL Ph within the dense phase in the coexistence phase simulation. Region of Ph linker-SAM is highlighted in the map. B. The correlations between the contact probability per residue for linker-SAM alone (x-axis) and corresponding contacts in FL Ph (y-axis) in the CG coexistence phase simulations. C-F. Two-dimensional (as in **Figure 5**) and one-dimensional (bottom) intermolecular contact maps of mini-Ph (C), mini-$Ph^{PHC3L}$ (D), mini-$Ph^{SAMsurf}$ (E), and mini-PHC3 (F). One-dimensional contact map represents the average contact probability per residue along the protein sequence.

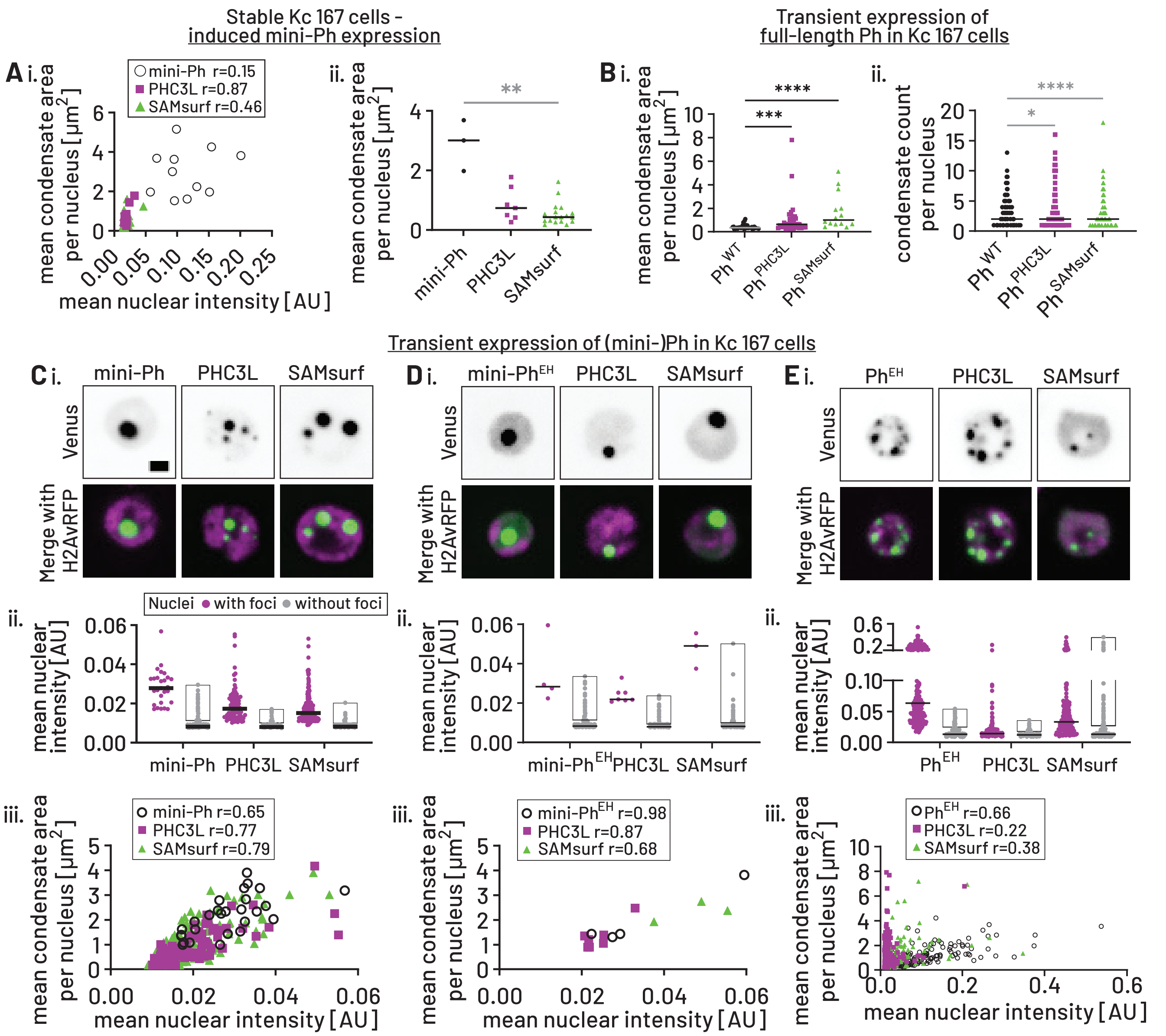


iv.

| | mini-Ph | | PHC3L | | SAMsurf | |
|---|---|---|---|---|---|---|
| | Condensate count per nucleus | | | | | |
| Mean | 1.0 | | 1.3 | | 1.3 | |
| CI of mean | 1.0 | 1.0 | 1.2 | 1.4 | 1.2 | 1.4 |
| Median | 1.0 | | 1.0 | | 1.0 | |
| CI of median | 1.0 | 1.0 | 1.0 | 1.0 | 1.0 | 1.0 |
| | Mean condensate area per nucleus [μm²] | | | | | |
| Mean | 1.3 | | 0.9 | | 1.0 | |
| CI of mean | 1.0 | 1.7 | 0.6 | 1.2 | 0.9 | 1.2 |
| Median | 1.2 | | 0.8 | | 0.9 | |
| CI of median | 1.0 | 1.9 | 0.5 | 1.3 | 0.7 | 1.3 |

iv.

| | mini-Ph$^{EH}$ | | PHC3L | | SAMsurf | |
|---|---|---|---|---|---|---|
| | Condensate count per nucleus | | | | | |
| Mean | 1.0 | | 1.3 | | 1.0 | |
| CI of mean | 1.0 | 1.0 | 0.6 | 2.0 | 1.0 | 1.0 |
| Median | 1.0 | | 1.0 | | 1.0 | |
| CI of median | 1.0 | 1.0 | 1.0 | 3.0 | 1.0 | 1.0 |
| | Mean condensate area per nucleus [μm²] | | | | | |
| Mean | 2.0 | | 1.3 | | 2.4 | |
| CI of mean | 0.1 | 3.9 | 0.8 | 1.8 | 1.4 | 3.4 |
| Median | 1.4 | | 1.2 | | 2.4 | |
| CI of median | 1.3 | 3.8 | 0.9 | 2.5 | 1.9 | 2.7 |

iv.

| | Ph$^{EH}$ | | PHC3L | | SAMsurf | |
|---|---|---|---|---|---|---|
| | Condensate count per nucleus | | | | | |
| Mean | 3.8 | | 3.9 | | 3.6 | |
| CI of mean | 3.4 | 4.2 | 3.5 | 4.4 | 2.9 | 4.2 |
| Median | 3.0 | | 2.0 | | 2.0 | |
| CI of median | 2.0 | 4.0 | 2.0 | 3.0 | 1.0 | 2.0 |
| | Mean condensate area per nucleus [μm²] | | | | | |
| Mean | 0.6 | | 1.0 | | 0.8 | |
| CI of mean | 0.5 | 0.8 | 0.7 | 1.4 | 0.5 | 1.1 |
| Median | 0.5 | | 0.9 | | 0.5 | |
| CI of median | 0.5 | 0.8 | 0.5 | 1.1 | 0.4 | 0.9 |

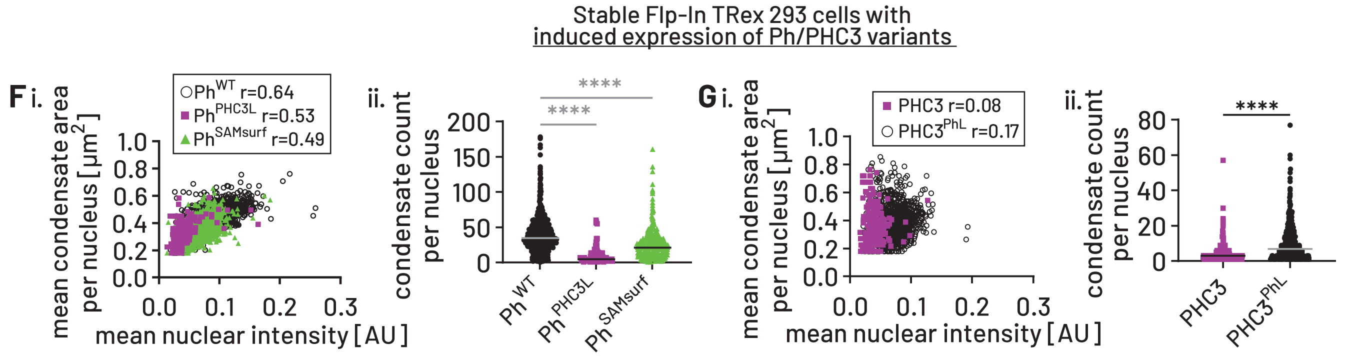

**Figure S6: Additional analyses of cell imaging – related to Figure 6.** A. Additional analyses of condensates in stable Kc167 cells with inducible mini-Ph expression from Fig. 6. i. Quantification of condensate size dependence on relative mini-Ph concentration. Each point represents the mean condensate size for a given nucleus (y-axis) vs. the mean nuclear Venus intensity of the nucleus (x-axis). Pearson correlation coefficients (r) were calculated for each dataset. ii. Mean condensate sizes per nucleus from cells with similar mini-Ph expression levels. Cells were selected in a window of nuclear mean intensities spanning the $5^{th}$ percentile in the mini-$Ph^{PHC3L}$ dataset to the $25^{th}$ percentile in the mini-Ph dataset. Comparisons are to mini-Ph. **: $p<0.005$. B. Additional analyses of condensates in transiently transfected Kc167 cells with full-length Ph expression from Fig. 6. i. Mean condensate sizes per nucleus from cells with similar Ph expression levels. Cells were selected based in a window of nuclear mean intensities spanning the $5^{th}$ to $25^{th}$ percentile in the Ph dataset. Comparisons are to Ph. ***: $p<0.0005$, ****: $p<0.0001$ ii. Quantification of condensates per nucleus. Comparisons are to Ph. *: $p<0.05$, ****: $p<0.0001$. C-E. Analyses of condensates from Kc167 cells transiently transfected with mini-Ph variants (C), mini-$Ph^{EH}$ variants (D) and $Ph^{EH}$ variants (E). i. Representative images of overexpressed (mini-)Ph variants (individual z-slice, H2Av-RFP is a nuclear marker). ii. To compare thresholds for condensate formation, cells were stratified into with (magenta) and without (gray) condensates and the mean nuclear Venus intensity plotted. Bar indicates median and gray box the 10% highest expressing cells without condensates. Data are compiled from three experiments. For C: mini-Ph n=2162 cells (27 with condensates); PHC3L n=1155 (84 with condensates); SAMsurf n=1651 cells (217 with condensates). For D: mini-$Ph^{EH}$ n=2316 cells (4 with condensates); PHC3L n=2417 (7 with condensates); SAMsurf n=2170 cells (3 with condensates). For E: $Ph^{EH}$ n=955 cells (189 with condensates); PHC3L n=1007 (307 with condensates); SAMsurf n=993 cells (139 with condensates). iii. Quantification of condensate size dependence on relative protein concentration as in A ii. Iv. Overview of condensate count per nucleus and mean condensate area per nucleus from cells with similar protein expression levels. For C, the expression window spans the $5^{th}$ to $25^{th}$ percentile in the mini-Ph dataset. For D, all cells were included. For E, the window spanned the $5^{th}$ to $25^{th}$ percentile in the $Ph^{EH}$ dataset. Mean and median are shown along with the respective 95% confidence intervals and were color-coded in comparison to mini-Ph, mini-$Ph^{EH}$ or $Ph^{EH}$ (appearing white), respectively, with smaller values in shades of blue and higher values in shades of orange. F, G. Additional analyses of condensates in mammalian cell lines with inducible Ph expression (F) or PHC3 expression (G) from Fig. 6. Fi, Gi. Quantification of condensate size dependence on relative Ph or PHC3 concentration as in A ii. Fii, Gii. Quantification of condensates per nucleus. Comparisons are to Ph (Fii) or PHC3 (Gii). ****: $p<0.0001$.

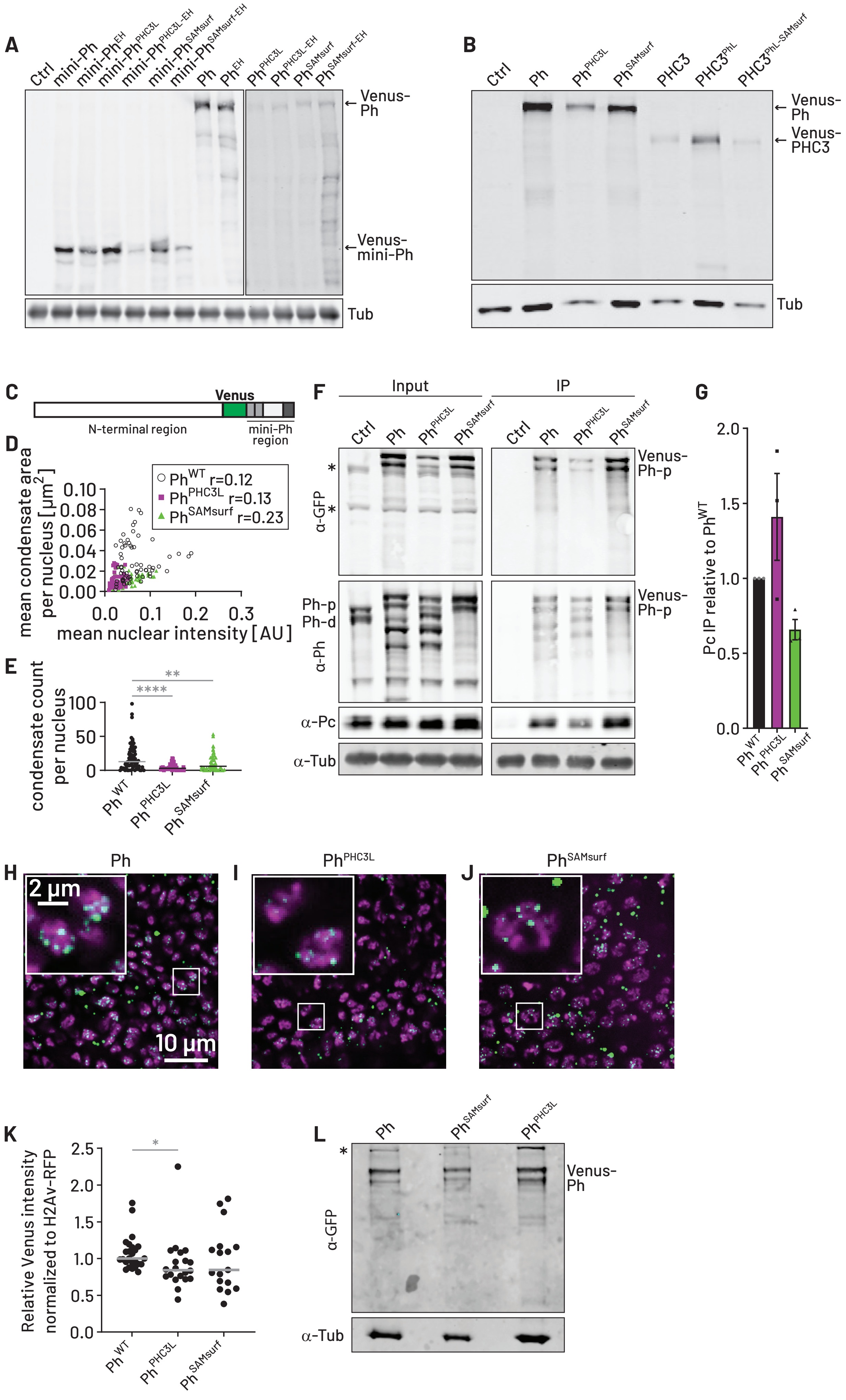

A
Ctrl
mini-Ph
mini-Ph^EH
mini-Ph^PHC3L
mini-Ph^PHC3L-EH
mini-Ph^SAMsurf
mini-Ph^SAMsurf-EH
Ph
Ph^EH
Ph^PHC3L
Ph^PHC3L-EH
Ph^SAMsurf
Ph^SAMsurf-EH
Venus-Ph
Venus-mini-Ph
Tub
B
Ctrl
Ph
Ph^PHC3L
Ph^SAMsurf
PHC3
PHC3^PhL
PHC3^PhL-SAMsurf
Venus-Ph
Venus-PHC3
Tub
C
Venus
N-terminal region
mini-Ph region
D
mean condensate area per nucleus [µm²]
mean nuclear intensity [AU]
Ph^WT r=0.12
Ph^PHC3L r=0.13
Ph^SAMsurf r=0.23
E
condensate count per nucleus
**
****
Ph^WT
Ph^PHC3L
Ph^SAMsurf
F
Input
IP
Ctrl
Ph
Ph^PHC3L
Ph^SAMsurf
α-GFP
Venus-Ph-p
Ph-p
Ph-d
α-Ph
α-Pc
α-Tub
G
Pc IP relative to Ph^WT
Ph^WT
Ph^PHC3L
Ph^SAMsurf
H
Ph
2 µm
10 µm
I
Ph^PHC3L
J
Ph^SAMsurf
K
Relative Venus intensity normalized to H2Av-RFP
Ph^WT
Ph^PHC3L
Ph^SAMsurf
L
Ph
Ph^SAMsurf
Ph^PHC3L
α-GFP
Venus-Ph
α-Tub

**Fig. S7: Additional analysis of cells and wing discs – related to Figure 6 and 7.** A. Representative Western Blot of Venus (mini)-Ph variants in transiently transfected Kc167 cells. Tub is a loading control. (control: Kc167 cells transfected with H2Av-RFP but no Venus transgene). The blot corresponds to cells analyzed in Figure 6F-H and Figure S6E-G. All samples are from the same blot, but the last 4 lanes are differently adjusted for visualization. B. Representative Western Blot of stable Flp-In TRex 293 cell lines (control: Flp-In cells without Venus transgene). Tub is a loading control. The blot corresponds to cells analyzed in Figure 6I-N. C. Schematic of edited Ph indicating the position of the internal Venus tag. D. Quantification of condensate size dependence on relative Ph concentration in cell lines with endogenous edits. Each point represents the mean condensate size for a given nucleus (y-axis) vs. the mean nuclear Venus intensity of the nucleus (x-axis). Pearson correlation coefficients (r) were calculated for each dataset. E. Quantification of condensates per nucleus. Comparisons are against Ph. **: $p<0.005$, ****: $p<0.0001$ F. Representative Western Blot for anti-Venus co-IP from Kc167 cell lines with endogenous edits of *ph.* Pc is a PRC1 subunit and Tub is a loading control and remains associated to the anti-Venus resin despite stringent washing. * indicate non-specific band recognized by the GFP antibody. (control: unedited Kc-167 cells). Note that we do not know why there are two Venus-Ph-p bands, but we did not detect Venus-Ph-d in genotyping analysis. G. Quantification of co-immunoprecipitation experiments (n=3). Bars show mean +/- SEM. H-J. Representative images of Ph condensates in wing imaginal discs expressing Venus-Ph (H), -$Ph^{PHC3L}$ (I) and -$Ph^{SAMsurf}$ (J) (individual z-slice). The Venus channel (in green) was merged with the nuclear marker H2Av-RFP (in magenta). K. Quantification of Venus-Ph intensities from posterior compartment of wing imaginal discs. The signal was normalized to H2Av-RFP and datasets were compared against Ph (*: $p<0.05$). L. Representative Western Blot from wing imaginal discs expressing Ph variants.

```
Ph SAM    …PISSWSVDDVSNFIRELPGCQDYVDDFIQQEIDGQALLLLKEKHLVNAMGMKLGPALKIVAKVESIKEVPPPGEAKDPGAQ
PHC1 SAM  …NPSRWSVEEVYEFIASLQGCQEIAEEFRSQEIDGQALLLLKEEHLMSAMNIKLGPALKICAKINVLKET------------
PHC2 SAM  …EPTKWNVEDVYEFIRSLPGCQEIAEEFRAQEIDGQALLLLKEDHLMSAMNIKLGPALKIYARISMLKDS------------
PHC3 SAM  …EPSIWTVDDVWAFIHSLPGCQDIADEFRAQEIDGQALLLLKEDHLMSAMNIKLGPALKICARINSLKES------------
SAMsurf   …PISSWSVKRVSNFIRELPGCQDYVKRFIQQEIDGQALLLLKEKHLVNAMGMKLGPALKIVAKVESIKEVPPPGEAKDPGAQ
                  ↑                ↑
               "16/17"          "33/34"
```

**Table S1: Multiple sequence alignment of SAM sequences – related to Figure 3.** Sequences of Ph and PHC SAMs were aligned with Ph SAMsurf mutant highlighting acidic-to-basic mutations.

**Protein**

| | Mobile fraction | | | Fast fraction | | | $\tau_{slow}$ [s] | | | $\tau_{fast}$ [s] | | |
|---|---|---|---|---|---|---|---|---|---|---|---|---|
| | Median | CI of median | | Median | CI of median | | Median | CI of median | | Median | CI of median | |
| **mini-Ph WT** | 0.92 | 0.89 | 0.94 | 0.40 | 0.33 | 0.45 | 53.98 | 44.85 | 59.86 | 5.46 | 3.85 | 6.34 |
| **mini-Ph PHC3L + DNA** | 0.34 | 0.19 | 0.53 | | | | 224.10 | 166.90 | 299.40 | | | |
| **mini-Ph PHC3L - DNA** | 0.18 | 0.18 | 0.22 | | | | 151.90 | 142.10 | 159.60 | | | |
| **mini-Ph SAMsurf** | 0.64 | 0.60 | 0.69 | | | | 97.34 | 80.52 | 117.60 | | | |
| **mini-Ph EH** | 0.74 | 0.71 | 0.79 | 0.19 | 0.16 | 0.32 | 99.02 | 91.34 | 105.60 | 11.54 | 9.56 | 13.71 |
| **mini-Ph PHC3L EH + DNA** | 0.77 | 0.74 | 0.83 | 0.14 | 0.13 | 0.16 | 116.80 | 98.21 | 128.20 | 16.22 | 12.00 | 18.57 |
| **mini-Ph PHC3L EH - DNA** | 0.92 | 0.90 | 0.94 | 0.45 | 0.40 | 0.47 | 85.84 | 77.79 | 93.29 | 10.36 | 9.85 | 11.39 |
| **mini-Ph SAMsurf EH** | 0.29 | 0.25 | 0.40 | 0.21 | 0.11 | 0.26 | 152.10 | 110.20 | 518.30 | 7.83 | 4.11 | 10.42 |

**DNA**

| | Mobile fraction | | | Fast fraction | | | $\tau_{slow}$ [s] | | | $\tau_{fast}$ [s] | | |
|---|---|---|---|---|---|---|---|---|---|---|---|---|
| | Median | CI of median | | Median | CI of median | | Median | CI of median | | Median | CI of median | |
| **mini-Ph WT** | 0.78 | 0.71 | 0.81 | 0.62 | 0.58 | 0.65 | 60.96 | 53.16 | 78.12 | 6.42 | 4.92 | 8.44 |
| **mini-Ph PHC3L + DNA** | 0.92 | 0.77 | 1.00 | 0.18 | 0.12 | 0.23 | 126.10 | 104.60 | 161.70 | 19.79 | 7.61 | 35.24 |
| **mini-Ph PHC3L - DNA** | | | | | | | | | | | | |
| **mini-Ph SAMsurf** | 0.90 | 0.86 | 0.98 | 0.27 | 0.24 | 0.34 | 70.70 | 55.69 | 87.26 | 8.60 | 7.79 | 10.68 |
| **mini-Ph EH** | 0.88 | 0.84 | 0.92 | 0.33 | 0.25 | 0.39 | 52.16 | 46.04 | 85.50 | 8.26 | 6.99 | 9.63 |
| **mini-Ph PHC3L EH + DNA** | 0.93 | 0.87 | 0.99 | 0.19 | 0.15 | 0.24 | 90.29 | 76.07 | 100.60 | 14.34 | 11.47 | 17.24 |
| **mini-Ph PHC3L EH - DNA** | | | | | | | | | | | | |
| **mini-Ph SAMsurf EH** | 0.90 | 0.74 | 1.00 | 0.21 | 0.17 | 0.28 | 99.90 | 66.19 | 135.90 | 7.12 | 5.76 | 12.52 |

**Table S2: Summary of FRAP data – related to Figure 4.** Overview of fit parameters derived from individual FRAP traces in Figure S4. Median values for protein (top) and DNA (bottom) are shown along with the lower and upper limits of the 95% confidence interval.

| cell line | alleles detected | junction sequence | description | color code |
| --- | --- | --- | --- | --- |
| Ph$^{WT}$ | ph-p | TSCTSTTTTTTTSG | truncated at editing site (S is last aligned aa) | ph-p or -d upstream |
| | ph-d | SSVPFSVSPSQTAICRQRRQR* | truncated near editing site | Venus |
| | ph-p-Venus | TSCTSTTTTTTTSGGMVPILVELDGD | in frame edit, missing aa2-11 of Venus | downstream ph-p |
| Ph$^{PHC3L}$ | ph-p | TTTTYMVACEQCGKM | in frame deletion of aa 1287-1361 (HD1 and first 3 aa of FCS) | |
| | ph-d | TTTTTTSVMDRRISPRR* | truncated after editing site | |
| | ph-p-Venus | TSTTTTTTSGGMVSKGEEL | in frame edit | |
| Ph$^{SAMsurf}$ | SAMsurf ph-p | only Venus fusion detected | | |
| | SAMsurf ph-d | not detected | | |
| | SAMsurf Venus | TSCTSTTTTTTTSGGMVSKGEELFTGVVP | | |

**Table S3: Genotyping results from edited Kc cells – related to Methods.**

| | Kc | | $Ph^{WT}$ | | $Ph^{PHC3L}$ | | $Ph^{SAMsurf}$ | |
|---|---|---|---|---|---|---|---|---|
| | **Mean** | **SEM** | **Mean** | **SEM** | **Mean** | **SEM** | **Mean** | **SEM** |
| ph-p | 0.23 | 0.02 | 0.97 | 0.1 | 0.48 | 0.02 | 0.55 | 0.04 |
| ph-d | 0.43 | 0.05 | 0.8 | 0.11 | 0.65 | 0.15 | 0 | 0 |
| psc | 0.57 | 0.14 | 0.87 | 0.11 | 2.28 | 0.55 | 0.71 | 0.13 |
| suz2 | 0.49 | 0.12 | 0.9 | 0.13 | 1.26 | 0.21 | 0.99 | 0.02 |
| chinmo | 0.71 | 0.14 | 0.72 | 0.14 | 0.44 | 0.08 | 0.63 | 0.14 |
| cycb | 1.28 | 0.17 | 0.89 | 0.07 | 1 | 0.11 | 1.03 | 0.02 |
| tsh | 2.01 | 0.27 | 1.42 | 0.33 | 4.21 | 0.28 | 3.11 | 0.5 |
| zfh1 | 0.38 | 0.06 | 0.72 | 0.16 | 0.47 | 0.14 | 0.53 | 0.09 |
| zfh2 | 0.03 | 0 | 0.88 | 0.13 | 0.59 | 0.09 | 0.82 | 0.11 |
| bbg | 5.41 | 1.51 | 0.7 | 0.15 | 0.49 | 0.1 | 0.5 | 0.11 |
| ptc | 0.1 | 0.02 | 0.79 | 0.11 | 0.59 | 0.12 | 1.23 | 0.32 |
| pask | 0.49 | 0.18 | 1.27 | 0.24 | 1.05 | 0.38 | 1.28 | 0.42 |
| thor | 2.83 | 0.87 | 1.2 | 0.18 | 0.81 | 0.09 | 0.94 | 0.11 |
| sans | 0.23 | 0.08 | 0.91 | 0.06 | 0.72 | 0.06 | 1.54 | 0.05 |
| cg44251 | 3.61 | 1.16 | 1.07 | 0.17 | 1.7 | 0.55 | 0.54 | 0.11 |
| acxd | 1.53 | 0.8 | 0.58 | 0.27 | 0.37 | 0.15 | 0.36 | 0.15 |
| pnr | 0.01 | 0 | 0.78 | 0.11 | 0.83 | 0.19 | 0.87 | 0.13 |
| socs36e | 0.79 | 0.12 | 0.8 | 0.12 | 0.91 | 0.22 | 0.87 | 0.16 |
| stat92e | 1.74 | 0.27 | 0.84 | 0.09 | 0.68 | 0.05 | 0.63 | 0.06 |
| upd2 | 1.34 | 0.36 | 1.15 | 0.21 | 0.72 | 0.07 | 0.44 | 0.03 |
| upd3 | 1.57 | 0.45 | 0.92 | 0.12 | 0.71 | 0.11 | 0.97 | 0.29 |
| mir_8 | 0.74 | 0.42 | 0.58 | 0.24 | 0.62 | 0.28 | 0.08 | 0.01 |
| mir_bantam | 0.27 | 0.04 | 0.54 | 0.26 | 0.45 | 0.21 | 0.66 | 0.37 |
| mir_184 | 0.66 | 0.34 | 1.25 | 0.71 | 0.66 | 0.25 | 0.9 | 0.34 |

**Table S4: Summary of RT-qPCR data – related to Figure 7.** Overview of mRNA expression as fold change over $Ph^{WT}$ from RT-qPCR in Figure 7 along with additional targets. Mean values and standard error of the mean (n=3) are shown. Cells were color coded for improved visualization with the value of 1 appearing white (no fold change compared to $Ph^{WT}$), lower expression levels in shades of blue and higher expression levels in shades of orange.

| | Kc | | $Ph^{WT}$ | | $Ph^{PHC3L}$ | | $Ph^{SAMsurf}$ | | | Kc | | $Ph^{WT}$ | | $Ph^{PHC3L}$ | | $Ph^{SAMsurf}$ | |
|---|---|---|---|---|---|---|---|---|---|---|---|---|---|---|---|---|---|
| **Ph** | Mean | SEM | Mean | SEM | Mean | SEM | Mean | SEM | **Psc** | Mean | SEM | Mean | SEM | Mean | SEM | Mean | SEM |
| ph-p promoter | 3.85 | 2.34 | 3.54 | 1.24 | 3.97 | 2.14 | 2.33 | 1.33 | | 1.28 | 0.58 | 0.58 | 0.24 | 0.63 | 0.23 | 0.23 | 0.11 |
| psc promoter | 1.19 | 0.73 | 1.57 | 0.56 | 0.8 | 0.47 | 1.36 | 0.61 | | 0.87 | 0.42 | 0.8 | 0.3 | 0.55 | 0.28 | 0.55 | 0.23 |
| psc upstream | 1.3 | 0.88 | 1.42 | 0.61 | 0.58 | 0.25 | 0.98 | 0.45 | | 0.91 | 0.45 | 0.63 | 0.23 | 0.49 | 0.24 | 0.38 | 0.19 |
| suz2 promoter | 1.67 | 0.93 | 1.4 | 0.4 | 0.73 | 0.43 | 0.96 | 0.39 | | 0.93 | 0.42 | 0.61 | 0.12 | 0.6 | 0.36 | 0.29 | 0.1 |
| zfh1 promoter | 0.85 | 0.43 | 1.32 | 0.71 | 0.8 | 0.4 | 1.46 | 0.8 | | 0.37 | 0.16 | 0.3 | 0.18 | 0.33 | 0.14 | 0.19 | 0.11 |
| zfh1 upstream | 0.87 | 0.32 | 1.01 | 0.57 | 0.29 | 0.11 | 0.62 | 0.32 | | 0.44 | 0.19 | 0.33 | 0.17 | 0.24 | 0.13 | 0.16 | 0.07 |
| zfh2 peak 1 | 2.11 | 1.41 | 0.89 | 0.45 | 0.6 | 0.29 | 0.66 | 0.37 | | 1.53 | 0.77 | 0.36 | 0.14 | 0.57 | 0.33 | 0.17 | 0.04 |
| zfh2 peak 2 | 0.47 | 0.32 | 1.43 | 0.62 | 1.01 | 0.6 | 1.28 | 0.63 | | 0.42 | 0.25 | 0.43 | 0.15 | 1.03 | 0.66 | 0.28 | 0.07 |
| chinmo promoter | 1.02 | 0.39 | 2.33 | 1.72 | 0.61 | 0.27 | 0.65 | 0.4 | | 0.36 | 0.17 | 0.23 | 0.09 | 0.55 | 0.4 | 0.25 | 0.05 |
| chinmo upstream | 13.78 | 7.79 | 19.58 | 8.27 | 5.13 | 2.44 | 7.26 | 4.04 | | 4.78 | 2.54 | 3.42 | 1.04 | 1.41 | 0.55 | 0.92 | 0.37 |
| cycb promoter | 0.16 | 0.08 | 0.49 | 0.35 | 0.21 | 0.12 | 0.28 | 0.17 | | 0.27 | 0.14 | 0.48 | 0.27 | 0.5 | 0.31 | 0.16 | 0.09 |
| tsh promoter | 1.61 | 0.76 | 2 | 1.03 | 1.14 | 0.61 | 1.72 | 0.86 | | 1.02 | 0.48 | 0.7 | 0.32 | 0.78 | 0.42 | 0.45 | 0.18 |
| pnr promoter | 3.06 | 1.11 | 1.4 | 0.75 | 0.99 | 0.52 | 1.04 | 0.66 | | 1.17 | 0.44 | 0.36 | 0.2 | 0.43 | 0.24 | 0.17 | 0.09 |
| socs36e promoter | 0.12 | 0.08 | 0.71 | 0.57 | 0.19 | 0.1 | 0.18 | 0.11 | | 0.23 | 0.15 | 0.36 | 0.24 | 0.36 | 0.23 | 0.1 | 0.04 |
| stat92e promoter | 0.29 | 0.13 | 0.77 | 0.51 | 0.5 | 0.28 | 0.46 | 0.3 | | 0.23 | 0.12 | 0.27 | 0.19 | 0.35 | 0.18 | 0.11 | 0.06 |
| upd1 upstream | 3.87 | 1.81 | 4.31 | 2.57 | 2.99 | 1.58 | 4.4 | 2.76 | | 2.09 | 0.98 | 1.72 | 0.74 | 2.04 | 1.1 | 1.36 | 0.77 |
| upd2 promoter | 1.08 | 0.44 | 1 | 0.51 | 0.76 | 0.26 | 0.87 | 0.37 | | 0.74 | 0.24 | 0.47 | 0.23 | 0.53 | 0.18 | 0.33 | 0.13 |
| upd3 promoter | 0.69 | 0.4 | 0.89 | 0.61 | 0.6 | 0.33 | 0.71 | 0.43 | | 0.43 | 0.25 | 0.33 | 0.2 | 0.46 | 0.29 | 0.17 | 0.09 |
| bxd | 4.18 | 3.25 | 4.48 | 2.04 | 4.09 | 2.52 | 3.14 | 1.68 | | 1.07 | 0.44 | 0.94 | 0.33 | 1.54 | 0.97 | 0.66 | 0.28 |
| cad | 4.28 | 1.8 | 5.72 | 2.06 | 5.52 | 2.32 | 3.62 | 1.5 | | 1.53 | 0.42 | 0.88 | 0.15 | 1.59 | 0.54 | 0.78 | 0.34 |
| neg. | 0.09 | 0.06 | 0.18 | 0.09 | 0.03 | 0.02 | 0.03 | 0.01 | | 0.18 | 0.07 | 0.25 | 0.15 | 0.25 | 0.19 | 0.1 | 0.02 |
| **GFP** | | | | | | | | | **IgG** | | | | | | | | |
| ph-p promoter | 0.04 | 0.03 | 0.26 | 0.09 | 0.02 | 0.01 | 0.12 | 0.06 | | 0.05 | 0.02 | 0.08 | 0.06 | 0.04 | 0.04 | 0.04 | 0.04 |
| psc promoter | 0.05 | 0.03 | 0.12 | 0.06 | 0.02 | 0 | 0.05 | 0.01 | | 0.06 | 0.03 | 0.11 | 0.09 | 0.02 | 0.01 | 0.01 | 0 |
| psc upstream | 0.04 | 0.03 | 0.14 | 0.1 | 0.03 | 0.01 | 0.02 | 0.01 | | 0.05 | 0.02 | 0.1 | 0.09 | 0.03 | 0.03 | 0.01 | 0 |
| suz2 promoter | 0.04 | 0.03 | 0.15 | 0.06 | 0.03 | 0.02 | 0.04 | 0.01 | | 0.05 | 0.02 | 0.09 | 0.07 | 0.03 | 0.01 | 0.02 | 0.01 |
| zfh1 promoter | 0.04 | 0.03 | 0.08 | 0.06 | 0.01 | 0.01 | 0.03 | 0.02 | | 0.04 | 0.02 | 0.1 | 0.08 | 0.02 | 0.01 | 0 | 0 |
| zfh1 upstream | 0.03 | 0.02 | 0.11 | 0.08 | 0.01 | 0 | 0.06 | 0.03 | | 0.03 | 0.01 | 0.07 | 0.06 | 0.02 | 0.01 | 0.01 | 0.01 |
| zfh2 peak 1 | 0.04 | 0.02 | 0.05 | 0.04 | 0 | 0 | 0.02 | 0.02 | | 0.06 | 0.03 | 0.11 | 0.09 | 0.02 | 0.01 | 0.05 | 0.04 |
| zfh2 peak 2 | 0.07 | 0.05 | 0.13 | 0.08 | 0.06 | 0.03 | 0.02 | 0 | | 0.05 | 0.01 | 0.12 | 0.1 | 0.05 | 0.04 | 0.02 | 0.01 |
| chinmo promoter | 0.01 | 0.01 | 0.28 | 0.24 | 0.03 | 0.02 | 0.04 | 0.02 | | 0.03 | 0.01 | 0.02 | 0.02 | 0.04 | 0.04 | 0 | 0 |
| chinmo upstream | 0.04 | 0.03 | 1.4 | 0.44 | 0.16 | 0.05 | 0.47 | 0.2 | | 0.05 | 0.02 | 0.11 | 0.07 | 0.02 | 0.02 | 0.01 | 0 |
| cycb promoter | 0.04 | 0.03 | 0.08 | 0.06 | 0 | 0 | 0.02 | 0.02 | | 0.06 | 0.03 | 0.14 | 0.1 | 0.04 | 0.03 | 0.02 | 0.02 |
| tsh promoter | 0.06 | 0.04 | 0.18 | 0.08 | 0.02 | 0.01 | 0.09 | 0.04 | | 0.07 | 0.02 | 0.13 | 0.11 | 0.03 | 0.02 | 0.03 | 0.02 |
| pnr promoter | 0.04 | 0.02 | 0.1 | 0.09 | 0.01 | 0.01 | 0.02 | 0.01 | | 0.04 | 0.02 | 0.12 | 0.09 | 0.02 | 0.02 | 0.03 | 0.02 |
| socs36e promoter | 0.05 | 0.03 | 0.14 | 0.12 | 0.01 | 0 | 0.01 | 0.01 | | 0.05 | 0.03 | 0.13 | 0.11 | 0.03 | 0.02 | 0.03 | 0.02 |
| stat92e promoter | 0.04 | 0.02 | 0.11 | 0.1 | 0.01 | 0 | 0.03 | 0.02 | | 0.03 | 0.01 | 0.11 | 0.09 | 0.02 | 0.01 | 0.02 | 0 |
| upd1 upstream | 0.05 | 0.03 | 0.25 | 0.18 | 0.08 | 0.04 | 0.17 | 0.11 | | 0.04 | 0.02 | 0.06 | 0.04 | 0.03 | 0.01 | 0.02 | 0.02 |
| upd2 promoter | 0.04 | 0.02 | 0.07 | 0.05 | 0.02 | 0 | 0.03 | 0.01 | | 0.04 | 0.02 | 0.09 | 0.08 | 0.03 | 0.02 | 0.02 | 0.01 |
| upd3 promoter | 0.04 | 0.03 | 0.1 | 0.07 | 0.01 | 0.01 | 0.02 | 0.01 | | 0.08 | 0.05 | 0.13 | 0.12 | 0.04 | 0.03 | 0.02 | 0.02 |
| bxd | 0.08 | 0.06 | 0.58 | 0.19 | 0.23 | 0.05 | 0.38 | 0.18 | | 0.09 | 0.04 | 0.21 | 0.19 | 0.03 | 0.02 | 0.09 | 0.07 |
| cad | 0.04 | 0.02 | 0.44 | 0.05 | 0.23 | 0.03 | 0.46 | 0.13 | | 0.05 | 0.02 | 0.07 | 0.05 | 0.02 | 0.01 | 0.02 | 0.01 |
| neg. | 0.04 | 0.02 | 0.04 | 0.03 | 0.02 | 0.02 | 0.03 | 0.01 | | 0.03 | 0.01 | 0.07 | 0.05 | 0.04 | 0.03 | 0.01 | 0.01 |

**Table S5: Summary of ChIP-qPCR data – related to Figure 7.** Overview of % Input data from ChIP-qPCR presented in Fig 7 along with additional targets. Mean values and standard error of the mean (n=3) are shown for Ph-ChIP (top left), Psc-ChIP (top right), GFP (bottom left) and IgG (bottom right). Cells were color coded for improved visualization. The median of all means for a given data set appears as white, lower signals in shades of blue and higher signals in shades of orange. For IgG, the color code of the Ph dataset was applied for better comparison.

| Name | | Sequence | Note |
|---|---|---|---|
| **dsDNA template** | | ACGCGTCGGTGTTAGAGCCTGTAACTCGGTGTTAGAGCCTGTA<br>ACTAGTCTGCAGGTTACCTAGGACTTCAGGCACCCAGGCTTCG<br>TACATCAAGCTTCATCGGGGGCTATAAAAGGGGGTTCCTCGAG<br>GTTCCCTGGGACACTGTCTAGAGAATTC | Phase separation assays, EMSA |
| **Genotyping primers for endogenous *ph* edits** | **ph-p/ph-d (616)**<br>**ph-p (2012)**<br>**ph-d (2013)**<br>**ph-p/ph-d (2057)**<br>**Venus (2011)** | CGATATCCTAGGAATCAACAGCACCACCAATCCGGGCAAC<br>GGCAGTGACATTGGAGGTAC<br>TTGGCAAAGCGTCCGATACTG<br>CACAGGGAAGGACGGTGCAGCTAATG<br>TCTCGTTGGGGTCTTTGCTC | For PCR amplification |
| **aTub** | | TCGCGTGTGAAACACTTCCA<br>CAGCAGGCGTTTCCAATCTG | For RT-qPCR |
| **ph-p** | | CAAAACGCCTAAGCCGACG<br>GCTTTCTGTGTCCGCTCTTTT | For RT-qPCR |
| **ph-d** | | TAGTTTCCCGCGTTTCACCT<br>GTGGTGAGTCGGAACGTGAT | For RT-qPCR |
| **psc** | | TTCTAACGGCCGTCAATCCG<br>AACTGTGGCAGAAGGAGTGC | For RT-qPCR |
| **suz2** | | CGCGGAAAAATCGTACCTCC<br>GCCTCTTTCAATGGGGTCAAAA | For RT-qPCR |
| **chinmo** | | TGAGTGCGCGAATCCAACTA<br>AACACGCCAAATGCTGCTTG | For RT-qPCR |
| **cycb** | | GCGTCTGCCTATCTTCGTGT<br>AAGTTCTCCGAAGCGTTCTCA | For RT-qPCR |
| **tsh** | | CAAGTTCCAGCAAACCGTCAA<br>GGGCAGAGCTCCGGCATT | For RT-qPCR |
| **zfh1** | | CGATCCCGATCTGCCCTATG<br>GTAGGGTCGTTGACCGGAAT | For RT-qPCR |
| **zfh2** | | CTAACGCAAACTCTCAGCAGC<br>TGGGCCCGCAATATTTTAAGC | For RT-qPCR |
| **bbg** | | GAGATTCGTGCACAAGCGGA<br>GCGAATCAAAGCCTCCTTCG | For RT-qPCR |
| **ptc** | | CGGAGCTTCTCAGGGCAAAT<br>CCGTGGAGGTAAAAGGGCAT | For RT-qPCR |
| **pask** | | TGCGTTAATGAAGCCAAGCAC<br>GACCCTCAGCTCGTGGTTAC | For RT-qPCR |
| **thor** | | ATCCAGATGCCCGAGGTGTA<br>CGTAGATAAGTTTGGTGCCTCC | For RT-qPCR |
| **sans** | | ACCGGGACAACTAACTGGTA<br>ATCTTTGGCGGCTTTGTGAA | For RT-qPCR |
| **cg44251** | | TAACGCACAATGGCCTACAA<br>GCAGACCAACATATCGCCCT | For RT-qPCR |
| **acxd** | | CCACACATGTTGCCGACTTC<br>AATCTCATTGAGCACCCGCA | For RT-qPCR |
| **pnr** | | CGCCTGGTGAGTGCAACTG<br>CAAAGTGTGGTGGTCCGAGT | For RT-qPCR |
| **socs36e** | | CCAAAAGTAACGAGCACACCG<br>GGACCTCCGATTGTTTTCTCT | For RT-qPCR |
| **stat92e** | | ACCGGTTATGTGAAGAGCACA<br>TCCAAAGTCACCATTTCCCAG | For RT-qPCR |
| **upd2** | | CGAGTACAAGTTCCTGCCGA<br>CAAGTCTTTAGCTTCACCGCAC | For RT-qPCR |
| **upd3** | | CTGAACGAAACGCACAGCAA<br>ATTCTGCAGGATCCTTTGGC | For RT-qPCR |
| **mir_8** | | TTTCTTGCCCCAGTGGGTTT<br>GGCTAGACCGAGACTAACGC | For RT-qPCR |
| **mir_bantam** | | CTGTCATCCCAAGGCGAGTT<br>CGCTGGTATGCTTGGAATGC | For RT-qPCR |

| | | |
|---|---|---|
| **mir_184** | ACGTTTTACGGGGCAGAGAG<br>AACAAAACGCGGAGCAACTC | For RT-qPCR |
| **ph-p promoter** | GCATGTGCGCTGAATATGTG<br>ACCCGTGCATTTCGGTATAG | For ChIP-qPCR |
| **psc promoter** | TTCGCGGGCACCTTTATT<br>AACTACGTCTGCTGGCTTATG | For ChIP-qPCR |
| **psc upstream** | CAAGCCACATCCAGTACATACA<br>CCTTCTCCATTCCACACATCTT | For ChIP-qPCR |
| **suz2 promoter** | CGATAGAGAGTGGGAGAGAAGT<br>GCTTGAAAGTGTCGGCAATC | For ChIP-qPCR |
| **zfh1 promoter** | GCCCGAATCACCTCGTACAT<br>CTGTTTGTGTGTGTGCTCCG | For ChIP-qPCR |
| **zfh1 upstream** | CAACAGGTGCAACCGATGGA<br>AGGTGGCGTGGGTGTTTAAT | For ChIP-qPCR |
| **zfh2 peak1** | GGTCGGACTTTCGGAGAATAAA<br>GAGAGGCAGGCTTACTTGTT | For ChIP-qPCR |
| **zfh2 peak2** | ACGAGAAAGAGGCCGAAATG<br>GAGGCGCTGATGGGTAAATAG | For ChIP-qPCR |
| **chinmo promoter** | TTCCGACTGGGGAAGCAAAA<br>CAAGAAACACGTGGAGTGCC | For ChIP-qPCR |
| **chinmo upstream** | CAATACACACGCGCGATCAG<br>TTTGCACACACCCGCTATCT | For ChIP-qPCR |
| **cycb promoter** | AGAGATGCGATACGTGCGAT<br>GGTAGCACTGTTTCAGCCCT | For ChIP-qPCR |
| **tsh promoter** | AGGCTAAAAGTGACGAGGGC<br>ACTCAAACTGTGTCGGTCCC | For ChIP-qPCR |
| **pnr promoter** | GGCTTAAACTGCGAAGGGGA<br>TCTGGGGACAGCCCACTTT | For ChIP-qPCR |
| **socs36e promoter** | AACCCGTGGCATCATTAGCA<br>TCTAAGCATTAGCGGGGCTG | For ChIP-qPCR |
| **stat92e promoter** | GCCGAATGGGCGCATTTTAG<br>AGTAACGATTGCCTGGTGCT | For ChIP-qPCR |
| **upd1 upstream** | TAGGCCCATCGGTGAAACTG<br>CGTCTTCGGAGTGACGCTTA | For ChIP-qPCR |
| **upd2 promoter** | TCCGCACCGACCAATAACAA<br>CTCAGTCGGCGTCGTTATCA | For ChIP-qPCR |
| **upd3 promoter** | CGCTTCTTTGGCGATTCGAG<br>GGCGGAGACTGACTGAACTC | For ChIP-qPCR |
| **bxd** | CCATAAGAAATGCCACTTTGC<br>CTCTCACTCTCTCACTGTGAT | For ChIP-qPCR |
| **cad** | AGGAGTGTGTAGAGCCTGTG<br>CCCCCACGCCTCTAAACAAA | For ChIP-qPCR |
| **neg.** | TCAAGCCGAACCCTCTAAAAT<br>AACGCCAACAAACAGAAAATG | For ChIP-qPCR |

**Table S6: Nucleic acid and oligonucleotide sequences – related to Methods.** All primers were purchased from IDT.

| Protein | Sequence | Note | Figures |
|---|---|---|---|
| **mini-Ph$^{WT}$** | MDYKDDDDKGSNGSKDLPKAMIKPNVLTHVIDGFIIQEANEPFPVTRQRYADKDVSDEPPKKKATMQEDIKLSGIASAPGSDMVACEQCGKMEHKAKLKRKRYCSPGCSRQAKNGIGGVGSGETNGLGTGGIVGVDAMALVDRLDEAMAEEKMQTEATPKLSESFPILGASTEVPPMSLPVQAAISAPSPLAMPLGSPLSVALPTLAPLSVVTSGAAPKSSEVNGTDRPPISSWSVDDVSNFIRELPGCQDYVDDFIQQEIDGQALLLLKEKHLVNAMGMKLGPALKIVAKVESIKEVPPPGEAKDPGAQ | *in vitro* experiments, purified from Sf9 cells; Ext. coefficient: 11460 $M^{-1}cm^{-1}$ | 1; 4; S1; S2; S4 |
| **mini-Ph$^{PHC3L}$** | MDYKDDDDKGSNGSKDLPKAMIKPNVLTHVIDGFIIQEANEPFPVTRQRYADKDVSDEPPKKKATMQEDIKLSGIASAPGSDMVACEQCGKMEHKAKLKRKRYCSPGCSRQAKNGIG**SKKFALSRWNRKPDNQSLGHRGRRPSGPDGAAREHILRQLPITYPSAEEDLASHEDSVPSAMTTRLRRQSERERERELRDVRIRKMPENSDLLPVAQ**PPISSWSVDDVSNFIRELPGCQDYVDDFIQQEIDGQALLLLKEKHLVNAMGMKLGPALKIVAKVESIKEVPPPGEAKDPGAQ | *in vitro* experiments, purified from Sf9 cells; Ext. coefficient: 18450 $M^{-1}cm^{-1}$ | 1; 4; S1; S2; S4 |
| **mini-Ph$^{SAMsurf}$** | MDYKDDDDKGSNGSKDLPKAMIKPNVLTHVIDGFIIQEANEPFPVTRQRYADKDVSDEPPKKKATMQEDIKLSGIASAPGSDMVACEQCGKMEHKAKLKRKRYCSPGCSRQAKNGIGGVGSGETNGLGTGGIVGVDAMALVDRLDEAMAEEKMQTEATPKLSESFPILGASTEVPPMSLPVQAAISAPSPLAMPLGSPLSVALPTLAPLSVVTSGAAPKSSEVNGTDRPPISSWSV**KR**VSNFIRELPGCQDYV**KR**FIQQEIDGQALLLLKEKHLVNAMGMKLGPALKIVAKVESIKEVPPPGEAKDPGAQ | *in vitro* experiments, purified from Sf9 cells; Ext. coefficient: 11460 $M^{-1}cm^{-1}$ | 1; 4; S1; S2; S4 |
| **mini-Ph$^{EH}$** | MEKTRNGSKDLPKAMIKPNVLTHVIDGFIIQEANEPFPVTRQRYADKDVSDEPPKKKATMQEDIKLSGIASAPGSDMVACEQCGKMEHKAKLKRKRYCSPGCSRQAKNGIGGVGSGETNGLGTGGIVGVDAMALVDRLDEAMAEEKMQTEATPKLSESFPILGASTEVPPMSLPVQAAISAPSPLAMPLGSPLSVALPTLAPLSVVTSGAAPKSSEVNGTDRPPISSWSVDDVSNFIRELPGCQDYVDDFIQQEIDGQALLLLKEKHLVNAMGMKLGPA**R**KIVAKVESIKEVRDHHHHHH | *in vitro* experiments, purified from E. coli; Ext. coefficient: 9970 $M^{-1}cm^{-1}$ | 4; S1; S2; S4 |
| **mini-Ph$^{PHC3L-EH}$** | MEKTRNGSKDLPKAMIKPNVLTHVIDGFIIQEANEPFPVTRQRYADKDVSDEPPKKKATMQEDIKLSGIASAPGSDMVACEQCGKMEHKAKLKRKRYCSPGCSRQAKNGIG**SKKFALSRWNRKPDNQSLGHRGRRPSGPDGAAREHILRQLPITYPSAEEDLASHEDSVPSAMTTRLRRQSERERERELRDVRIRKMPENSDLLPVAQ**PPISSWSVDDVSNFIRELPGCQDYVDDFIQQEIDGQALLLLKEKHLVNAMGMKLGPA**R**KIVAKVESIKEVRDHHHHHH | *in vitro* experiments, purified from E. coli; Ext. coefficient: 16960 $M^{-1}cm^{-1}$ | 4; S1; S2; S4 |
| **mini-Ph$^{SAMsurf-EH}$** | MEKTRNGSKDLPKAMIKPNVLTHVIDGFIIQEANEPFPVTRQRYADKDVSDEPPKKKATMQEDIKLSGIASAPGSDMVACEQCGKMEHKAKLKRKRYCSPGCSRQAKNGIGGVGSGETNGLGTGGIVGVDAMALVDRLDEAMAEEKMQTEATPKLSESFPILGASTEVPPMSLPVQAAISAPSPLAMPLGSPLSVALPTLAPLSVVTSGAAPKSSEVNGTDRPPISSWSV**KR**VSNFIRELPGCQDYV**KR**FIQQEIDGQALLLLKEKHLVNAMGMKLGPA**R**KIVAKVESIKEVRDHHHHHH | *in vitro* experiments, purified from E. coli; Ext. coefficient: 9970 $M^{-1}cm^{-1}$ | 4; S1; S2; S4 |
| **HisSUMO-PhL-SAM** | MGSSHHHHHHGSGLVPRGSASMSDSEVNQEAKPEVKPEVKPETHINLKVSDGSSEIFFKIKKTTPLRRLMEAFAKRQGKEMDSLRFLYDGIRIQADQTPEDLDMEDNDIIEAHREQIGGGVGSGETNGLGTGGIVGVDAMALVDRLDEAMAEEKMQTEATPKLSESFPILGASTEVPPMSLPVQAAISAPSPLAMPLGSPLSVALPTLAPLSVVTSGAAPKSSEVNGTDRPPISSWSVDDVSNFIRELPGCQDYVDDFIQQEIDGQALLLLKEKHLVNAMGMKLGPALKIVAKVESIKEVPPPGEAKDPGAQ | *in vitro* experiments, purified from E. coli | 2; S2; S3 |
| **HisSUMO-PHC3L-SAM** | MGSSHHHHHHGSGLVPRGSASMSDSEVNQEAKPEVKPEVKPETHINLKVSDGSSEIFFKIKKTTPLRRLMEAFAKRQGKEMDSLRFLYDGIRIQADQTPEDLDMEDNDIIEAHREQIGG**SKKFALSRWNRKPDNQSLGHRGRRPSGPDGAAREHILRQLPITYPSAEEDLASHEDSVPSAMTTRLRRQSERERERELRDVRIRKMPENSDLLPVAQ**PPISSWSVDDVSNFIRELPGCQDYVDDFIQQEIDGQALLLLKEKHLVNAMGMKLGPALKIVAKVESIKEVPPPGEAKDPGAQ | *in vitro* experiments, purified from E. coli | 2; S2; S3 |
| **HisSUMO-PhL-SAMsurf** | MGSSHHHHHHGSGLVPRGSASMSDSEVNQEAKPEVKPEVKPETHINLKVSDGSSEIFFKIKKTTPLRRLMEAFAKRQGKEMDSLRFLYDGIRIQADQTPEDLDMEDNDIIEAHREQIGGGVGSGETNGLGTGGIVGVDAMALVDRLDEAMAEEKMQTEATPKLSESFPILGASTEVPPMSLPVQAAISAPSPLAMPLGSPLSVALPTLAPLSVVTSGAAPKSSEVNGTDRPPISSWSV**KR**VSNFIRELPGCQDYV**KR**FIQQEIDGQALLLLKEKHLVNAMGMKLGPALKIVAKVESIKEVPPPGEAKDPGAQ | *in vitro* experiments, purified from E. coli | S3 |
| **PhL-SAM variants (here WT, mutant SAM variants below)** | MHHHHHHAMKGVDSPSAELDKKAENLYFQGTRGVGSGETNGLGTGGIVGVDAMALVDRLDEAMAEEKMQTEATPKLSESFPILGASTEVPPMSLPVQAAISAPSPLAMPLGSPLSVALPTLAPLSVVTSGAAPKSSEVNGTDRPPISSWSVDDVSNFIRELPGCQDYVDDFIQQEIDGQALLLLKEKHLVNAMGMKLGPALKIVAKVESIKEV | AUC experiments | 3 |
| **PhL-SAM** | GVGSGETNGLGTGGIVGVDAMALVDRLDEAMAEEKMQTEATPKLSESFPILGASTEVPPMSLPVQAAISAPSPLAMPLGSPLSVALPTLAPLSVVTSGAAPKSSEVNGTDRPPISSWSVDDVSNFIRELPGCQDYVDDFIQQEIDGQALLLLKEKHLVNAMGMKLGPALKIVAKVESIKEVPPPGEAKDPGAQ | Simulations | 2; S3 |
| **PHC3L-SAM** | **RYNVSCSKKFALSRWNRKPDNQSLGHRGRRPSGPDGAAREHILRQLPITYPSAEEDLASHEDSVPSAMTTRLRRQSERERERELRDVRIRKMPENSDLLPVAQ**PPISSWSVDDVSNFIRELPGCQDYVDDFIQQEIDGQALLLLKEKHLVNAMGMKLGPALKIVAKVESIKEVPPPGEAKDPGAQ | Simulations | 2; S2 |
| **PhL-SAMsurf (similar for individual 16/17 and 33/34 mutants)** | GVGSGETNGLGTGGIVGVDAMALVDRLDEAMAEEKMQTEATPKLSESFPILGASTEVPPMSLPVQAAISAPSPLAMPLGSPLSVALPTLAPLSVVTSGAAPKSSEVNGTDRPPISSWSV**KR**VSNFIRELPGCQDYV**KR**FIQQEIDGQALLLLKEKHLVNAMGMKLGPALKIVAKVESIKEVPPPGEAKDPGAQ | Simulations | 3; S3 |
| **PHC1L-PHC1SAM** | SCSHQFRLKRKKMKEFQEANYARVRRRGPRRSSSDIARAKIQGKCHRGQEDSSRGSDNSSYDEALSPTSPGPLSVRAGHGERDLGNPNTAPPTPELHGINPVFLSWSVEEVYEFIASLQGCQEIAEEFRSQEIDGQALLLLKEEHLMSAMNIKLGPALKICAKINVLKET | Simulations | 2; S2 |
| **PhL-PHC1SAM** | **GVGSGETNGLGTGGIVGVDAMALVDRLDEAMAEEKMQTEATPKLSESFPILGASTEVPPMSLPVQAAISAPSPLAMPLGSPLSVALPTLAPLSVVTSGAAPKSSEVNGTDR**WSVEEVYEFIASLQGCQEIAEEFRSQEIDGQALLLLKEEHLMSAMNIKLGPALKICAKINVLKET | Simulations | 2; S2 |
| **PHC2L-PHC2SAM** | TKRVGLFHSDRSKLQKAGAATHNRRRASKASLPPLTKDTKKQPTGTVPLSVTAALQLTHSQEDSSRCSDNSSYEEPLSPISASSSTSRRRQGQRDLELPDMHMRDLVGMGHHFLPSWNVEDVYEFIRSLPGCQEIAEEFRAQEIDGQALLLLKEDHLMSAMNIKLGPALKIYARISMLKDS | Simulations | 2; S2 |
| **PhL-PHC2SAM** | **GVGSGETNGLGTGGIVGVDAMALVDRLDEAMAEEKMQTEATPKLSESFPILGASTEVPPMSLPVQAAISAPSPLAMPLGSPLSVALPTLAPLSVVTSGAAPKSSEVNGTDR**WNVEDVYEFIRSLPGCQEIAEEFRAQEIDGQALLLLKEDHLMSAMNIKLGPALKIYARISMLKDS | Simulations | 2; S2 |
| **PHC3L-PHC3SAM** | SKKFALSRWNRKPDNQSLGHRGRRPSGPDGAAREHILRQLPITYPSAEEDLASHEDSVPSAMTTRLRRQSERERERELRDVRIRKMPENSDLLPVAQTEPSIWTVDDVWAFIHSLPGCQDIADEFRAQEIDGQALLLLKEDHLMSAMNIKLGPALKICARINSLKES | Simulations | 2; S2 |
| **PhL-PHC3SAM** | **GVGSGETNGLGTGGIVGVDAMALVDRLDEAMAEEKMQTEATPKLSESFPILGASTEVPPMSLPVQAAISAPSPLAMPLGSPLSVALPTLAPLSVVTSGAAPKSSEVNGTDR**TEPSIWTVDDVWAFIHSLPGCQDIADEFRAQEIDGQALLLLKEDHLMSAMNIKLGPALKICARINSLKES | Simulations | 2; S2 |
| **PhL** | GVGSGETNGLGTGGIVGVDAMALVDRLDEAMAEEKMQTEATPKLSESFPILGASTEVPPMSLPVQAAISAPSPLAMPLGSPLSVALPTLAPLSVVTSGAAPKSSEVNGTDR | Simulations | 2 |
| **PHC1L** | SCSHQFRLKRKKMKEFQEANYARVRRRGPRRSSSDIARAKIQGKCHRGQEDSSRGSDNSSYDEALSPTSPGPLSVRAGHGERDLGNPNTAPPTPELHGINPVFLS | Simulations | 2 |
| **PHC2L** | TKRVGLFHSDRSKLQKAGAATHNRRRASKASLPPLTKDTKKQPTGTVPLSVTAALQLTHSQEDSSRCSDNSSYEEPLSPISASSSTSRRRQGQRDLELPDMHMRDLVGMGHHFLPS | Simulations | 2 |
| **PHC3L** | RYNVSCSKKFALSRWNRKPDNQSLGHRGRRPSGPDGAAREHILRQLPITYPSAEEDLASHEDSVPSAMTTRLRRQSERERERELRDVRIRKMPENSDLLPVAQ | Simulations | 2 |
| **PhSAM WT** | PPISSWSVDDVSNFIRELPGCQDYVDDFIQQEIDGQALLLLKEKHLVNAMGMKLGPALKIVAKVESIKEVPPPGEAKDPGAQ | Simulations | 3; S3 |
| **SAM D16K/D17R** | PPISSWSV**KR**VSNFIRELPGCQDYVDDFIQQEIDGQALLLLKEKHLVNAMGMKLGPALKIVAKVESIKEVPPPGEAKDPGAQ | Simulations (and AUC) | 3; S3 |
| **SAM D33K/D34R** | PPISSWSVDDVSNFIRELPGCQDYV**KR**FIQQEIDGQALLLLKEKHLVNAMGMKLGPALKIVAKVESIKEVPPPGEAKDPGAQ | Simulations (and AUC) | 3; S3 |
| **SAMsurf** | PPISSWSV**KR**VSNFIRELPGCQDYV**KR**FIQQEIDGQALLLLKEKHLVNAMGMKLGPALKIVAKVESIKEVPPPGEAKDPGAQ | Simulations (and AUC) | 3; S3 |
| **SAMsurf EH** | PPISSWSV**KR**VSNFIRELPGCQDYV**KR**FIQQEIDGQALLLLKEKHLVNAMGMKLGPA**R**KIVAKVESIKEVPPPGEAKDPGAQ | Simulations (and AUC) | 3; S3 |
| **Ph-p full-length** | Uniprot: P39769 | Simulations | S5 |
| **Mini Ph WT** | ISNGSKDLPKAMIKPNVLTHVIDGFIIQEANEPFPVTRQRYADKDVSDEPPKKKATMQEDIKLSGIASAPGSDMVACEQCGKMEHKAKLKRKRYCSPGCSRQAKNGIGGVGSGETNGLGTGGIVGVDAMALVDRLDEAMAEEKMQTEATPKLSESFPILGASTEVPPMSLPVQAAISAPSPLAMPLGSPLSVALPTLAPLSVVTSGAAPKSSEVNGTDRPPISSWSVDDVSNFIRELPGCQDYVDDFIQQEIDGQALLLLKEKHLVNAMGMKLGPALKIVAKVESIKEV | Simulations | 5; S5 |
| **Mini Ph PHC3L** | ISNGSKDLPKAMIKPNVLTHVIDGFIIQEANEPFPVTRQRYADKDVSDEPPKKKATMQEDIKLSGIASAPGSDMVACEQCGKMEHKAKLKRKRYCSPGCSRQAKNGIGRYNVSC**SKKFALSRWNRKPDNQSLGHRGRRPSGPDGAAREHILRQLPITYPSAEEDLASHEDSVPSAMTTRLRRQSERERERELRDVRIRKMPENSDLLPVAQTEPSI**PPISSWSVDDVSNFIRELPGCQDYVDDFIQQEIDGQALLLLKEKHLVNAMGMKLGPALKIVAKVESIKEV | Simulations | 5; S5 |
| **Mini Ph SAMsurf** | ISNGSKDLPKAMIKPNVLTHVIDGFIIQEANEPFPVTRQRYADKDVSDEPPKKKATMQEDIKLSGIASAPGSDMVACEQCGKMEHKAKLKRKRYCSPGCSRQAKNGIGGVGSGETNGLGTGGIVGVDAMALVDRLDEAMAEEKMQTEATPKLSESFPILGASTEVPPMSLPVQAAISAPSPLAMPLGSPLSVALPTLAPLSVVTSGAAPKSSEVNGTDRPPISSWSV**KR**VSNFIRELPGCQDYV**KR**FIQQEIDGQALLLLKEKHLVNAMGMKLGPALKIVAKVESIKEV | Simulations | 5; S5 |
| **mPHC3** | KPPQAIVKPQILTHVIEGFVIQEGLEPFPVSRSSLLIEQPVKKRPLLDNQVINSVCVQPELQNNTKHADNSSDTEMEDMIAEETLEEMDSELLKCEFCGKMGYANEFLRSKRFCTMSCAKRYNVSCSKKFALSRWNRKPDNQSLGHRGRRPSGPDGAAREHILRQLPITYPSAEEDLASHEDSVPSAMTTRLRRQSERERERELRDVRIRKMPENSDLLPVAQTEPSIWTVDDVWAFIHSLPGCQDIADEFRAQEIDGQALLLLKEDHLMSAMNIKLGPALKICARINSLKES | Simulations | 5; S5 |

| | | | |
|---|---|---|---|
| **mini-Ph variants (here WT)** | MVSKGEELFTGVVPILVELDGDVNGHKFSVSGEGEGDATYGKLTLKLICTTGKLPVPWPTLVTTLGYGLQCFARYPDHMKQHDFFKSAMPEGYVQERTIFFKDDGNYKT<br>RAEVKFEGDTLVNRIELKGIDFKEDGNILGHKLEYNYNSHNVYITADKQKNGIKANFKIRHNIEDGGVQLADHYQQNTPIGDGPVLLPDNHYLSYQSALSKDPNEKRDHM<br>VLLEFVTAAGITLGMDELYKHRYTSLYKKAGSEFHVRGRSATRNMIKPNVLTHVIDGFIIQEANEPFPVTRQRYADKDVSDEPPKKKATMQEDIKLSGIASAPGSDMVAC<br>EQCGKMEHKAKLKRKRYCSPGCSRQAKNGIGGVGSGETNGLGTGGIVGVDAMALVDRLDEAMAEEKMQTEATPKLSESFPILGASTEVPPMSLPVQAAISAPSPLAMP<br>LGSPLSVALPTLAPLSVVTSGAAPKSSEVNGTDRPPISSWSVDDVSNFIRELPGCQDYVDDFIQQEIDGQALLLLKEKHLVNAMGMKLGPALKIVAKVESIKEVPPPGEA<br>KDPGAQ | Cell experiments | 6; S6; S7 |
| **Ph variants (here WT)** | MVSKGEELFTGVVPILVELDGDVNGHKFSVSGEGEGDATYGKLTLKLICTTGKLPVPWPTLVTTLGYGLQCFARYPDHMKQHDFFKSAMPEGYVQERTIFFKDDGNYKT<br>RAEVKFEGDTLVNRIELKGIDFKEDGNILGHKLEYNYNSHNVYITADKQKNGIKANFKIRHNIEDGGVQLADHYQQNTPIGDGPVLLPDNHYLSYQSALSKDPNEKRDHM<br>VLLEFVTAAGITLGMDELYKHRYTSLYKKAGSEFHMDRRALKFMQKRADTESDTTTPVSTTASQGISASAILAGGTLPLKDNSNIREKPLHHNYNHNNNNSSQHSHSH<br>QQQQQQQVGGKQLERPLKCLETLAQKAGITFDEKYDVASPPHPGIAQQQATSGTGPATGSGSVTPTSHRHGTPPTGRRQTHTPSTPNRPSAPSTPNTNCNSIARHTSLT<br>LEKAQNPGQQVAATTTVPLQISPEQLQQFYASNPYAIQVKQEFPTHTTSGSGTELKHATNIMEVQQQLQLQQQLSEANGGGAASAGAGGAASPANSQQSQQQQHSTAIST<br>MSPMQLAAATGGVGGDWTQGRTVQLMQPSTSFLYPQMIVSGNLLHPGGLGQQPIQVITAGKPFQGNGPQMLTTTTQNAKQMIGGQAGFAGGNYATCIPTNHNQSPQTVL<br>FSPMNVISPQQQQNLLQSMAAAAQQQQLTQQQQQFNQQQQQQLTQQQQQLTAALAKVGVDAQGKLAQKVVQKVTTTSSAVQAATGPGSTGSTQTQQVQQVQQQQQQTTQ<br>TTQQCVQVSQSTLPVGVGGQSVQTAQLLNAGQAQQMQIPWFLQNAAGLQPFGPNQIILRNQPDGTQGMFIQQQPATQTLQTQQNQIIQCNVTQTPTKARTQLDALAPKQQQ<br>QQQQVGTTNQTQQQQLAVATAQLQQQQQQLTAAALQRPGAPVMPHNGTQVRPASSVSTQTAQNQSLLKAKMRNKQQPVRPALATLKTEIGQVAGQNKVVGHLTTVQQQ<br>QQATNLQQVVNAAGNKMVVMSTTGTPITLQNGQTLHAATAAGVDKQQQQLQLFQKQQILQQQQMLQQQIAAIQMQQQQAAVQAQQQQQQQVSQQQQVNAQQQQAVAQQQ<br>QAVAQAQQQQREQQQQVAQAQAQHQQALANATQQILQVAPNQFITSHQQQQQQQLHNQLIQQQLQQQAQAQVQAQVQAQAQQQQQQREQQQNIIQQIVVQQSGATSQQTSQ<br>QQQHHQSGQLQLSSVPFSVSSSTTPAGIATSSALQAALSASGAIFQTAKPGTCSSSSPTSSVVTITNQSSTPLVTSSTVASIQQAQTQSAQVHQHQQLISATIAGGTQQQPQ<br>GPPSLTPTTNPILAMTSMMNATVGHLSTAPPVTVSVTSTAVTSSPGQLVLLSTASSGGGGSIPATPTKETPSKGPTATLVPIGSPKTPVSGKDTCTTPKSSTPATVSASV<br>EASSSTGEALSNGDASDRSSTPSKGATTPTSKQSNAAVQPPSSTTPNSVSGKEEPKLATCGSLTSATSTSTTTTITNGIGVARTTASTAVSTASTTTTSSGTFITSCTST<br>TTTTTSSISNGSKDLPKAMIKPNVLTHVIDGFIIQEANEPFPVTRQRYADKDVSDEPPKKKATMQEDIKLSGIASAPGSDMVACEQCGKMEHKAKLKRKRYCSPGCSRQA<br>KNGIGGVGSGETNGLGTGGIVGVDAMALVDRLDEAMAEEKMQTEATPKLSESFPILGASTEVPPMSLPVQAAISAPSPLAMPLGSPLSVALPTLAPLSVVTSGAAPKSS<br>EVNGTDRPPISSWSVDDVSNFIRELPGCQDYVDDFIQQEIDGQALLLLKEKHLVNAMGMKLGPALKIVAKVESIKEVPPPGEAKDPGAQ | Cell experiments (except endogenous edits) and fly experiments | 6; 7; S6; S7 |
| **PHC3 WT** | MVSKGEELFTGVVPILVELDGDVNGHKFSVSGEGEGDATYGKLTLKLICTTGKLPVPWPTLVTTLGYGLQCFARYPDHMKQHDFFKSAMPEGYVQERTIFFKDDGNYKT<br>RAEVKFEGDTLVNRIELKGIDFKEDGNILGHKLEYNYNSHNVYITADKQKNGIKANFKIRHNIEDGGVQLADHYQQNTPIGDGPVLLPDNHYLSYQSALSKDPNEKRDHM<br>VLLEFVTAAGITLGMDELYKHRYIDYKDDDDKTSLYKKAGSTMDTEPNPGTSSVSTTTSSTTTTTITTSSSRMQQPQISVYSGSDRHAVQVIQQALHRPPSSAAQYLQQMY<br>AAQQQHLMLHTAALQQQHLSSSQLQSLAAVQASLSSGRPSTSPTGSVTQQSSMSQTSINLSTSPTPAQLISRSQASSSTSGSITQQTMLLGSTSPTLTASQAQMYLRAQM<br>LIFTPATTVAAVQSDIPVVSSSSSSSCQSAATQVQNLTLRSQKLGVLSSSQNGPPKSTSQTQSLTICHNKTTVTSSKISQRDPSPESNKKGESPSLESRSTAVTRTSSIHQ<br>LIAPASYSPIQPHSLIKHQRIPLHSPPSKVSHHQLILQQQQQIQPITLQNSTQDPPPSQHCIPLQNHGLPPAPSNAQSQHCSPIQSHPSPLTVSPNQSQSAQQSVVVSPPP<br>PHSPSQSPTIIIHPQALIQPHPLVSSALQPGPDLQQSTANQVQATAQLNLPSHLPLPASPVVHIGPVQQSALVSPGQQIVSPSHQQYSSLQSSPIPIASPPQMSTSPPAQIPP<br>LPLQSMQSLQVQPEILSQGQVLVQNALVSEEELPAAEALVQLPFQTLPPPQTVAVNLQVQPPAPVDPPVVYQVEDVCEEEMPEESDECVRMDRTPPPPTLSPAAITVGR<br>GEDLTSEHPLLEQVELPAVASVSASVIKSPSDPSHVSVPPPPLLLPAATTRSNSTSMHSSIPSIENKPPQAIVKPQILTHVIEGFVIQEGLEPFPVSRSSLLIEQPVKKRPL<br>LDNQVINSVCVQPELQNNTKHADNSSDTEMEDMIAEETLEEMDSELLKCEFCGKMGYANEFLRSKRFCTMSCAKRYNVSCSKKFALSRWNRKPDNQSLGHRGRRPS<br>GPDGAAREHILRQLPITYPSAEEDLASHEDSVPSAMTTRLRRQSERERERELRDVRIRKMPENSDLLPVAQTEPSIWTVDDVWAFIHSLPGCQDIADEFRAQEIDGQALL<br>LLKEDHLMSAMNIKLGPALKICARINSLKES | Cell experiments | 6; S6; S7 |
| **PHC3 PhL** | MVSKGEELFTGVVPILVELDGDVNGHKFSVSGEGEGDATYGKLTLKLICTTGKLPVPWPTLVTTLGYGLQCFARYPDHMKQHDFFKSAMPEGYVQERTIFFKDDGNYKT<br>RAEVKFEGDTLVNRIELKGIDFKEDGNILGHKLEYNYNSHNVYITADKQKNGIKANFKIRHNIEDGGVQLADHYQQNTPIGDGPVLLPDNHYLSYQSALSKDPNEKRDHM<br>VLLEFVTAAGITLGMDELYKHRYTSLYKKAGSTMDTEPNPGTSSVSTTTSSTTTTTITTSSSRMQQPQISVYSGSDRHAVQVIQQALHRPPSSAAQYLQQMYAAQQQHLM<br>LHTAALQQQHLSSSQLQSLAAVQASLSSGRPSTSPTGSVTQQSSMSQTSINLSTSPTPAQLISRSQASSSTSGSITQQTMLLGSTSPTLTASQAQMYLRAQMLIFTPATTV<br>AAVQSDIPVVSSSSSSSCQSAATQVQNLTLRSQKLGVLSSSQNGPPKSTSQTQSLTICHNKTTVTSSKISQRDPSPESNKKGESPSLESRSTAVTRTSSIHQLIAPASYSP<br>IQPHSLIKHQRIPLHSPPSKVSHHQLILQQQQQIQPITLQNSTQDPPPSQHCIPLQNHGLPPAPSNAQSQHCSPIQSHPSPLTVSPNQSQSAQQSVVVSPPPPHSPSQSPT<br>IIIHPQALIQPHPLVSSALQPGPDLQQSTANQVQATAQLNLPSHLPLPASPVVHIGPVQQSALVSPGQQIVSPSHQQYSSLQSSPIPIASPPQMSTSPPAQIPPLPLQSMQSL<br>QVQPEILSQGQVLVQNALVSEEELPAAEALVQLPFQTLPPPQTVAVNLQVQPPAPVDPPVVYQVEDVCEEEMPEESDECVRMDRTPPPPTLSPAAITVGRGEDLTSEHP<br>LLEQVELPAVASVSASVIKSPSDPSHVSVPPPPLLLPAATTRSNSTSMHSSIPSIENKPPQAIVKPQILTHVIEGFVIQEGLEPFPVSRSSLLIEQPVKKRPLLDNQVINSVC<br>VQPELQNNTKHADNSSDTEMEDMIAEETLEEMDSELLKCEFCGKMGYANEFLRSKRFCTMSCAKRYNVSC**GVGSGETNGLGTGGIVGVDAMALVDRLDEAMAEEK**<br>**MQTEATPKLSESFPILGASTEVPPMSLPVQAAISAPSPLAMPLGSPLSVALPTLAPLSVVTSGAAPKSSEVNGTDR**TEPSIWTVDDVWAFIHSLPGCQDIADEFRAQE<br>IDGQALLLLKEDHLMSAMNIKLGPALKICARINSLKES | Cell experiments | 6; S6; S7 |
| **PHC3 PhL-PHC3SAMsurf** | MVSKGEELFTGVVPILVELDGDVNGHKFSVSGEGEGDATYGKLTLKLICTTGKLPVPWPTLVTTLGYGLQCFARYPDHMKQHDFFKSAMPEGYVQERTIFFKDDGNYKT<br>RAEVKFEGDTLVNRIELKGIDFKEDGNILGHKLEYNYNSHNVYITADKQKNGIKANFKIRHNIEDGGVQLADHYQQNTPIGDGPVLLPDNHYLSYQSALSKDPNEKRDHM<br>VLLEFVTAAGITLGMDELYKHRYTSLYKKAGSTMDTEPNPGTSSVSTTTSSTTTTTITTSSSRMQQPQISVYSGSDRHAVQVIQQALHRPPSSAAQYLQQMYAAQQQHLM<br>LHTAALQQQHLSSSQLQSLAAVQASLSSGRPSTSPTGSVTQQSSMSQTSINLSTSPTPAQLISRSQASSSTSGSITQQTMLLGSTSPTLTASQAQMYLRAQMLIFTPATTV<br>AAVQSDIPVVSSSSSSSCQSAATQVQNLTLRSQKLGVLSSSQNGPPKSTSQTQSLTICHNKTTVTSSKISQRDPSPESNKKGESPSLESRSTAVTRTSSIHQLIAPASYSP<br>IQPHSLIKHQRIPLHSPPSKVSHHQLILQQQQQIQPITLQNSTQDPPPSQHCIPLQNHGLPPAPSNAQSQHCSPIQSHPSPLTVSPNQSQSAQQSVVVSPPPPHSPSQSPT<br>IIIHPQALIQPHPLVSSALQPGPDLQQSTANQVQATAQLNLPSHLPLPASPVVHIGPVQQSALVSPGQQIVSPSHQQYSSLQSSPIPIASPPQMSTSPPAQIPPLPLQSMQSL<br>QVQPEILSQGQVLVQNALVSEEELPAAEALVQLPFQTLPPPQTVAVNLQVQPPAPVDPPVVYQVEDVCEEEMPEESDECVRMDRTPPPPTLSPAAITVGRGEDLTSEHP<br>LLEQVELPAVASVSASVIKSPSDPSHVSVPPPPLLLPAATTRSNSTSMHSSIPSIENKPPQAIVKPQILTHVIEGFVIQEGLEPFPVSRSSLLIEQPVKKRPLLDNQVINSVC<br>VQPELQNNTKHADNSSDTEMEDMIAEETLEEMDSELLKCEFCGKMGYANEFLRSKRFCTMSCAKRYNVSC**GVGSGETNGLGTGGIVGVDAMALVDRLDEAMAEEK**<br>**MQTEATPKLSESFPILGASTEVPPMSLPVQAAISAPSPLAMPLGSPLSVALPTLAPLSVVTSGAAPKSSEVNGTDR**TEPSIWTV**KR**VWAFIHSLPGCQDIA**KR**FRAQE<br>IDGQALLLLKEDHLMSAMNIKLGPALKICARINSLKES | Cell experiments | 6; S6; S7 |
| **Design of endogenous edited Ph variants (here WT with C-terminal extension from T2A-PuroR)** | MDRRALKFMQKRADTESDTTTPVSTTASQGISASAILAGGTLPLKDNSNIREKPLHHNYNHNNNNSSQHSHSHQQQQQQQVGGKQLERPLKCLETLAQKAGITFDEKY<br>DVASPPHPGIAQQQATSGTGPATGSGSVTPTSHRHGTPPTGRRQTHTPSTPNRPSAPSTPNTNCNSIARHTSLTLEKAQNPGQQVAATTTVPLQISPEQLQQFYASNPY<br>AIQVKQEFPTHTTSGSGTELKHATNIMEVQQQLQLQQQLSEANGGGAASAGAGGAASPANSQQSQQQQHSTAISTMSPMQLAAATGGVGGDWTQGRTVQLMQPSTSFL<br>YPQMIVSGNLLHPGGLGQQPIQVITAGKPFQGNGPQMLTTTTQNAKQMIGGQAGFAGGNYATCIPTNHNQSPQTVLFSPMNVISPQQQQNLLQSMAAAAQQQQLTQQQQQ<br>FNQQQQQQLTQQQQQLTAALAKVGVDAQGKLAQKVVQKVTTTSSAVQAATGPGSTGSTQTQQVQQVQQQQQQTTQTTQQCVQVSQSTLPVGVGGQSVQTAQLLNAGQAQ<br>QMQIPWFLQNAAGLQPFGPNQIILRNQPDGTQGMFIQQQPATQTLQTQQNQIIQCNVTQTPTKARTQLDALAPKQQQQQQQVGTTNQTQQQQLAVATAQLQQQQQQLTAAA<br>LQRPGAPVMPHNGTQVRPASSVSTQTAQNQSLLKAKMRNKQQPVRPALATLKTEIGQVAGQNKVVGHLTTVQQQQQATNLQQVVNAAGNKMVVMSTTGTPITLQNGQT<br>LHAATAAGVDKQQQQLQLFQKQQILQQQQMLQQQIAAIQMQQQQAAVQAQQQQQQQVSQQQQVNAQQQQAVAQQQQAVAQAQQQQREQQQQVAQAQAQHQQALANATQQI<br>LQVAPNQFITSHQQQQQQQLHNQLIQQQLQQQAQAQVQAQVQAQAQQQQQQREQQQNIIQQIVVQQSGATSQQTSQQQQHHQSGQLQLSSVPFSVSSSTTPAGIATSSALQ<br>AALSASGAIFQTAKPGTCSSSSPTSSVVTITNQSSTPLVTSSTVASIQQAQTQSAQVHQHQQLISATIAGGTQQQPQGPPSLTPTTNPILAMTSMMNATVGHLSTAPPVTV<br>SVTSTAVTSSPGQLVLLSTASSGGGGSIPATPTKETPSKGPTATLVPIGSPKTPVSGKDTCTTPKSSTPATVSASVEASSSTGEALSNGDASDRSSTPSKGATTPTSKQS<br>NAAVQPPSSTTPNSVSGKEEPKLATCGSLTSATSTSTTTTITNGIGVARTTASTAVSTASTTTTSSGTFITSCTSTTTTTTTSGG**MVSKGEELFTGVVPILVELDGDVNGH**<br>**KFSVSGEGEGDATYGKLTLKLICTTGKLPVPWPTLVTTLGYGLQCFARYPDHMKQHDFFKSAMPEGYVQERTIFFKDDGNYKTRAEVKFEGDTLVNRIELKGIDFKED**<br>**GNILGHKLEYNYNSHNVYITADKQKNGIKANFKIRHNIEDGGVQLADHYQQNTPIGDGPVLLPDNHYLSYQSALSKDPNEKRDHMVLLEFVTAAGITLGMDELYKGGS**I<br>SNGSKDLPKAMIKPNVLTHVIDGFIIQEANEPFPVTRQRYADKDVSDEPPKKKATMQEDIKLSGIASAPGSDMVACEQCGKMEHKAKLKRKRYCSPGCSRQAKNGIGGV<br>GSGETNGLGTGGIVGVDAMALVDRLDEAMAEEKMQTEATPKLSESFPILGASTEVPPMSLPVQAAISAPSPLAMPLGSPLSVALPTLAPLSVVTSGAAPKSSEVNGTDR<br>PPISSWSVDDVSNFIRELPGCQDYVDDFIQQEIDGQALLLLKEKHLVNAMGMKLGPALKIVAKVESIKEVPPPGEAKDPGAQSGGEGRGSLLTCGDVEENPG | Endogenous edits | 7; S7 |

**Table S7: Protein sequences from experiments and simulations – related to Methods.** N-terminal tags and leader sequences as well as C-terminal tags or extensions are underlined. Mutations such as linker swaps or SAM mutations as well as the internal Venus tag of the endogenous Ph edit are bold. For those cases, where only the representative WT sequence is shown, sequence context of mutated regions (PHC3L, SAMsurf, EH) was same as in mini-Ph variants for *in vitro* experiments.
